# Unprotected Peptide Macrocyclization and Stapling via A Fluorine-Thiol Displacement Reaction

**DOI:** 10.1101/2020.09.09.290379

**Authors:** Md Shafiqul Islam, Samuel L. Junod, Si Zhang, Zakey Yusuf Buuh, Yifu Guan, Kishan H Kaneria, Zhigang Lyu, Vincent Voelz, Weidong Yang, Rongsheng E. Wang

## Abstract

Stapled peptides serve as a powerful tool for probing protein-protein interactions, but its application has been largely impeded by the limited cellular uptake. Here we report the discovery of a facile peptide macrocyclization and stapling strategy based on a fluorine thiol displacement reaction (FTDR), which renders a class of peptide analogues with enhanced stability, affinity, and cell permeability. This new approach enabled selective modification of the orthogonal fluoroacetamide side chains in unprotected peptides, with the identified 1,3-benzenedimethanethiol linker promoting alpha helicity of a variety of peptide substrates, as corroborated by molecular dynamics simulations. The cellular uptake of these stapled peptides was universally enhanced compared to the classic ring-closing metathesis (RCM) stapled peptides. Pilot mechanism studies suggested that the uptake of FTDR-stapled peptides may involve multiple endocytosis pathways. Consistent with the improved cell permeability, the FTDR-stapled lead Axin analogues demonstrated better inhibition of cancer cell growth than the RCM-stapled analogues.

**Graphical Abstract:** 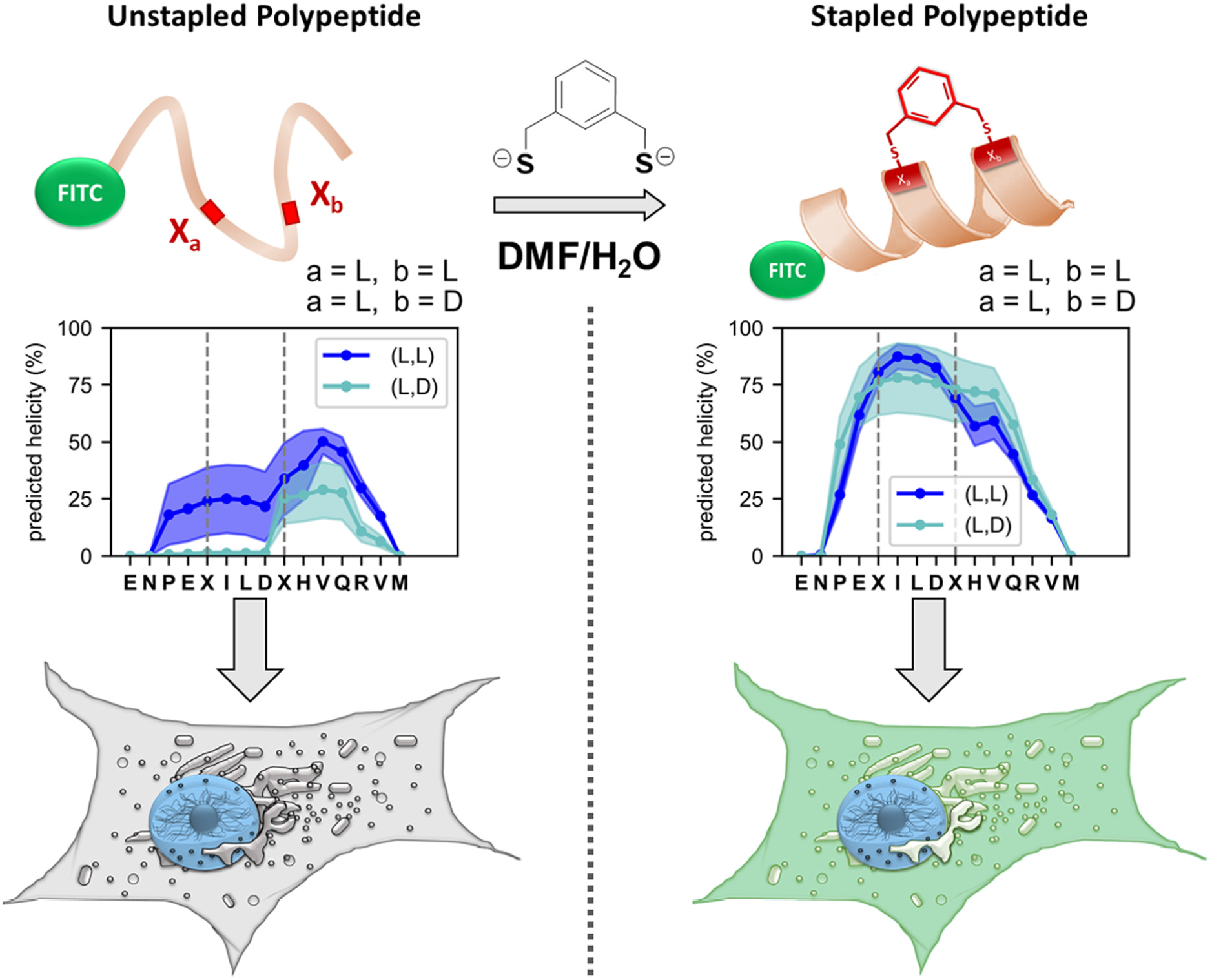

## Main

Protein-protein interactions (PPIs) regulate important molecular processes including gene replication, transcription activation, translation, and transmembrane signal transduction.^1–3^ Aberrant PPIs have been consequently implicated in the development of diseases such as cancer, infections, and neurodegenerative diseases.^3,4^ Targeting PPIs has since emerged as a promising therapeutic strategy that specifically inhibits specific molecular pathways without compromising other functions of the involved proteins.^5^ Yet this avenue is challenging due to the generally flat, shallow, and extended nature of PPI interfaces.^1,6,7^ In this regard, efficient mimicking of peptides involved in PPIs is a long-standing direction in PPI inhibitor development.^6^ Driven by entropy and the interactions with water, short peptides are mostly unstructured random coils in aqueous solutions.^8^ Given that over fifty percent of PPIs involve α-helices,^9,10^ one common approach uses chemical stapling to restore and stabilize the bioactive helical conformation of short peptides.^11^ By introducing side-chain to side-chain crosslinking of residues positioned at the same face (an *i, i+4* or *i*, *i*+7 fashion) of the α helix, the structure of peptides are locked in folded conformation due to the reduced entropy penalty.^11^

Using olefin-containing unnatural amino acids and ring-closing metathesis (RCM) mediated chemical crosslinking, Grubbs, Verdine, and Walensky et. al. have done seminal work to develop RCM-stapled peptides as a promising class of therapeutics.^12–18^ The hydrocarbon stapled peptides not only maintained the desired tertiary structures and the associated targeting specificities, but also possessed increased binding affinity and protease resistance.^1,15,18^ One hydrocarbon stapled peptide, a lead p53 mimic (ALRN-6924), has been in clinical testing to treat a diverse set of tumors including acute myeloid leukemia (AML).^2^ Yet, RCM based stapling necessitates the use of metal-based catalysts that were sometimes incompatible with other functional groups present in peptides such as a thiourea moiety.^19^ The increased hydrophobicity brought by hydrocarbon linkers may also incur issues in targeting specificity and aqueous solubility.^19^ To date, a number of other stapling strategies based on different crosslinking chemistry have been established,^7,11,20–27^ each bearing different strengths and weaknesses.^7,24,25^ For instance, many approaches required additional catalysts such as metal complexes or photoinitiators. Direct crosslinking based on bromo or iodo-alkyl/benzyl chemistry, on the other hand, could produce over-alkylated species due to their increased cross-reactivity.^20–22,25^ More importantly, the vast majority of stapled peptides still had limited cell permeability.^28^ As a result, the development of stapled peptides with designable cell permeability remains challenging and requires a delicate balance between positive charges, hydrophobicity, alpha-helicity, and staple position.^28–30^

Thus, novel strategies capable of stapling unprotected peptides in a straightforward, chemoselective, and clean manner, as well as promoting cellular uptake are highly sought.^11^ Unlike other readily reacting α-haloacetamides, a fluoroacetamide functional group has been considered biologically inert due to the poor leaving capability of fluorine.^31^ A number of probes consisting of the radiolabeled [^18^F]fluoroacetamide have been thereby developed as positron emission tomography (PET) agents for in vivo diagnostics,^32–34^ suggesting the biorthogonality of the fluoroacetamide functional group. Nevertheless, fluoroacetamide was recently incorporated into proteins as the side-chain of an unnatural amino acid, and was revealed to react with the thiol group of cysteine within protein-confined close proximity.^31^ Although driven by the proximity effect, this type of reaction may happen intermolecularly if at higher concentrations. Further, with a more nucleophilic sulfhydryl functional group, fluorine could be more efficiently displaced. Together with the biorthogonality of fluoroacetamide, this suggests that a fluorine-thiol displacement reaction (FTDR) happening intermolecularly could be highly selective and would satisfy the requirements of a “click” chemistry reaction, making it potentially attractive in the context of bioconjugations. Herein, we report the discovery of a fluorine displacement reaction using different thiol-containing linkers, which could be used for the mild functionalization of unprotected peptides. This versatile approach allowed the facile preparation of constrained macrocyclized peptides of different linker sizes, and led to the identification of stapled peptides that possessed improved target binding and cellular permeability compared to hydrocarbon stapled peptides. The peptide analogues stapled via FTDR using the 1,3-benzenedimethanethiol linker in general demonstrated approximately five-fold enhanced cellular uptake than the hydrocarbon stapled ones. The corresponding Axin analogues also showed enhanced growth inhibition of cancer cell growth as a result of increased cellular uptake, demonstrating the potential of FTDR-stapled peptides as probes for targeting intracellular compartments.

## Results

### FTDR-based coupling with a model compound/peptide

We started first by using the reported fluoroacetamide **1** as a model compound that is UV active and allows the facile monitoring of reaction progress on LC-MS.^35^ Evaluation of its reaction with common nucleophiles such as methyl hydrazine and cysteine at a mildly basic Tris (tris(hydroxymethyl)-aminomethane) buffer (Supplementary Information Figure S1) revealed that although there was no reaction between fluoroacetamide and methyl hydrazine after 12h of incubation at 37°C, approximately 40.4% of fluoroacetamide has been converted by cysteine to the fluorine-displaced adduct. This suggests that the fluoroacetamide functional group reacts specifically to thiols at high concentrations. We then attempted the fluorine thiol displacement reaction with a more nucleophilic benzyl thiol,^36,37^ and have observed a further enhanced reaction (Figure S2), rendering a relative yield of 52.7%. With this encouraging result, we further attempted the reaction on a protected amino acid building block **9** which has a natural amino acid backbone but possesses a fluoroacetamide functional group in the side chain (Figure 1a). Consistent with the previously observed FTDR results, the alpha fluoride was efficiently replaced by benzyl thiol, giving the desired product **17** at a yield of 73% after purification. We then moved on to evaluate the chemoselectivity of this FTDR as a general macrocyclization method using a seven amino acid long model peptide **18** that bears multiple unprotected functional groups (Figure 1b). Given the documented ability of the sulfur to stabilize the negative charge,^26^ 1,4-benzenedimethanethiol, a previously reported bifunctional thiol linker,^37^ was deprotonated with a stoichiometric amount of base and was then incubated with the model peptide **18** in a mixture of water/DMF. The reaction was monitored by LC-MS analysis (Figure S3), which indicated significant conversion of the starting material and the mono-substituted intermediate into the macrocyclized peptide product **19**. The observed transformation was >90% completed after 12h, and the yield was approximately 62% after HPLC purification. Given the observed conversion with benzenedimethanethiol, the FTDR based coupling appears to be chemoselective and orthogonal to functional groups in natural amino acid side chains such as carboxylic acids, amines, and phenolic alcohols.

**Figure 1.**
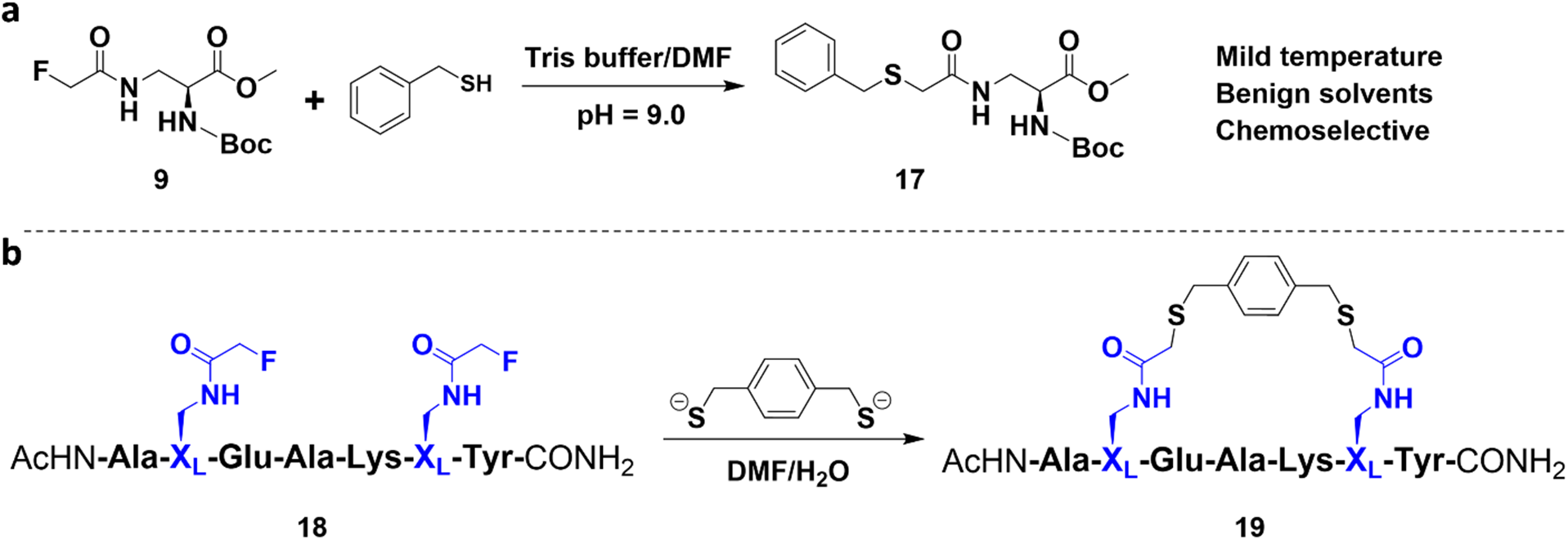
**a**, A model fluorine thiol displacement reaction (FTDR) between benzyl thiol and the α-fluoroacetamide containing amino acid building block **9**. **b**, A model macrocyclization carried out between unprotected model peptide **18** and a commercially available linker 1,4-benzenedimethanethiol based on FTDR. See SI for full characterization.

### Substrate scope with various linkers

We then decided to evaluate the FTDR-based cyclization on varied peptides of interesting chemical and biological properties. The Axin mimetic analogue first caught our attention as it was a classic peptide used for stapling,^14,23^ and its blocking of the Axin-β-catenin PPI has been shown to inhibit the Wnt signaling pathway that is crucial for the development of colorectal cancers and acute myeloid leukemia.^14,38–40^ A pair of fluoroacetamide-containing amino acids with a combination of possible chirality (X_L_ or X_D_) were incorporated into the i and i+4 positions to render the fifteen amino acid long fluoroacetamide containing Axin analogues (X_L_, X_L_ for peptide **20**, X_D_, X_D_ for **27**, X_L_, X_D_ for **34**, and X_D_, X_L_ for **38**) (Figure 2a,b). With the optimized reaction conditions, we attempted FTDR-based macrocyclization on these peptides with aliphatic dithiols of various lengths or more rigid and reactive benzenedimethanethiol linkers. As demonstrated in Figure 2, most linkers can cyclize fluoroacetamide-containing peptides in satisfactory yields (Table S1). Efficient macrocyclization was observed on both X_L_ (**20**) or X_D_ (**27**) enantiomer combination with aliphatic linkers from 5-carbon to 8-carbon long, and benzenedimethanethiol linkers at meta or para positions. The oppositely angled fluoroacetamide-containing side chains in the X_L_, X_D_ (**34**) or X_D_, X_L_ (**38**) substrates seemed to bring in more rigid conformations and thereby can be only crosslinked by 7-carbon and 8-carbon long aliphatic linkers as well as 1,3-benzenedimethanethiol as the only tolerated aromatic linker. For all the substrates being tested, macrocyclization appeared to proceed more efficiently than those in small molecule model reactions, presumably due to the complete deprotonation of thiols in advance that ensured the generation of more nucleophilic thiolate anions.^41^ Taken together, FTDR-based coupling represents an efficient macrocyclization approach in synthesizing cyclic unprotected peptides with flexible linker choices.

**Figure 2.**
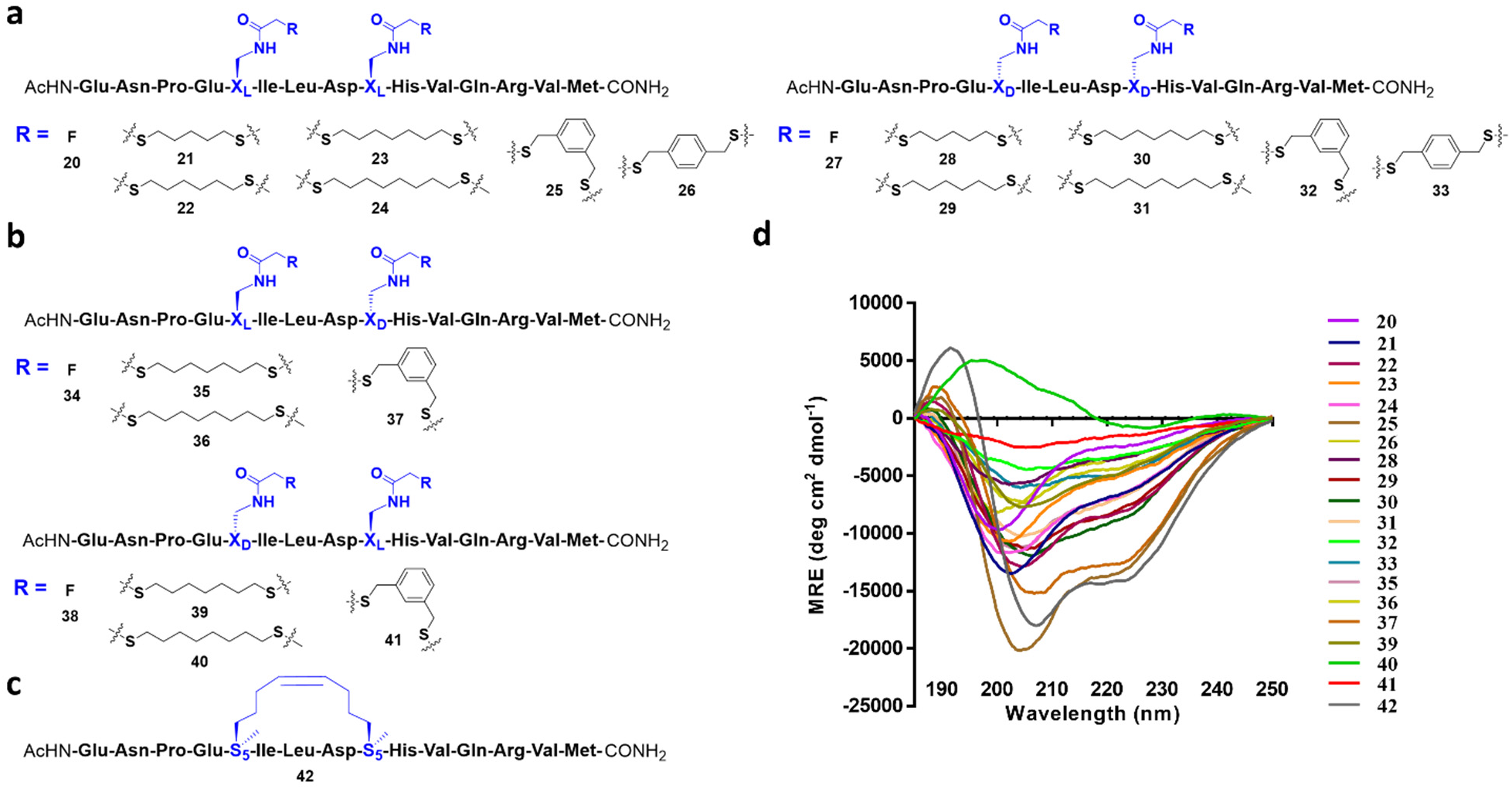
FTDR coupling between an unprotected Axin peptide analogue and various dithiol linkers. **a**, Coupling with the unprotected analogue **20** that has both L-fluoroacetamide substrates and the analogue **27** that has both D-fluoroacetamide substrates, respectively. **b**, Coupling with the unprotected analogue **34** that has L-fluoroacetamide and D-fluoroacetamide (N→C direction), and analogue **38** that has D-fluoroacetamide and L-fluoroacetamide (N→C direction), respectively. **c**, A reported Axin analogue **42** was prepared based on ring-closing metathesis. **d**, Circular dichroism (CD) spectra of all peptide analogues. See SI for full characterizations.

### Linker and chirality requirement in FTDR-based stapling

With the library of cyclized Axin peptide analogues on hand, we went on to evaluate the stapling effect of these dithiol linkers crosslinked at the *i*, *i*+4 positions. As a positive control, we prepared the RCM-stapled Axin analogue **42** (Figure 2c) that had a reported helicity of around 51%.^14^ Circular dichroism (CD) experiments were performed to measure the alpha helicity of these peptides (Figure 2d). As demonstrated in Table 1, the highest helicity was achieved with peptides **25** and **37**, with a mean value of 46% and 44%, respectively, which were close to the measured helicity of the RCM stapled control **42**. Although **25** (L,L) and **37** (L,D) possess different substrate chirality’s, both analogues were stapled well by the 1,3-benzenedimethanethiol linker, suggesting this aromatic linker could universally promote the alpha helicity of fluoroacetamide-containing peptides. In comparison, cyclized peptides with chirality combinations of D,D (**28** – **33**) or D,L (**39** – **41**) all failed to display enhanced alpha helicity to a comparable extent, even for the ones (**32** with a helicity of 13%, **41** with a helicity of 8%) stapled by the 1,3-benzenedimethanethiol linker. Consistent to the literature report,^42^ substitution with both D-configured amino acids at *i*, *i*+4 positions largely destabilized the intrinsic helix conformation. Yet, the inclusion of D-amino acids at the *i* position has been well documented, with the resulting D,L crosslinked peptide substrates usually inheriting enhanced stabilization of alpha helixes and improved binding affinity in comparison to the L,L crosslinked peptides.^20,42–45^ On the other hand, there was little observation of D-amino acid’s beneficial effects at the *i*+4 position either, as most crosslinked L,D analogues resulted in a negative effect to the peptides’ alpha helix conformation.^43,45^ Nevertheless, D-propargylglycine was once substituted at *i*+4,^27^ and compared to its L-enantiomer, Copper-mediated Huisgen 1,3-dipolar cycloaddition between L-azido norleucine and D-propargylglycine at *i*, *i*+4 positions resulted in much less distortion to the peptide backbone conformation.^27^ These suggested that for stapling-induced alpha helical stabilization the chirality requirements of substrate side chains could largely depend on the chemistry used for stapling, and specifically the chemical structures desired for both side chains and crosslinkers.

**Table 1.** Summary of the helicity of peptides stapled at *i*, *i*+4 positions.

| Peptide | Fluoro<br>substrates | Helicity<br>(%) | Peptide | Fluoro<br>substrates | Helicity<br>(%) |
| --- | --- | --- | --- | --- | --- |
| **21** | L<sub>i</sub>, i+4L | 24% | **35** | L<sub>i</sub>, i+4D | 25% |
| **22** | L<sub>i</sub>, i+4L | 29% | **36** | L<sub>i</sub>, i+4D | 14% |
| **23** | L<sub>i</sub>, i+4L | 18% | **37** | L<sub>i</sub>, i+4D | 44% |
| **24** | L<sub>i</sub>, i+4L | 25% | **39** | D<sub>i</sub>, i+4L | 19% |
| **25** | L<sub>i</sub>, i+4L | 46% | **40** | D<sub>i</sub>, i+4L | 5% |
| **26** | L<sub>i</sub>, i+4L | 17% | **41** | D<sub>i</sub>, i+4L | 8% |
| **28** | D<sub>i</sub>, i+4D | 14% | **42**<sup>*</sup> | S<sub>5</sub>, S<sub>5</sub> | 48% |
| **29** | D<sub>i</sub>, i+4D | 28% | **43** | L<sub>i</sub>, i+4L | 50% |
| **30** | D<sub>i</sub>, i+4D | 31% | **44** | L<sub>i</sub>, i+4D | 45% |
| **31** | D<sub>i</sub>, i+4D | 25% | **45** | D<sub>i</sub>, i+4L | 9% |
| **32** | D<sub>i</sub>, i+4D | 13% | **46**<sup>*</sup> | S<sub>5</sub>, S<sub>5</sub> | 52% |
| **33** | D<sub>i</sub>, i+4D | 19% |  |  |  |

\*: Control peptides stapled by ring-closing metathesis.

To explore if the L,D versus D,L substrate preference for FTDR-based stapling is generally applicable to different peptide sequences, we turned our attention to another model peptide, which binds to the C-terminal region of an HIV-1 capsid assembly polyprotein (HIV C-CA) that is key to viral assembly and core condensation.^46^ The 12-mer long peptide was previously demonstrated to have efficiently stabilized alpha-helical conformation once RCM-based stapling was performed at the indicated *i* and *i*+4 positions (**46**, Figure 3).^26,47^ Using the optimized linker (1,3-benzenedimethanethiol), we performed FTDR-based stapling on the same sequence positions, and obtained analogues **43**-**45** that have fluoroacetamide substurates of different chirality combinations (Figure 3a, Table S2). FITC labeling was applied to the N-terminal during solid-phase synthesis in order to facilitate follow-up biological characterizations. Notably, stapling of the unprotected FITC-labelled peptides **43**-**45** proceeded smoothly despite the presence of FITC’s thiourea moiety. As shown in CD spectra (Figure 3b) and Table 1, stapling with L,L or L,D substrates generated peptides **43** and **44** that possessed similar helicity to the control peptide stapled by RCM. On the contrary, the crosslinked D,L substrate-containing analogue **45** only displayed a minimal level of helicity.

**Figure 3.**
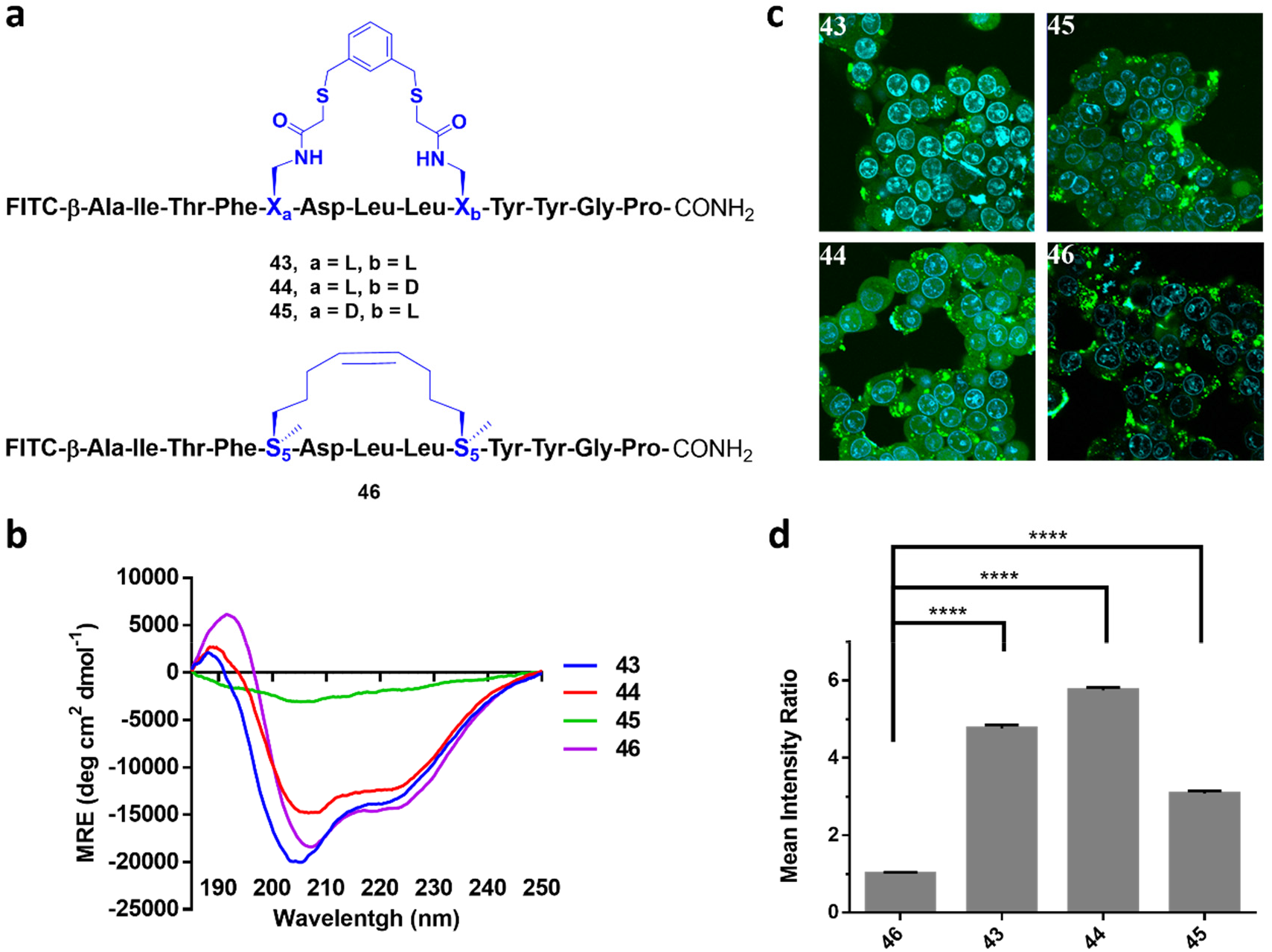
**a**, FTDR coupling between unprotected HIV C-CA binding peptide analogues and 1,3-benzenedimethanethiol. The reported HIV C-CA binding peptide **46** (stapled by RCM) was prepared as well. See SI for full characterization. **b**, CD spectra of the crosslinked peptide analogues. **c**, Fluorescent confocal microscopy images of the HEK293T cells treated with peptides **43-46**. Blue: nucleus stained by Hoechst 33342; Green: FITC-labelled peptides. **d**, Quantification analysis of the cell penetration of peptides **43-46.** The intracellular intensity of peptide **46** was normalized as 1. “****” represents p < 0.0001.

Taken together, FTDR-based stapling works most efficiently with the rigid meta-benezenemethane dithiol linker, and generally prefers the L,L or L,D fluoroacetamide substrates on different peptide targets, with the D,L substrates least tolerated. To gain insight to the molecular mechanisms driving this preference, we performed extensive molecular dynamics simulations for stapled Axin peptides **21-41** and HIV C-CA binding peptides **43-45**, generating an aggregate of 1.8 ms of trajectory data (Table S3) on the Folding@home distributed computing platform.^48–50^ Markov State Models^51,52^ of the conformational dynamics (see Methods) showed that peptide stapling stabilizes helical conformations (Figure 4a,c), and slows folding by an order of magnitude to the ~10 μs time scale (Figure 4b). Predicted helicity profiles for stapled and unstapled Axin peptides (Figures 4c, S7) and stapled HIV C-CA binding peptides (Figure 4d) also showed that crosslinked D,L substrate-containing peptide analogues disrupt helicity at the D-amino acid position more than those crosslinked L,D substrate-containing peptides. This is perhaps due to subtle differences in linker strain, combined with a differential helix nucleation propensities in the N-vs-C-terminal direction for stapled peptides.^53^ In summary, predicted helicities from simulations compare well with experimental values from CD spectroscopy (Figures 4e, S8-S9)

**Figure 4.**
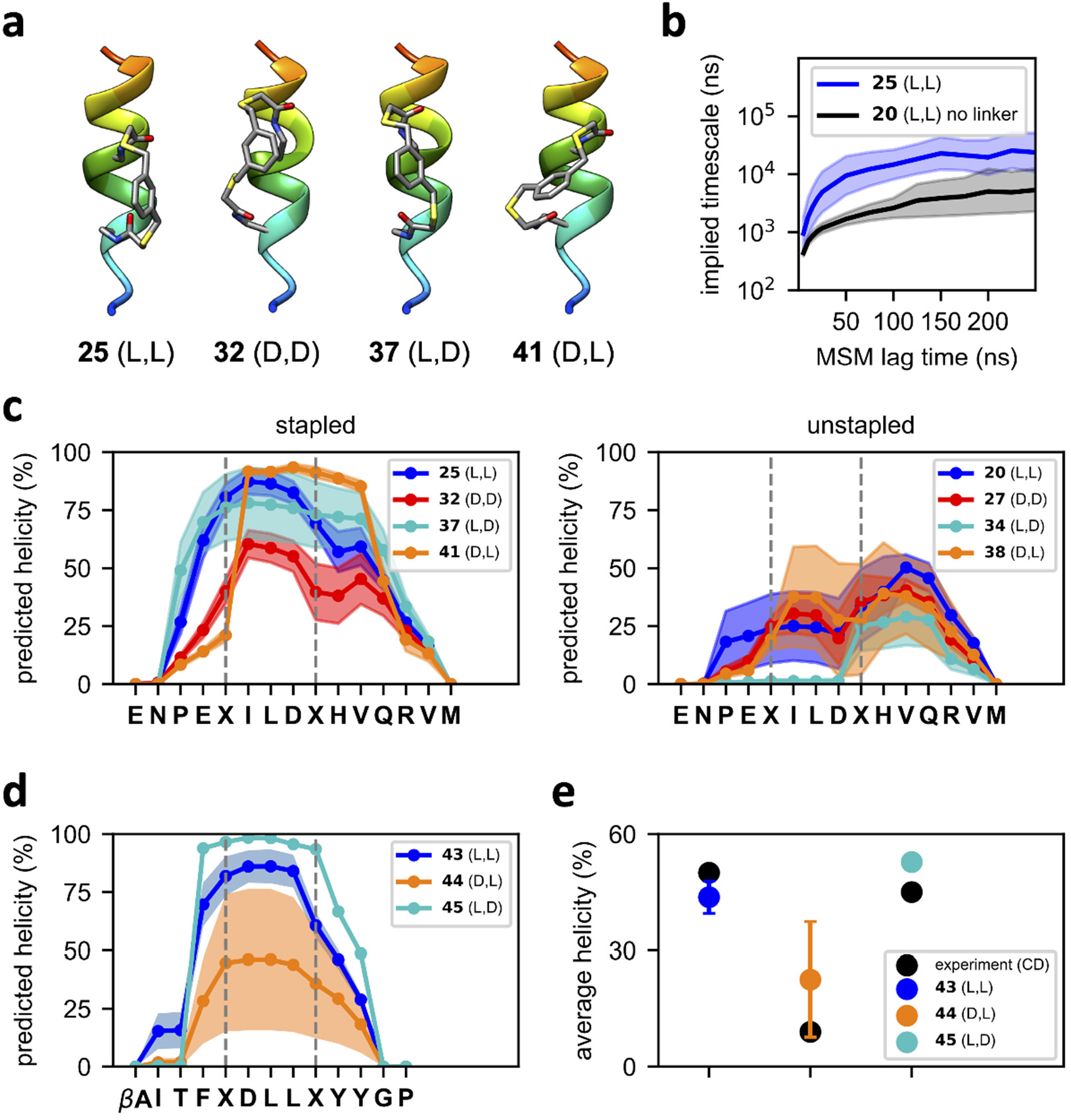
**a**, Energy-minimized structures of 1,3-benzenedimethanethiol crosslinked Axin peptides **25**, **32**, **37** and **41**. **b**, The slowest implied timescale of peptides **25** and **20** (no linker) shown as a function of MSM lag time. **c**, Predicted per-residue helicity profiles of stapled and unstapled Axin peptides. Uncertainties were estimated from a bootstrap procedure where five MSMs were constructed by sampling the input trajectory data with replacement. **d**, Predicted per-residue helicity profiles of HIV C-CA binding peptides **43**, **44** and **45**. **e**, Comparison of experimental (CD) and predicted average helicities of peptides **43**, **44** and **45.**

### Cell permeability of FTDR-stapled peptide analogues

Cellular uptake of peptides has been revealed to be a complicated process driven by hydrophobicity, positive charge, and alpha-helicity, etc.^28^ The RCM-based stapling was previously reported to endow enhanced cellular permeability to HIV C-CA binding peptide **46** due to the stabilized secondary structure and the increased hydrophobicity.^47^ Particularly for stapled peptides, the staple type also serves as one of the deciding factors for cell penetration.^29,30^ Thus, we wondered if peptides stapled by FTDR can render at least comparable cell permeability. We first investigated the dose-dependent cytotoxicity of analogues (**43**-**46**) after incubation with HEK293T cells for 12h, and did not observe any significant effects on viability with up to 15 μM of peptides (Figure S10). Towards this end, all the FITC-labelled HIV C-CA binding analogues (**43**-**46**) at 10 μM concentration were incubated with HEK293T cells for 4h and were subsequently analyzed by confocal fluorescence microscopy (Figure 3c). Significant cellular uptake of the FTDR-stapled peptides **43** and **44** were observed, with peptides not only spreading in the cytosol but also existing in the endosomes showing punctated greenish fluorescence. In comparison, much less fluorescence was observed from cells treated with the RCM stapled control **46**. A similar trend has been previously reported with HIV C-CA binding peptides stapled by perfluoroarylation of cysteines,^26^ suggesting that the aromatic part in the linkers may enhance the cellular uptake. Additionally, the weaker uptake of the D,L-stapled peptide (**45**) also corroborated the previously reported positive correlation between helicity and cell permeability.^44^

Next, we asked if FTDR-based stapling on other peptide sequences can lead to enhanced cell penetration as well. To observe the broader applications of FTDR-based stapling on other peptides and cell lines, we chose another Axin derived peptide analog (**51**, Figure 5a), which displayed single-digit nM *Kd* after *i*, *i*+4 stapling by RCM, but showed limited cell permeability to DLD-1 cells.^14^ Previously, other amino acids in this sequence had to be mutated into arginine in order to increase the overall positive charges. The resulting lead analogue had improved cellular uptake at the expense of losing 6-7 folds of affinity towards the protein target β-catenin.^14^ We thereby synthesized the FITC labelled fluoroacetamide containing L,L substrate (**47**) and L,D substrate (**49**), and decided to focus on the DLD-1 cell line as reported in order to facilitate direct comparison. FTDR-based stapling with 1,3-benzenedimethanethiol proceeded smoothly, and the resulting analogues (**48**, **50**, Figure 5a, Table S2) did not affect the viability of DLD-1 cells at 10 μM concentration after 12h incubation (Figure S10). With confidence that there is no cytotoxicity, we treated DLD-1 cells with each of these peptides, along with the unstapled or RCM-stapled control peptide in parallel. As shown in Figure 5b and Figure S11, significant cellular uptake was observed from cells incubated with 10 μM FTDR stapled **48** or **50**, while there was much weaker fluorescence in other control treatment groups including the cells incubated with **51**. The uptake pattern of **48** and **50** were similar to those observed earlier for HIV C-CA binding peptides **43** and **44**. A quantitative analysis was achieved by applying the lognormal fitting to a histogram of the individual cell mean intensity (Figures S12 – S13), and the resulting mean cellular fluorescence revealed that the FTDR stapled L,L HIV C-CA binding peptide **43** had 4.76 ± 0.09 fold of mean intensity compared to that of the RCM control peptide intracellularly, and the stapled L,D mimetic **44** had 5.75 ± 0.07 fold mean intensity (Figure 3d, Table S4). Quite consistently, the L,L Axin derivative **48** showed a 4.86 ± 0.15 fold of mean intensity compared to the RCM control one, while the L,D Axin mimetic **50** showed a 5.05 ± 0.14 fold increase (Figure 5c, Table S5). In both cases, stapled L,D analogues demonstrated slightly stronger cellular uptake than the stapled L,L analogues. Moreover, the diminished uptake from unstapled peptides (e.g. 0.36 ± 0.02 fold for **47**, 0.33 ± 0.01 fold for **49**) further suggests the FTDR stapling is an essential requirement for cell permeability.

**Figure 5.**
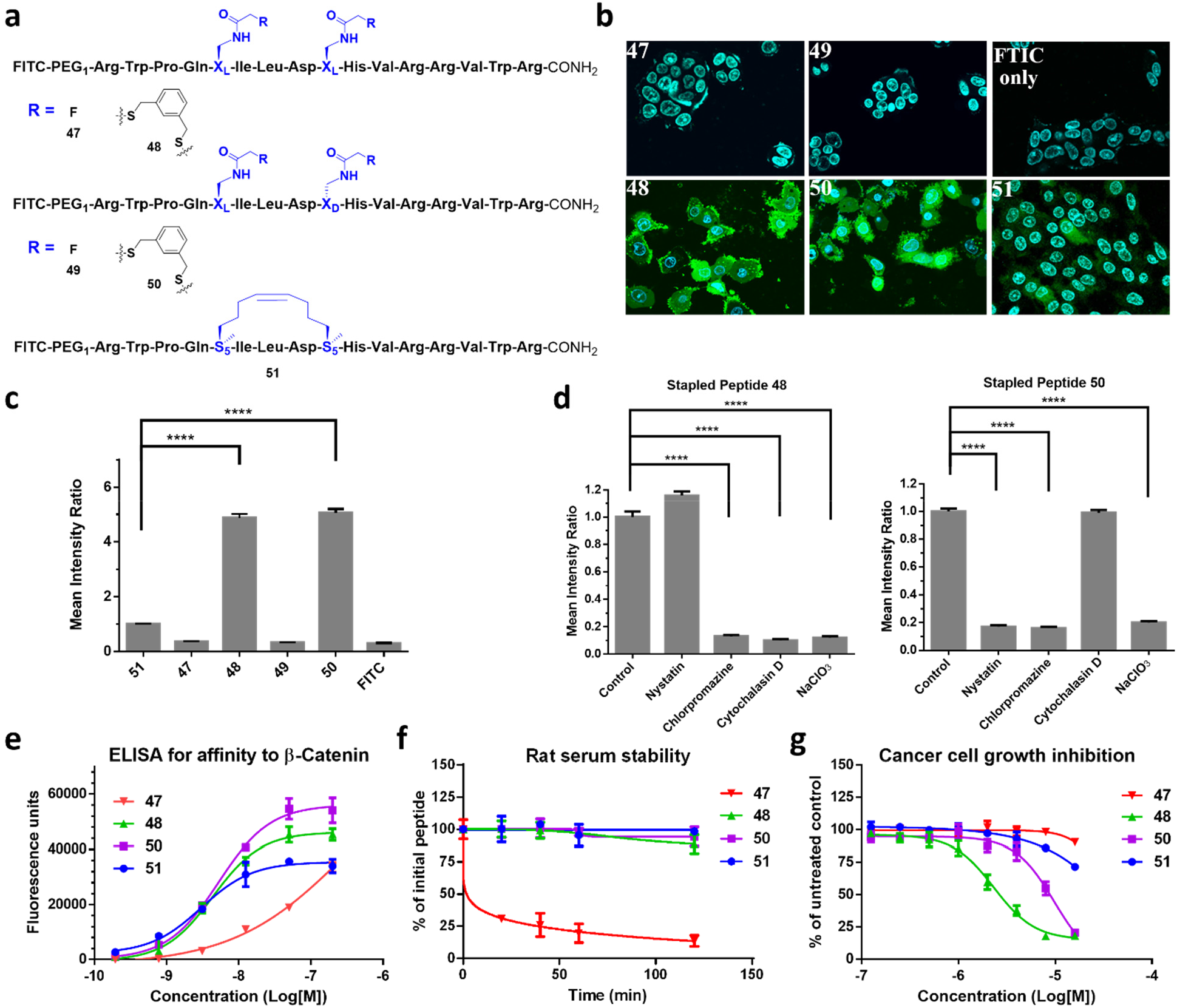
**a**, A FTDR coupling between unprotected Axin-derived peptide analogues (**47**, **49**) and 1,3-benzenedimethanethiol. The Axin analogue **51** (stapled by RCM and used for cell penetration studies) was prepared as well. See SI for full characterization. **b**, Fluorescent confocal microscopy images of the DLD-1 cells treated with peptides **47-51** and FITC only as a negative control. Blue: nucleus stained by Hoechst 33342; Green: FITC. **c**, Quantification analysis of the cell penetration of peptides **47-51.** The intracellular intensity of peptide **51** was normalized as 1. “****” represents p < 0.0001. **d**, Quantification analysis of the cell penetration of peptide **48** or **50** after the cells had been treated with blockers of endocytic pathways. The intracellular intensity of peptides in cells untreated with any blocker was normalized as 1. See SI for fluorescent microscopy images. “****” represents p < 0.0001. **e**, the binding affinity of unstapled (**47**) and stapled Axin analogues (**48, 50, 51**) to the target β-catenin protein. Peptides **48** and **50** were stapled via FTDR, while peptide **51** was stapled by ring-closing metathesis (RCM). **f**, the stability of the aforementioned peptide analogues in 100% rat serum. **g,** the inhibition of these peptide analogues on DLD-1 cancer cell growth over 5 days.

Despite intensive studies, the uptake mechanisms for cell-penetrating peptides remain ambiguous and largely varied due to their complicated nature.^54–56^ Small molecule inhibitors specific for each endocytic pathway have been routinely utilized to interrogate the cellular uptake mechanisms of many transporters.^57–60^ For example, nystatin as a sterol-binding agent was used to selectively block caveolin-dependent endocytosis,^60^ while chlorpromazine can selectively inhibit clathrin-mediated endocytosis.^58,60^ Cytochalasin D would specifically induce depolymerization of actin and ceased the subsequent apical endocytosis.^59^ Sodium chlorate, on the other hand, aborted the decoration of cell membranes with sulfated proteoglycans, affecting certain other uptake pathways.^57^ With those, RCM-stapled peptides were recently revealed to penetrate cells via clathrin- and caveolin-independent pathways that were partially mediated by cell surface proteoglycans.^29^ To understand the mechanisms behind the enhanced cellular uptake of peptides stapled by FTDR, we performed similar experimental investigations using the lead Axin derivatives **48** and **50**. The cell penetration experiments were repeated under conditions that blocked a different endocytotic pathway each round (Figures S14 – S15). Cell viabilities were measured immediately after the imaging experiments to ensure that the observed results were due to the active cellular uptake (Figure S16). As quantitatively summarized in Figure 5d and Tables S6-S7, both peptides had their uptake partially blocked by more than one pathway-specific inhibitors. Like RCM-stapled peptides, both FTDR-stapled peptides internalized partially through sulfated proteoglycans, as indicated by the reduced uptake in cells treated by sodium chlorate. Yet unlike hydrocarbon-stapled peptides, the L,L stapled analogue **48** also partially penetrated cells via endocytosis depending on clathrin (inhibited by chlorpromazine) and actin polymerization (inhibited by cytochalasin D). The L,D stapled **50** appeared to enter cells additionally through clathrin- and caveolin-dependent (blocked by nystatin) endocytosis. Interestingly, the chirality difference in the i+4 position between the stapled peptide **48** and **50** seemed to result in different peptide backbone conformations, thereby affecting their intake through distinguished endocytosis pathway. Taken together, our data suggested that FTDR-stapled peptides may penetrate cells through multiple endocytotic pathways in a distinct pattern compared to the RCM-stapled peptides, which could account for their enhanced cellular uptake than those observed for the RCM-stapled controls.

### Activity of FTDR-stapled peptide analogues

In addition to cell permeability, we wondered whether the structural features brought by the 1,3-benzenedimethanethiol crosslinker translates to other functional relevance. Thus, we performed ELISA assay to quantify the binding affinity of the lead Axin derivatives towards the target protein β-catenin (Figure 5e). The EC_50_ was 4.36 ± 1.75 nM for stapled analogue **48**, and 5.27 ± 2.29 nM for analogue **50**, which were similar to the EC_50_ of RCM-stapled **51** (3.03 ± 1.8 nM) and were at least 100-fold more potent than unstapled peptide **47**. Notably, a direct comparison of the fluorescence signals for all the peptides at 50 μM or 200 μM doses seemed to pinpoint the potentially better binding of FTDR stapled peptides than the RCM control **51**. Given that staples and the peptide conformational changes after stapling usually render the structures less prone to protease-mediated cleavage,^15,18,26^ we also tested the serum stability of these lead peptides in 100% rat serum. As shown in Figure 5f, all the stapled peptides remained mostly intact while unstapled control **47** was rapidly degraded, with only 30.9% left after 20 min of incubation. In order to see if their improved cellular uptake can translate to enhanced cellular activity, we finally examined these analogues’ inhibition of the growth of a Wnt-driven colorectal cancer cell line^14^ DLD-1 over a 5-day period (Figure 5g). FTDR stapled **48** and **50** potently impeded the cell growth with EC_50_s of 2.3 ± 0.4 μM, and 9.8 ± 2.0 μM, respectively. On the contrary, neither the unstapled control **47** nor the RCM-stapled analogue **51** displayed a significant growth inhibition until they were administered at the 16 μM concentration. Together, unprotected peptides stapled by FTDR appeared to recapitulate the advantages in biological functions seen in the classic RCM-stapled peptides, but also possessed enhanced cell permeability and the correlated growth inhibition of the targeted cancer cells.

## Discussion

In summary, we have demonstrated a new, mild, and clean synthetic strategy to cyclize and/or staple unprotected peptides. The developed fluorine-thiol displacement reaction (FTDR) approach operates at mild temperature in aqueous solutions and offers excellent chemoselectivity and functional group tolerance. We first exemplified its application as a general macrocyclization platform that can be compatible with a variety of linkers. Then we demonstrated its use in stapling peptides at *i*, *i*+4 positions, and further showed that the lead stapled peptides retained the structure features and biological properties reported in literature. The identification of the 1,3-benzenedimethanethiol as the optimal linker for stapling suggests that certain aromatic rigidity in the crosslinker region is required to maintain the alpha-helical conformation of peptide substrates stapled by FTDR. Further, both the experimental results and the molecular dynamics simulation consistently pinpointed the preference of L,L and L,D substrate chirality for the folding (alpha helicity) of the FTDR-stapled peptides, implicating the distinct helix nucleation propensities in the N-vs-C-terminal direction for this class of stapled peptides.

In terms of biological functions, the enhanced cellular uptake of *i*, *i*+4 stapled peptides and the associated distinct penetration mechanism suggest this FTDR-based stapling approach may expand the toolbox of chemical transformations to generate a new class of probes or therapeutic leads for intracellular targets. To our best knowledge, this is also the first reported effort to elucidate the cellular uptake mechanism of peptides stapled by strategies other than RCM. Our findings confirmed the previous observations that there could be more than one uptake mechanisms existing for stapled peptides.^29^ Further, they were consistent to the previous discovery that internalization of stapled peptides mainly correlated with the staple type.^29^ Accordingly, our approach in many aspects is complementary to the RCM-mediated peptide stapling. Current efforts are focused on optimizing the FTDR reaction conditions to further improve its efficiency, and expanding the substrate scope towards *i*, *i*+7 positions. Application of the FTDR-based stapling to probe protein-protein interactions related to the key signaling events in prostate cancer and neurodegenerative diseases are also under active investigation in our research group.

## Methods

**Chemical synthesis procedures, supporting tables**, and **supporting figures** are available as Supplementary Information.

### Peptides synthesis

Peptides were synthesized on rink amide resins following traditional Fmoc-based solid-phase chemistry. Most >10 mer peptides were synthesized on an automatic peptide synthesizer at the Macromolecular core facility of the Pennsylvania State University. Generally, approximately four equivalents of Fmoc-protected amino acid building blocks, four equivalents of HATU and eight equivalents of DIPEA were added to the resins resuspended in DMF for every round of coupling. The N-terminal of peptides were capped with acetic anhydride or FITC in DMF using DIPEA as the base. After capping, peptides were cleaved by incubating the beads with reagent H (trifluoroacetic acid (81% w/w), phenol (5% w/w), thioanisole (5% w/w), 1,2-ethanedithiol (2.5% w/w), dimethylsulfide (2% w/w), ammonium iodide (1.5% w/w), and water (3% w/w))^61^ at room temperature for 3h. The supernatant was collected and added diethyl ether to precipitate out the crude product.

### General procedures for peptide stapling

For the controls that require ring closing metathesis (RCM), RCM-based stapling was performed with protected peptides on resins following the reported procedures.^14^ The resins were mixed with approximately 0.5 equivalent of Grubbs I catalyst (5 mM) in dichloromethane and were incubated for 2h at room temperature before the solvents were drained off. The coupling process was repeated three times, followed by subsequent N-terminal deprotection and capping. The crude products were cleaved from resins as mentioned before, and precipitated out by 15-fold ice-cold diethyl ether. The crude mixture was re-dissolved in a water and methanol mix, and purified by reverse phase semi-preparative HPLC that operated at a flow rate of 10 mL/min, used water/0.1% TFA as solvent A and acetonitrile/0.1% TFA as solvent B.

For stapling based on the fluorine thiol displacement reaction (FTDR), 126 μL of the dithiol-containing linker (250 mM in DMF) was premixed with 210 μL of sodium hydroxide solution (250 mM) and 189 μL of DMF at room temperature for 40 min to completely deprotonate dithiols. After this, 105 μL of peptide solution (50 mM in water or DMF) was added to start the stapling based on FTDR. The reaction mixture was incubated at 37°C for around 12h till the reaction was complete as determined by LC-MS, and was then quenched with water and acetic acid. Subsequently, the solution was extracted by ethyl acetate three times to remove excess linkers. The remaining aqueous phase was lyophilized, resuspended in methanol, and added diethyl ether to precipitate out the stapled peptides. The crude product was further purified on HPLC as mentioned before. The yields for all the stapled peptides (Tables S1, S2) were determined after recovery from HPLC purification.

### Circular dichroism

Axin- and p53-derived peptides were dissolved in water, and HIV-targeting peptides were dissolved in 25% acetonitrile/water to ensure 100% solubility. The final concentration of the peptide samples was 70 μM. Circular dichroism (CD) measurements were performed on a Jasco model J-815 spectropolarimeter with a 1 mm Jasco quartz cell over the wavelength range of 180 – 250 nm. The scans were carried out at 0.2 nm resolution with 4 s average time at 25 °C. Data from three scans were averaged, base line corrected, and normalized to the mean residue ellipticity (MRE) following the equation: [θ]_λ_ = [θ]_obs_/(10 x l x C x n). [θ]_λ_ is MRE in deg x cm^2^ x dmol^-1^; [θ]_obs_ is the measured ellipticity; C is the concentration of peptides in M; l is the optical pathlength in cm; and n is the number of residues in peptides. The % alpha helicity was calculated from the MRE values at 222 nm, using the equation % helicity = ([θ]_222_ - [θ]_0_)/([θ]_max_ - [θ]_0_) based on the previously reported method.^62^ [θ]_222_ is the MRE value at 222 nm; [θ]_max_ is the maximum theoretical MRE value for a helix of n residues; and [θ]_0_ is the MRE value of the peptide in random coil conformation that usually equals to (2220-53T).^62^

### Molecular dynamics simulation

Molecular dynamics (MD) simulations of stapled/unstapled Axin peptides and HIV peptides were performed using the OpenMM 7.0 simulation package^63^ on the Folding@home distributed computing platform.^64^ The AMBER ff14SB force field^65^ was used for the peptide residues, while non-natural residues and linkers used GAFF^66^ with partial charges from AM1-BCC,^67^ parameterized using the *antechamber* package of AmberTools17.^68^ Axin peptide simulations were initiated from helical conformations taken from crystal structure PDB:1QZ7. These structures were solvated in a ~ (55 Å)^3^ cubic periodic box with TIP3P water molecules and Na^+^ and Cl^-^ counterions at 100 mM to neutralize charge, for a total of ~16K atoms. HIV peptide simulations were initiated from helical conformations taken from the crystal structure PDB:3V3B. These structures were solvated in a ~ (45 Å)^3^ cubic periodic box with Na^+^ and Cl^-^ counterions, for a simulation size of around 9K atoms.

Trajectory production runs were performed using stochastic (Langevin) integration at 300 K with a 2-fs time step. Covalent hydrogen bond lengths were constrained using the LINCS algorithm. PME electrostatics were used with a nonbonded cutoff of 9 Å. The NVT ensemble was enforced using a Berendsen thermostat. About 50 trajectories were generated for each peptide design, reaching an average trajectory length of ~1.0 μs. In total, 1.8 milliseconds of aggregate simulation data were generated for all designs, with about ~70 μs of trajectory data per design (Table S3). Trajectory snapshots for the peptide coordinates were saved for every 500 ps. For all analysis described below, the first 100 ns was discarded from each trajectory to help remove systematic bias.

### Markov state model construction

To describe the conformational dynamics of each peptide, Markov state models (MSMs) were constructed from the trajectory data using the MSMBuilder 3.8.0 software package.^69^ This involved the following steps: (1) performing dimensionality reduction using time-structure-based independent component analysis (tICA),^70,71^ (2) conformational clustering in the reduced space to define and assign the trajectory data to discrete metastable states, and (3) estimating the transition rates between metastable states from the observed transitions between states.

The tICA method is a popular approach to project protein coordinates to a low-dimensional subspace representing the degrees of freedom along which the slowest motions occur. In tICA, structural features *f_i_* are computed for each trajectory frame, and the time-lagged correlation matrix **C**(Δ*t*) of elements *C_ij_* = <*f_i_*(*t*) *f_j_*(*t*+Δ*t*)>*t* of all pairs of features *i* and *j* are computed, where *t* is time, and Δ*t* is the tICA lag time. The tICA components (tICs) are linear combinations of features that capture the greatest time-lagged variance, which can be found by maximizing the objective function <α |**C**(Δ*t*)| α > subject to the constraint that each component has unit variance (i.e. <α|Σ |α_*i*_ > = 1). As features, we computed all pairwise distances between Cα and Cβ atoms (435 and 300 distance pairs for Axin and HIV C-CA binding peptides, respectively) using the MDTraj package.^72^ A tICA lag time of Δ*t* = 5 ns was chosen. After projecting trajectory data to the four largest tICs, conformational clustering was performed using the *k*-centers algorithm to define 50 discrete states for the construction of Markov State Models (MSMs). The GMRQ cross-validation algorithm was used to determine the optimal number of states (Figure S4).

MSM transition matrices **T**^(τ)^ were constructed at lag times τ ranging from 1 to 300 ns, using a maximum likelihood estimator. Transition matrix elements *T_ij_*^(*τ*)^ contain the probability of transitioning from state *i* to state *j* in time τ. MSM implied timescales are computed as *t_i_* = τ/ln λ_*i*_ where λ_*i*_ are the eigenvalues of the MSM transition matrix. The implied timescales plateau with increasing lag time, indicating that dyamics is approximately Markovian (i.e. memory-less) beyond a lag time of 100 ns (Figure S5). The slowest implied timescale can be interpreted as the folding/unfolding relaxation time of the peptide, which generally ooccurs along the principal tICA component, tIC_1_. Example projections of trajectory data to the first two tICs are shown in Figure S6.

### Analysis of secondary structure

The secondary structure content of simulation snapshots was calculated using the DSSP algorithm implemented in MDTraj.^6^ We computed the ensemble average of helicity <*h*> from the trajectory data for each design, using the equilibrium populations *⊓i* of each microstate *i* predicted by the MSM, as <*h*> = Σ_*i*_ π_*i*_ *h_i_*, where *h_i_* is the average helicity of snapshots belonging to state *i*.

### Cell viability assay

Wild type DLD-1 cells were cultured in complete growth media (RPMI, 10% FBS and 1% penicillin/streptomycin) at 37°C / 5% CO_2_. After harvesting, cells were placed onto 96-well flat bottom white plates at 10e4 cells/well, and were incubated overnight. The next morning, serially diluted axin analogues (final concentration 0 μM, 5 μM, 10 μM, and 15 μM) were added to the cell culture media and incubated for 12 h. The cells were then treated with CellTiter-Glo^®^ reagent that had been prepared following the manufacturer’s protocol at 100 μL/well. The chemiluminescent signals from the control sample (treated with 0 μM peptides) were used as the 100% control. For HIV-1 peptide analogues, the samples were incubated with HEK293T cells instead, and the rest of the procedure was the same as mentioned before. For DLD-1 cells, after confocal imaging for the penetration pathway studies, viability measurements were carried out similarly via directly mixing with the CellTiter-Glo reagent. Cells treated with vehicle groups were also analyzed and counted as the 100% control.

### Confocal imaging

All images were captured on an Olympus FV3000 confocal laser scanning microscope with a NA 1.05 30X silicone-immersion objective (UPLSAPO 30XS). Hoechst 33342 dye and FITC-labeled peptides were excited with 405 nm and 488 nm lasers (Coherent OBIS), respectively. Images were analyzed with NIH ImageJ software. Using the “analyze particles” function, we measured the area and intensity of individual cells (Fig. S12A).^73^ Particles in the extracellular matrix that the “analyze particle” function selected were manually deselected and removed from the data set (Fig. S12B). Cells with cytoplasmic membrane aggregation of the FITC-labeled peptide had the cytoplasmic membrane excluded from the mean intensity measurement (Figure S12C). From a normal distribution curve applied to a histogram of the individual cell mean intensity, we found some outliers in the data collected (Fig. S13A). However, from the distribution of data we found the best fit curve to be lognormal. Outliers were then removed using a Grubbs test (Fig. S13B).^74^ We determined the optimal bin size for the histogram using the Freedman-Draconis rule.^75^

### Cell penetration assay

DLD-1 cells were seeded on 35 mm optical dishes at 3e5 per well and were incubated overnight at 37°C /5% CO_2_. FITC-labelled Axin analogues were added and incubated for 12 h at 10 μM final concentration.^14^ To visualize the nucleus, the cells were incubated with Hoechst 33342 (1 μg/mL, PBS, ThermoFisher) for 10 min. Cells were imaged in buffer mimicking the physiological conditions in the cytoplasm (20 mM HEPES, 110 mM KOAc, 5 mM NaOAc, 2 mM MgOAc, 1 mM EGTA, pH 7.3). FITC labelled HIV-1 C-CA peptides were imaged similary, except that the incubation was with HEK293T cells for 4h.^26^

### Cell penetration pathway study

The plated DLD-1 cells were incubated for 1 h with 25 μg/mL nystatin for blocking caveolin-mediated endocytosis, 5 μg/mL chlorpromazine for blocking clathrin-dependent endocytosis, 10 μg/mL cytochalasin D for inhibiting actin polymerization, and 80 mM NaClO_3_ for disrupting proteglycan synthesis according to the conditions used by literature.^29^ After this, the FITC-labelled peptide analogues were added at a final concenteration of 10 μM, and the mixture was incubated at 37°C/5% CO_2_ for 4 h. The cells were washed with PBS and incubated with Hoechst 33342 (1 μg/mL, PBS) for 10 min. After another round of washing, cells were imaged in the buffer as aforementioned. Subsequent CellTiter-Glo viability measurements were also performed right after confocal imaging.

### ELISA assay for binding affinity

The beta-catenin protein (Abcam, ab63175) (1 μg/mL, 50 μL/well) was coated onto a 96-well flat bottom black plate (Nunc, MaxiSorp) at room temperature for 2 h. The wells were washed with 0.05% tween containing PBS buffer, and then blocked with 1% BSA containing PBS buffer at room temperature for 2 h. The FITC labelled Axin peptide derivatives (including the unstapled control) were each serially diluted (200 μM, 50 μM, 12.5 μM, 3.125 μM, 0.781 μM, 0.195 μM, and 0) in 100 μL of blocking solution (1% BSA, PBS buffer), added to the wells, and incubated at room temperature for another 2 h. The wells were then washed three times with PBS buffer (0.05% tween) and treated with HRP-conjugated anti-FITC antibody (100 μL/well, Abcam, ab196968) for 1h at room temperature. After this final incubation, the wells were washed 5 times with 300 μL of PBS buffer (0.05% tween), and then treated with the QuantaBlu fluorogenic peroxidase substrates that had been prepared following the manufacturer’s protocol (100 μL/well). The fluorescence signals were recorded by the H1 synergy plate reader (Biotek) at the excitation wavelength of 325 nm and the emission wavelength of 420 nm.

### Serum stability assay

Based on the ELISA-based assay of EC_50_’s (~ 4 nM) for all the stapled peptides’ binding with beta-catenin, FITC-labelled peptide samples (L_i_, _i+4_L, L_i_, _i+4_D or the RCM control) were dissolved in 100% rat serum (Sigma Aldrich) at an initial concentration of 4 μM. Given the EC_50_ value (~ 300 nM) of the unstapled peptide, a concentration of 300 μM was initially used for incubation with rat serum. Each sample mixture was then equally aliquoted into 15 tubes, and were incubated at 37°C. At every specified time point (0, 20, 40, 60 and 120 min), 3 of the tubes were collected, and immediately diluted 1/1000 with PBS buffer (1% BSA), followed by flash freezing in liquid nitrogen and short-term storage at −80°C. On the day of the ELISA assay, all the samples were slowly thawed on ice, and the remaining percentage of active peptides were determined by the ELISA assay following the procedure described above. The signals from the peptide sample mixture at 0 min were used as the 100% control.

### Cell growth inhibition assay

DLD-1 cells were cultured and maintained as mentioned before. Right before the assay, cells were resuspended in fresh RPMI media supplemented with 10% FBS, 100 IU/mL penicillin, and 100 μg/mL streptomycin, and plated onto 96-well whilte flat-bottom plates at 1000 cells per well (90 μL media/well). After overnight incubation, peptides samples were serially diluted as 10x stock in PBS buffer (10% DMSO), and were added to the plated cells at 10 μL stock solution/well in triplicate to make a final treatment concentration of 0 μM, 0.0625 μM, 0.125 μM, 0.25 μM, 0.5 μM, 1 μM, 2 μM, 4 μM, 8 μM, 16 μM. The wells on the edges of plates were filled with PBS buffer to avoid unwanted evaporation of samples. The samples on the plates were incubated at 37 C, 5% CO_2_ for 5 days, and the cell growth was evaluated by CellTiter Glo (Promega). The luminescence signals were recorded by the H1 synergy plate reader (Biotek) and were normalized against the control groups (100%) for which cells were only treated with the vehicle (PBS/1% DMSO).

## Acknowledgements

We thank the funding from the National Institute of Health (NIH) under the award number R35GM133468 (to R.E.W.), R01GM116204 (to W.Y.), R01GM22552 (to W.Y.), and R01GM123296 (to V.A.V.). We thank the support from the MD Anderson Cancer Center Leukemia SPORE P50 CA100632 (Career Enhancement Award) (to R.E.W.). Support for the NMR facility at Temple University by a CURE grant from the Pennsylvania Department of Health is gratefully acknowledged. High performance computing resources at Temple University were supported in part by the National Science Foundation through major research instrumentation grant CNS-1625061, the US Army Research Laboratory under contract number W911NF-16-2-0189, and NIH Research Resource computer instrumentation grant S10-OD020095. We also thank the participants of Folding@home, without whom this work would not be possible.

## Data availability

All the data supporting the findings of this study are available within the paper and the supplementary information, and are also available from the corresponding author upon request.

## Contributions

M.S.I., S.L.J., S.Z., V.V., W.Y., and R.E.W. designed the experiments. M.S.I., S.L.J., S.Z., Z.Y.B., Y.G., K.H.K., and Z.L conducted the experiments. M.S.I., S.L.J., S.Z., V.V., W.Y., and R.E.W. analyzed and discussed the results, wrote the manuscript. All authors reviewed the manuscript.

