## Supplementary for "Unprotected Peptide Macrocyclization and Stapling via A Fluorine-Thiol Displacement Reaction"

#### Table of Contents

Materials and Methods

Chemical Synthesis…………………………………………………… S2

Supplemental Tables…………………………………………………...... S7

Supplemental Figures…………………………………………………... S11

$^{1}$H- and $^{13}$C- NMR Spectra for New Compounds………………………………. S27

LC-Chromatogram and MS-Spectra for Peptides………………………………. S35

#### Materials and Methods:

##### Chemical Synthesis.

*General Procedures.* All chemicals and solvents used were purchased from either Fisher Scientific, Sigma Aldrich or VWR and were used directly without any further purification. All small molecules and building blocks were synthesized following traditional organic chemistry procedures. Regular phase flash column chromatography with manually loaded silica gel (grade 60, 230-400 mesh, Fisher Scientific) was used to purify synthesized compounds. High resolution ESI-MS was obtained at the Wistar Institute, using a ThermoFisher Scientific Q Exactive HF-X mass spectrometer coupled to a ThermoFisher Vanquish Horizon UHPLC system. NMR data was recorded on a 500 MHz Bruker Advance with TMS as an internal standard. Peptides were synthesized on solid-phase using Fmoc chemistry. After cleavage from resins, the crude mixtures were precipitated out by diethyl ether, resuspended, and purified using Waters 1525 series preparative high-performance liquid chromatography (HPLC) loaded with the XBridge Prep C18 column (25 cm x 19 mm, particle size 5 μm). All peptide mass spectra data were recorded on the Agilent 1100 series liquid chromatography-mass spectrometry (LC-MS) that was equipped with an Ascentis® Express C8 analytical column (5 cm x 2.1 mm, particle size 2.7 μm). The LC-MS program ran with mobile phase A (0.1% formic acid in water) and mobile phase B (0.1% formic acid in acetonitrile). For each run, the acetonitrile was linearly increased from 5% to 95% within 10 min.

*Small Molecule Model Reactions.* To a mixture of 20 μL of Tris buffer (3M) and 20 μL of DMF was added 10 μL of the stock DMF solution for the model compound **1** (640 mM). After 10 min of incubation at room temperature, 10 μL of methyl hydrazine, cysteine, or benzyl thiol (640 mM, DMF) together with 20 μL of water (for methyl hydrazine) or TCEP solution (2M) (for cysteine or benzyl thiol) were added. After adjusting its final pH to 9.0, the mixture was aliquoted equally into 4 sample vials (20 μL each), which were subsequently incubated at 37°C. At the desired time point (0,4h, 8h, and 12h) of incubation, one vial each was collected to run LC-MS analysis of the sample. To determine the reaction progress, the UV peak areas (254 nm) of the model compound **1** before and after the 12h incubation were integrated and the relative yields of the reaction were calculated based on the decreased percentage of the peak areas for compound **1**.

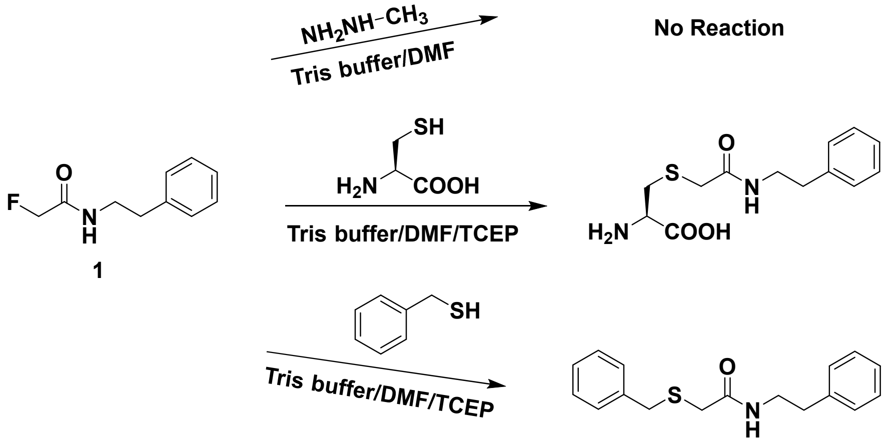




2

*Scheme S1:* Synthesis of 2-(9H-Fluoren-9-ylmethoxycarbonylamino)-3-(2-fluoro-acetylamino)-propionic acids (compounds **15**/**16**).

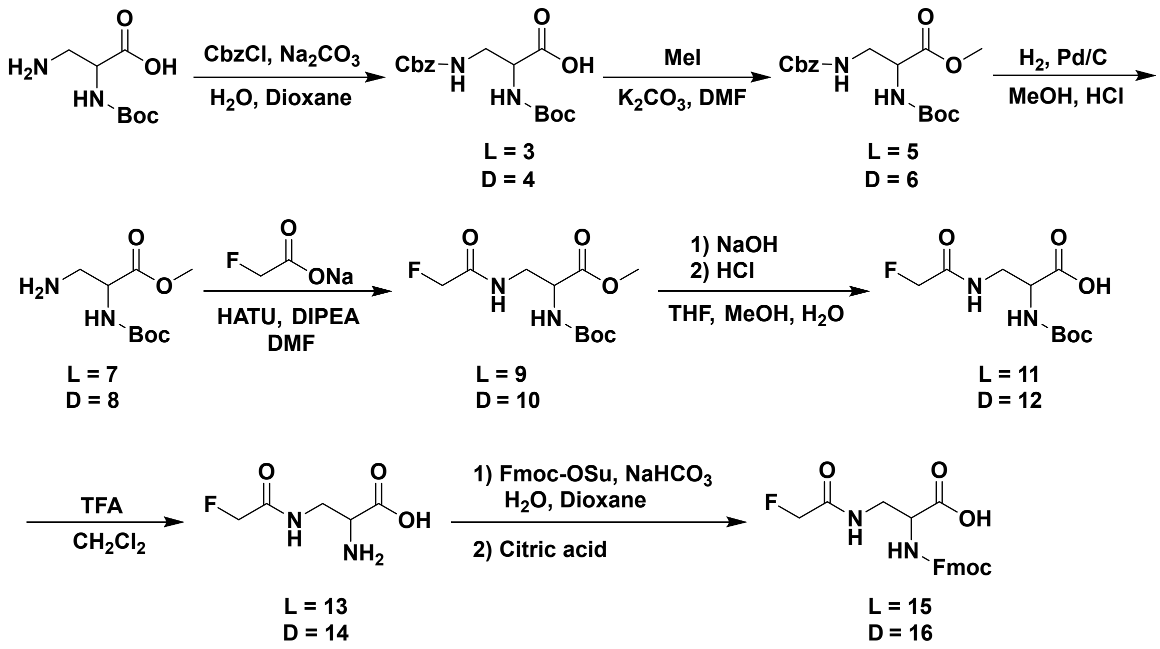


*Scheme S2:* Model fluorine-thiol displacement reaction (FTDR) between benzyl thiol and the fluoroacetamide-containing building block **9**.

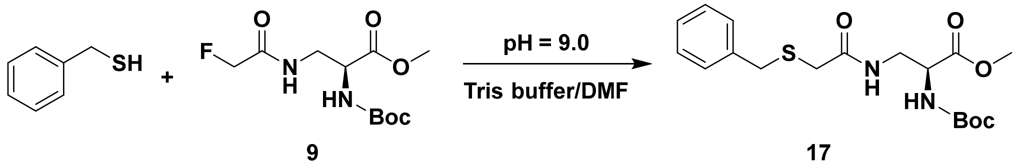


*Compound **1***. In a flame-dried 50 mL round bottom flask, sodium fluoroacetate (200 mg, 2 mmol) and HATU (912 mg, 2.4 mmol) were mixed with DIPEA (418μL, 2.4 mmol) in 15 mL anhydrous DMF. Around 10 min later, phenylethylamine (251 μL, 2 mmol) was added to the mixture. After overnight stirring, the reaction was quenched with water and the product was extracted with ethyl acetate. The collected organic layer was dried over anhydrous sodium sulfate, concentrated, and purified through silica gel column chromatography (hexane/ethyl acetate: 3/1) to afford the desired product as a white solid (277 mg, 1.52 mmol, 76% yield). $^1H$ NMR (500 MHz, $CDCl_3$): $\delta$ 7.34-7.31 (m, 2H), 7.26-7.20 (m, 3H), 4.76 (d, $J$ = 47.5 Hz, 2H), 3.60 (q, $J$ = 6.5 Hz, 2H), 2.86 (t, $J$ = 7.5 Hz, 2H); $^{13}C$ NMR (126 MHz, $CDCl_3$): $\delta$ 167.6 (d, $J$ = 17.1 Hz), 138.4, 128.8 (d, $J$ = 5.0 Hz), 126.7, 80.2 (d, $J$ = 186.1 Hz), 40.0, 35.6; $^{19}F$ NMR (471 MHz, $CDCl_3$): $\delta$ -224.78; HRMS (ESI) m/z calculated for $C_{10}H_{13}FNO$ $[M + H]^+$: 182.0976, found 182.0971.

*Compound **2***. The conversion of model compound **1** to the fluorine displaced product **2** was scaled up in 5 mL final volume following the procedure described in "*Small Molecule Model Reactions*". The reaction was quenched with 20 mL brine and the product was extracted with 30 mL of ethyl acetate three times. The organic layer was combined, dried, and concentrated under reduced pressure. The crude mixture was purified by flash column chromatography (25% ethyl acetate/hexane) to afford 80 mg white solid (70.7% yield). $^1$H NMR (500 MHz, $CDCl_3$): δ 7.12-7.39 (m, 10H), 6.73 (s, 1H), 3.57 (s, 2H,), 3.46 (q, 2H), 3.09 (s, 2H), 2.79 (t, 2H); $^{13}$C NMR (126 MHz, $CDCl_3$): δ 168.39, 138.65, 137.06, 128.97, 128.87, 128.77, 128.74, 127.46, 126.67, 40.71, 37.30, 35.43, 35.36. ESI-MS m/z calculated for $C_{17}H_{19}NOS$ $[M + H]^+$: 286.1, found 286.1.

*Compound 3*. Boc-Dap-OH (1 g, 4.9 mmol) was dissolved in the mixture of water (17 mL) and dioxane (47 mL). $Na_2CO_3$ (1.04 g, 9.8 mmol) in water (5 mL) was added to the flask, and the mixture was cooled on ice bath. Cbz-Cl (1.07 mL, 7.5 mmol) was then added dropwise, after which the mixture was stirred at room temperature overnight. The next day, dioxane was removed under reduced pressure, and the solution's pH was adjusted to approximately 9.0 with 1 M sodium hydroxide. The solution was extracted twice with ethyl acetate to remove the unreacted Cbz-Cl. The pH was subsequently adjusted to 4.0 with 1 M HCl and the acidic solution was then extracted three times by ethyl acetate, with the organic layers combined and dried over sodium sulfate. The organic phase was finally vacuum dried to afford 1.6 gram of crude product (96.5% yield) and was used directly for the next step without purification.

*Compound 4*. Compound **4** was synthesized following the same procedure for the enantiomer, compound **3**, with a 97.4% yield.

*Compound 5*. The previously synthesized compound **3** (1.6 g, 4.73 mmol) was dissolved in 20 mL of dry DMF. Potassium carbonate (2.29 g, 16.56 mmol) was added, and stirred at room temperature for 20 min. The reaction mixture was then cooled in an ice bath followed by the addition of methyl iodide (353 μL, 5.676 mmol). After overnight stirring at room temperature, the mixture was diluted with 200 mL water, and extracted three times with 60 mL ethyl acetate. The organic layers were combined, dried over anhydrous sodium sulfate, and concentrated under reduced pressure. The crude mixture was purified by flash column chromatography (ethyl acetate/hexane: 1/4) to finally afford 1.58 gram of product in viscous oil form (94% yield). $^1$H NMR (500 MHz, $CDCl_3$): $\delta$ 7.36-7.30 (m, 5H), 5.46 (s, 1H, br), 5.20 (s, 1H, br), 5.08 (s, 2H), 4.37 (s, 1H, br), 3.73 (s, 3H,), 3.58 (s, 2H, br) 1.42 (s, 9H); $^{13}$C NMR (126 MHz, $CDCl_3$): $\delta$ 171.18, 156.72, 155.43, 136.24, 128.56, 128.23, 128.15, 80.31, 67.03, 54.04, 52.71, 42.97, 28.30; HRMS(ESI) m/z calculated for $[C_{12}H_{17}N_2O_4$, M - Boc + H$]^+$ : 253.1188, found 253.1181.

*Compound 6*. Compound **6** has been synthesized following the same procedure for compound **5**, with a 93.5% yield. $^1$H NMR (500 MHz, $CDCl_3$): $\delta$ 7.35-7.28(m, 5H), 5.63 (s, 1H, br), 5.47 (s, 1H, br), 5.07 (s, 2H), 4.37 (s, 1H, br), 3.71 (s, 3H,), 3.56 (s, 2H, br) 1.43 (s, 9H); $^{13}$C NMR (126 MHz, $CDCl_3$): $\delta$ 171.29, 156.83, 155.56, 136.33, 128.52, 128.17, 128.12, 80.19, 66.94, 54.08, 52.63, 42.85, 28.29; HRMS(ESI) m/z calculated for $[C_{12}H_{17}N_2O_4$, M - Boc + H$]^+$: 253.1188, found 253.1182.

*Compound 7*. The previously synthesized compound **5** (1.58 g, 4.48 mmol) was dissolved in 20 mL of methanol and cooled on ice bath. A small amount of hydrogen chloride (370 μL) in methanol (5 mL) was added dropwise in order to quench the nucleophilic amine generated in situ during follow up hydrogenolysis. For hydrogenolysis, 10% Pd/C (200 mg) was added with subsequent stirring at room temperature under the hydrogen atmosphere. After overnight stirring, the reaction mixture was filtered through celite to remove Pd/C. The solvent was also evaporated under reduced pressure to afford crude compound **7** with almost quantitative yield. The crude product was used for the next step without any purification.

*Compound **8***. As an enantiomer to compound **7**, compound **8** has been synthesized following the same procedure for compound **7**.

*Compound **9***. Sodium fluoroacetate (100 mg, 1 mmol), HATU (418.3 mg, 1.1 mmol), and compound **7** (436.4 mg, 2 mmol) were dissolved in 10 mL of DMF and stirred at room temperature for 20 min. DIPEA (608.5 μL, 3.5 mmol) was then added. The mixture was stirred overnight, before the reaction was quenched with 100 mL of brine. After extraction (three times) with 50 mL ethyl acetate, the organic layers were combined and dried over anhydrous sodium sulfate. The crude was vacuum concentrated, loaded onto silica column and purified by ethyl acetate/hexane (1:2) to afford 215 mg of oil-like product with a 77.3% yield. $^{1}$H NMR (500 MHz, $CDCl_3$): $\delta$ 7.03 (s, 1H, br), 5.63 (s, 1H, br), 4.69-4.79 (d, *J*=50 Hz, 2H,), 4.39 (s, 1H, br), 3.71 (s, 3H,), 3.66 (t, *J*=5.0 Hz, 2H,), 1.38 (s, 9H); $^{13}$C NMR (126 MHz, $CDCl_3$): $\delta$ 170.95, 168.47, 155.69, 80.84, 80.35, 79.36, 53.54, 52.72, 40.92, 28.19; HRMS(ESI) m/z calculated for [$C_6H_{12}FN_2O_3$, M - Boc + H]$^+$: 179.0832, found 179.0824.

*Compound **10***. Compound **10** was prepared similarly to compound **9**, with a 75% yield. $^{1}$H NMR (500 MHz, $CDCl_3$): $\delta$ 6.94 (s, 1H, br), 5.55 (s, 1H, br), 4.81-4.72 (d, *J*=45 Hz, 2H,), 4.41 (s, 1H, br), 3.74 (s, 3H,), 3.68 (t, *J*=7.5 Hz, 2H,), 1.41 (s, 9H); $^{13}$C NMR (126 MHz, $CDCl_3$): $\delta$ 170.90, 168.41, 155.69, 80.87, 80.46, 79.40, 53.49, 52.81, 41.07, 28.23; HRMS(ESI) m/z calculated for [$C_6H_{12}FN_2O_3$, M - Boc + H]$^+$: 179.0832, found 179.0825.

*Compound **11***. Compound **9** (225 mg, 0.81 mmol) was dissolved in 4 mL of tetrahydrofuran and 2 mL of methanol. The solution was cooled in an ice-bath and 0.89 mL of 1 M sodium hydroxide (0.89 mmol) was added. After 15 min of stirring, the ice-bath was removed followed by subsequent stirring at room temperature for 1 h. The reaction was then quenched with 60 mL of water, and the mixture was washed once with 30 mL of ethyl acetate. The remaining mixture in the aqueous phase was cooled on ice-bath again, with the pH adjusted to 4.0 by 1 M HCl. At this point, the solution was extracted three times with 50 mL of ethyl acetate each. The organic layers were combined and dried over anhydrous sodium sulfate. The solvent was removed under reduced pressure by rotavapor, to afford 192.3 mg of crude product (~ 90% yield) that was used directly for the next step.

*Compound **12***. Compound **12** was synthesized following the same procedure for the enantiomer, **11**, with a 91% yield.

*Compound **13***. For Boc deprotection, compound **11** (200 mg, 0.76 mmol) was dissolved in 5 mL of dichloromethane and cooled in an ice-bath. Trifluoroacetic acid (2.5 mL) was added dropwise, and the mixture was kept stirring on the ice-bath for 10 min, after which the stirring was continued at room temperature for 1h. The solvent was removed under vacuum, to afford 111 mg of crude product with an 89.5% yield.

*Compound **14***. Compound **14** was prepared similarly from compound **12**, with an 88% yield.

*Compound **15***. Towards 200 mg compound **13** (1.22 mmol) in 3.5 mL of ice-cold 10% sodium carbonate solution, Fmoc-OSu (452 mg, 1.34 mmol) in 4 mL of dioxane was added dropwise. The mixture was left stirring overnight at room temperature. Dioxane was removed by vacuum, followed by the addition of 15 mL of water. The solution was washed with 10 mL of diethyl ether once. The aqueous phase was cooled with ice bath, and the solution's pH was adjusted to 3.5 by citric acid. Ethyl acetate was then used to extract the solution for three times (30 mL each). The combined organic phase was dried by anhydrous sodium sulfate, vacuum concentrated, and purified by flash column chromatography (2.5% Methanol /96.5% DCM /1% acetic acid). Finally, 396 mg of product in white solid form was obtained with a 91.5% yield. $^1$H NMR (500 MHz, $CD_3OD$): $\delta$ 7.72 (d, $J$=5 Hz, 2H), 7.60 (s, 2H, br), 7.32 (t, $J$=7.5 Hz, 2H), 7.25 (t, $J$=7.5 2H,), 4.70-4.79 (d, 1H), 4.35 (s, 1H, br), 4.27 (d, $J$=10 Hz, 2H), 4.15 (t, $J$=7.5 Hz, 1H,), 3.71-3.54 (m, 2H); $^{13}$C NMR (126 MHz, $CD_3OD$): $\delta$ 169.73, 169.58, 157.16, 143.86, 141.16, 127.39, 126.77, 124.85, 119.53, 80.35, 78.89, 66.77, 48.14, 39.63; HRMS(ESI) m/z calculated for [$C_{20}H_{20}FN_2O_5$, M + H]$^+$: 387.1356, found 387.1351.

*Compound **16***. Compound **16** has been synthesized following the same procedure for compound **15**, with a 90.7% final yield. $^1$H NMR (500 MHz, $CD_3OD$): $\delta$ 7.76 (d, J=10 Hz, 2H), 7.64 (m, 2H), 7.36 (t, $J$=7.5 Hz, 2H), 7.28 (t, $J$=7.5 Hz, 2H), 4.82-4.72 (dd, $J$=50 and 2.5 Hz, 2H), 4.37 (m, 1H), 4.32 (m, 2H), 4.20 (t, $J$=7.5 Hz, 1H), 3.74-3.56 (m, 2H); $^{13}$C NMR (126 MHz, $CD_3OD$): $\delta$ 171.99, 169.73, 157.17, 143.87, 141.18, 127.39, 126.77, 124.86, 119.52, 80.34, 78.83, 66.77, 48.12, 39.60; HRMS(ESI) m/z calculated for [$C_{20}H_{20}FN_2O_5$, M + H]$^+$: 387.1356, found 387.1350.

*Compound **17***. Towards a mixture of 2 mL of DMF and 4 mL of Tris (3M solution) were added 470 µL of compound **9** (600 mM in DMF), 500 µL of benzyl thiol (600 mM in DMF), and 525 µL of TCEP solution (600 mM). After the final pH was adjusted to 9.0, the mixture was left stirring at 37 °C for 12h. The reaction was then quenched with 20 mL of brine and the product was extracted with 30 mL of ethyl acetate three times. The organic layers were combined, dried over anhydrous sodium sulfate, and concentrated under reduced pressure. The crude mixture was purified by flash column chromatography (25% ethyl acetate/hexane) to finally afford 50.2 mg of compound **17** as a white solid (73% yield). $^1$H NMR (500 MHz, $CDCl_3$): δ 7.24-7.34 (m, 5H), 7.08 (s,1H), 5.49 (s, 1H), 4.39 (m, 1H,), 3.77 (s, 3H), 3.72 (s, 2H), 3.58 (m, 2H), 3.11 (s, 2H), 1.45 (s, 9H); $^{13}$C NMR (126 MHz, $CDCl_3$): δ 171.09, 169.48, 155.57, 136.95, 129.07, 128.75, 127.90, 80.39, 53.94, 52.78, 41.62, 36.93, 35.04, 28.31; HRMS(ESI) m/z calculated for [$C_{13}H_{19}N_2O_3S$, M - Boc + H]$^+$ : 283.1116, found 283.1109.

Table S1. Summary of the molecular weights (MWs) and yields of stapled Axin analogues.

| Peptide | Theoretical MW in Da | Observed MW [M+2H]/2 | Observed MW [M+3H]/3 | % yield |
|---|---|---|---|---|
| **21** | 2007.9 | 1005.1 | 670.5 | 54.8% |
| **22** | 2021.9 | 1012.2 | 675.2 | 59.2% |
| **23** | 2036.0 | 1019.2 | 679.9 | 48.6% |
| **24** | 2050.0 | 1026.1 | 684.5 | 51.9% |
| **25** | 2041.9 | 1022.2 | 681.9 | 62.6% |
| **26** | 2041.9 | 1022.1 | 681.8 | 52.2% |
| **28** | 2007.9 | 1005.2 | 670.5 | 58.2% |
| **29** | 2021.9 | 1012.2 | 675.2 | 61.7% |
| **30** | 2036.0 | 1019.1 | 679.9 | 60.4% |
| **31** | 2050.0 | 1026.1 | 684.6 | 55.3% |
| **32** | 2041.9 | 1022.2 | 681.9 | 59.3% |
| **33** | 2041.9 | 1022.2 | 681.9 | 64.9% |
| **35** | 2036.0 | 1019.1 | 679.9 | 53.9% |
| **36** | 2050.0 | 1026.1 | 684.6 | 43.2% |
| **37** | 2041.9 | 1022.2 | 681.9 | 58.4% |
| **39** | 2036.0 | 1019.1 | 679.9 | 39.4% |
| **40** | 2050.0 | 1026.1 | 684.5 | 49.8% |
| **41** | 2041.9 | 1022.2 | 681.9 | 50.9% |
| **42**[*] | 1870.0 | 936.2 | 624.5 | 73.4% |

*: Control peptides stapled by ring-closing metathesis.

Table S2. Summary of the molecular weights (MWs) and yields of stapled peptides for cell permeability studies.

| Peptide | Theoretical MW in Da | Observed MW [M+2H]/2 | Observed MW [M+3H]/3 | Observed MW [M+4H]/4 | Observed MW [M+5H]/5 | Observed MW [M+6H]/6 | % yield |
|---|---|---|---|---|---|---|---|
| **43** | 2081.8 | 1042.2 | | | | | 59.4% |
| **44** | 2081.8 | 1042.2 | | | | | 51.1% |
| **45** | 2081.8 | 1042.3 | | | | | 55.7% |
| **46*** | 1909.9 | 956.4 | | | | | 51.2% |
| **48** | 2871.3 | | 958.5 | 719.2 | 575.6 | 479.8 | 53.6% |
| **50** | 2871.3 | | 958.5 | 719.1 | 575.6 | 479.7 | 52.9% |
| **51*** | 2699.4 | | 901.3 | 676.2 | 542.5 | 451.1 | 62.8% |

*: Control peptides stapled by ring-closing metathesis.

Table S3. Summary of simulation trajectory data for stapled/unstapled Axin and HIV peptides.

| peptide | Fluoro substrates | Simulation time (μs) |
|---|---|---|
| **20** | $L_{i,\ i+4}L$ | 77.40 |
| **21** | $L_{i,\ i+4}L$ | 76.25 |
| **22** | $L_{i,\ i+4}L$ | 82.10 |
| **23** | $L_{i,\ i+4}L$ | 76.90 |
| **24** | $L_{i,\ i+4}L$ | 84.35 |
| **25** | $L_{i,\ i+4}L$ | 75.20 |
| **26** | $L_{i,\ i+4}L$ | 82.25 |
| **27** | $D_{i,\ i+4}D$ | 62.60 |
| **28** | $D_{i,\ i+4}D$ | 74.10 |
| **29** | $D_{i,\ i+4}D$ | 71.85 |
| **30** | $D_{i,\ i+4}D$ | 83.45 |
| **31** | $D_{i,\ i+4}D$ | 76.55 |
| **32** | $D_{i,\ i+4}D$ | 76.75 |
| **33** | $D_{i,\ i+4}D$ | 76.40 |
| **34** | $L_{i,\ i+4}D$ | 68.10 |
| **35** | $L_{i,\ i+4}D$ | 78.40 |
| **36** | $L_{i,\ i+4}D$ | 84.10 |
| **37** | $L_{i,\ i+4}D$ | 65.25 |
| **38** | $D_{i,\ i+4}L$ | 66.50 |
| **39** | $D_{i,\ i+4}L$ | 79.45 |
| **40** | $D_{i,\ i+4}L$ | 82.05 |
| **41** | $D_{i,\ i+4}L$ | 84.80 |
| **43** | $L_{i,\ i+4}L$ | 51.65 |
| **44** | $L_{i,\ i+4}D$ | 60.00 |
| **45** | $D_{i,\ i+4}L$ | 63.30 |
| | total | 1859.75 |
| | average | 74.39 |

Table S4. Quantification of the cell penetration of HIV C-CA binding peptides.

| Peptide | Mean Intensity Ratio | N (cell number) | Peptide | Mean Intensity Ratio | N (cell number) |
|---|---|---|---|---|---|
| **46*** | 1.00 ± 0.04 | 108 | **44** | 5.75 ± 0.07 | 121 |
| **43** | 4.76 ± 0.09 | 81 | **45** | 3.07 ± 0.08 | 115 |

*: Control peptides stapled by ring-closing metathesis.

Table S5. Quantification of the cell penetration of Axin analogues.

| Peptide | Mean Intensity Ratio | N (cell number) | Peptide | Mean Intensity Ratio | N (cell number) |
|---|---|---|---|---|---|
| **51*** | 1.00 ± 0.02 | 91 | **49** | 0.33 ± 0.01 | 102 |
| **47** | 0.36 ± 0.02 | 102 | **50** | 5.05 ± 0.14 | 110 |
| **48** | 4.86 ± 0.15 | 101 | **FITC only** | 0.30 ± 0.02 | 104 |

*: Control peptides stapled by ring-closing metathesis.

Table S6. Quantification of the cell penetration of stapled Axin analogue **48** during pathway blocking studies.

| Pathway Blocker | Mean Intensity Ratio | N | Pathway Blocker | Mean Intensity Ratio | N |
|---|---|---|---|---|---|
| Control* | 1.00 ± 0.04 | 52 | Cytochalasin D | 0.10 ± 0.01 | 60 |
| Nystatin | 1.16 ± 0.03 | 48 | $NaClO_3$ | 0.12 ± 0.01 | 63 |
| Chlorpromazine | 0.13 ± 0.01 | 61 | | | |

*: Control: solvent vehicle only.

Table S7. Quantification of the cell penetration of stapled Axin analogue **50** during pathway blocking studies.

| Pathway Blocker | Mean Intensity Ratio | N | Pathway Blocker | Mean Intensity Ratio | N |
|---|---|---|---|---|---|
| Control* | 1.00 ± 0.02 | 54 | Cytochalasin D | 0.99 ± 0.02 | 56 |
| Nystatin | 0.17 ± 0.01 | 59 | $NaClO_3$ | 0.20 ± 0.01 | 49 |
| Chlorpromazine | 0.16 ± 0.01 | 53 | | | |

*: Control: solvent vehicle only.

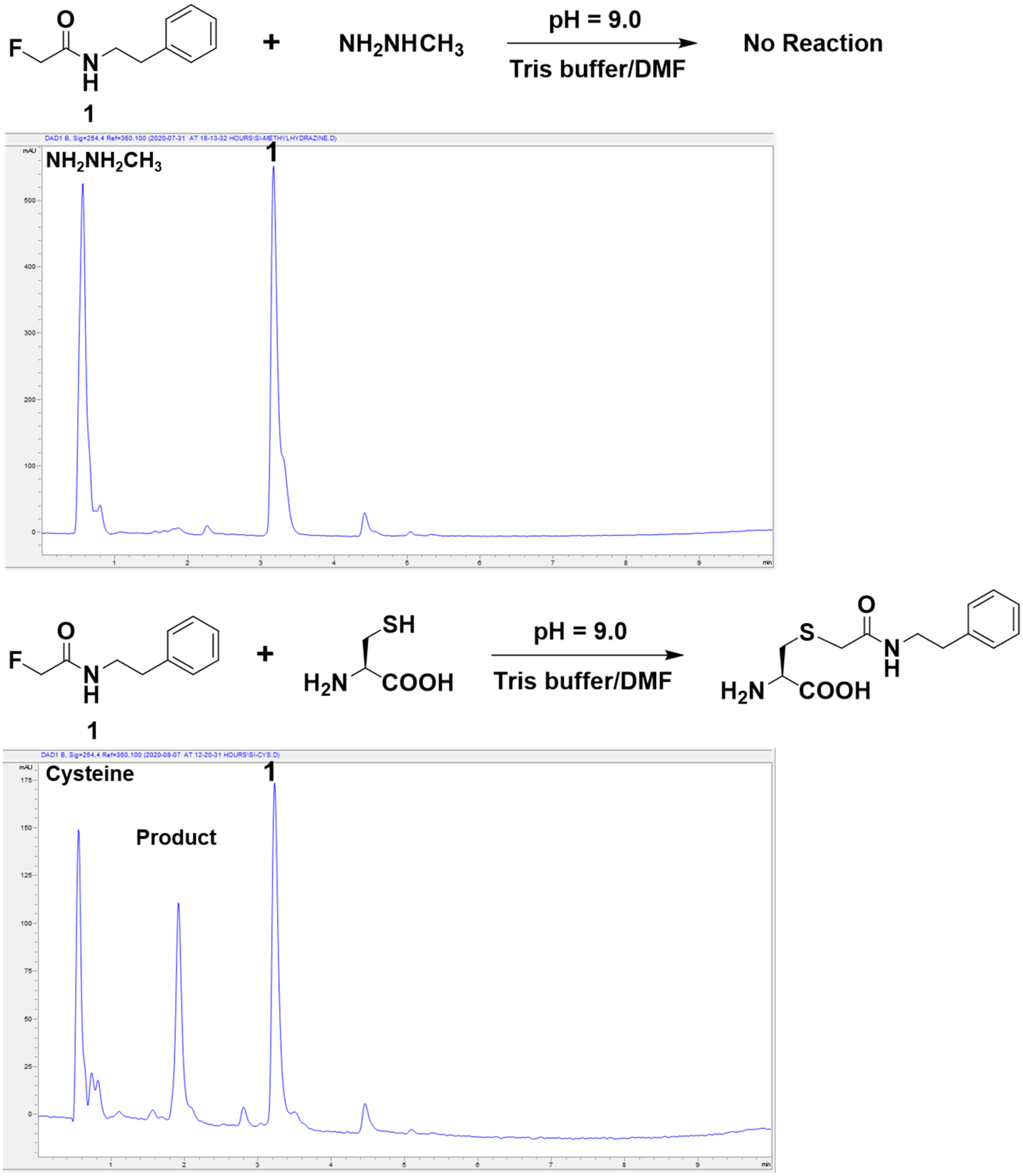


**Figure S1.** The reaction of the fluoroacetamide-containing model compound **1** with methyl hydrazine and cysteine, respectively. For the reaction with cysteine, the mixture contained 500 mM TCEP to ensure a reducing environment. The reaction progress was monitored by LC-MS after 12h of incubation at 37ºC.

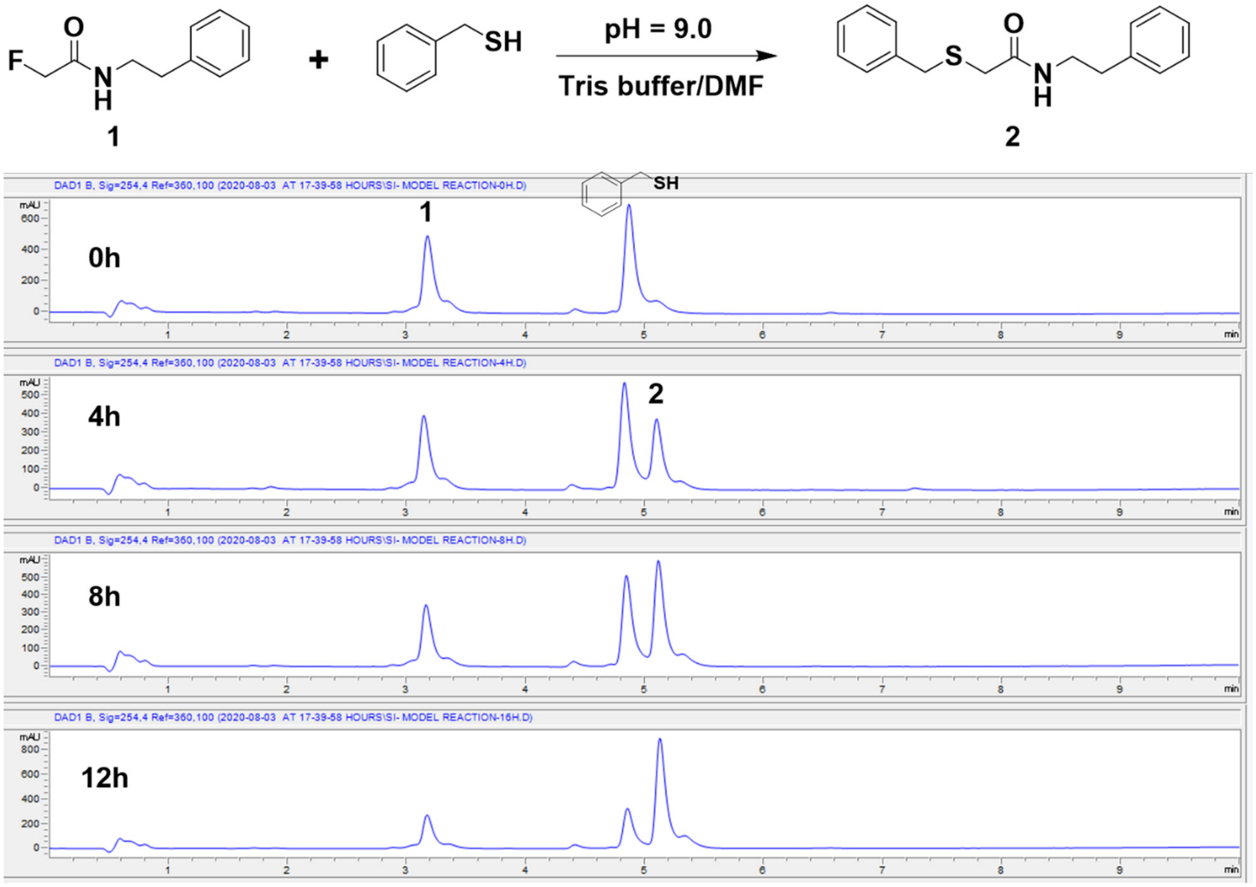


**Figure S2.** Time-dependent fluorine displacement reaction between the model compound **1** and benzyl thiol. The mixture contained 500 mM TCEP in the Tris/DMF solution. The reaction progress was monitored by LC-MS.

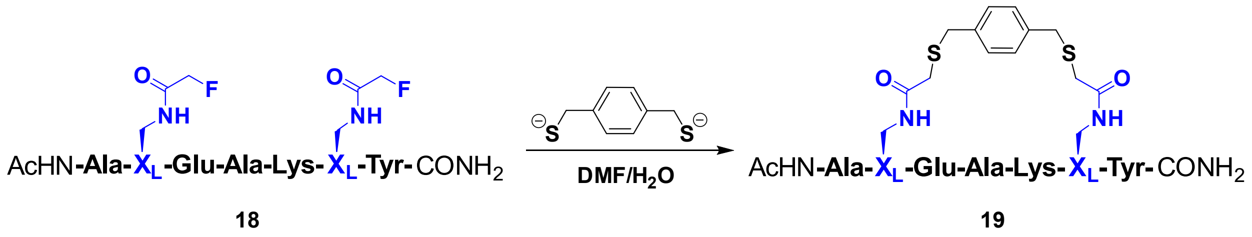


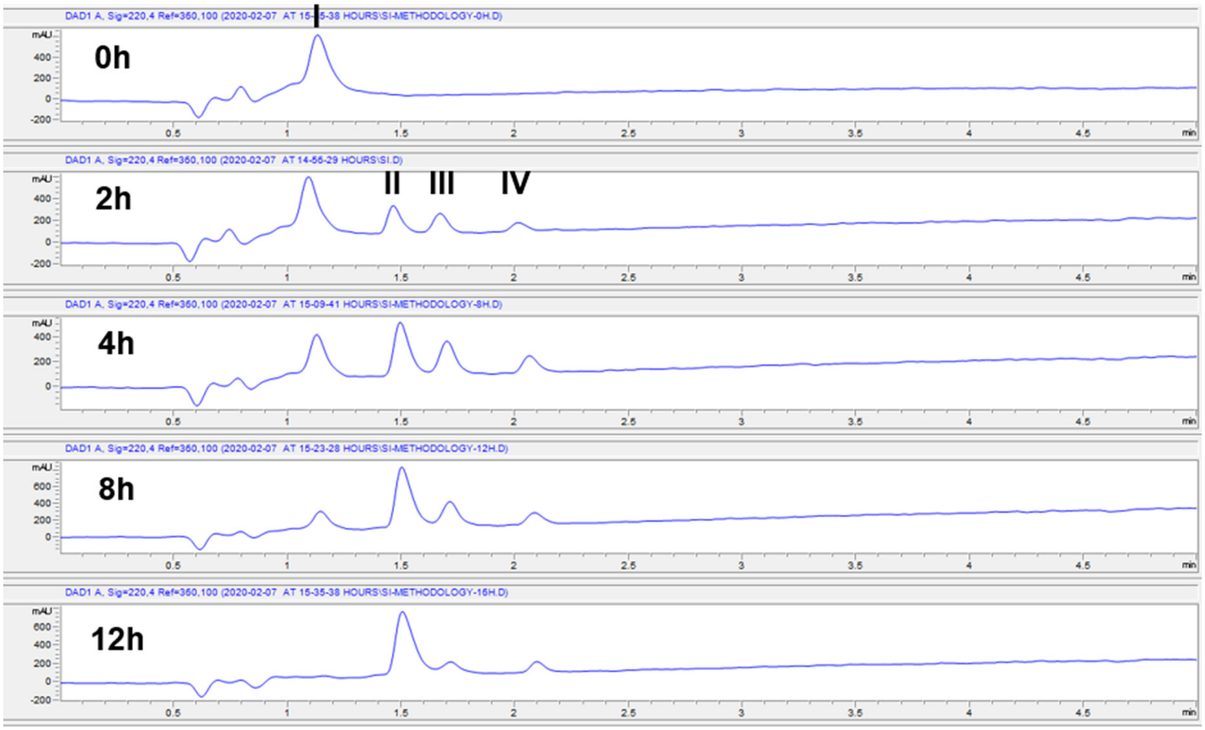


**Figure S3.** Time-dependent stapling of the linear model peptide **18** with the linker 1,4-benzenedimethanethiol. The reaction progress was monitored by LC-MS. The peak **I** is starting peptide **18**; peak **II** is the desired product **19**; peak **III** is the noncyclic byproduct modified by one equivalent of linker; peak **IV** is the noncyclic byproduct modified by two equivalents of linkers.

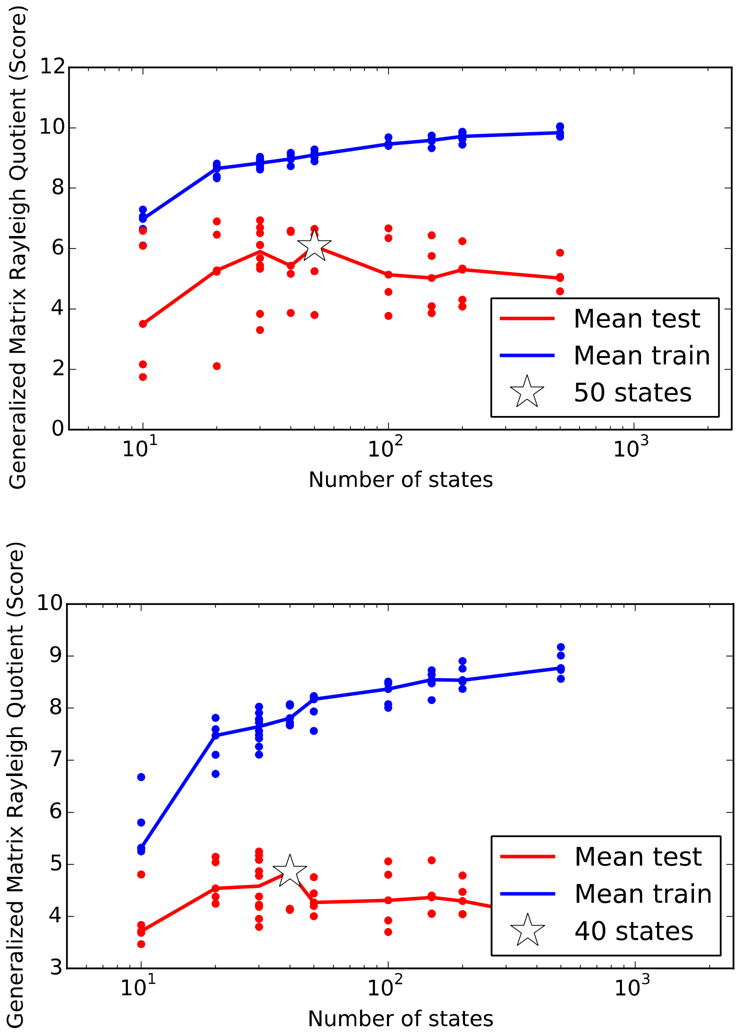


**Figure S4.** Generalized matrix Rayleigh quotient (GMRQ) scores, shown versus the number of states used to construct the MSMs, shown for MSMs of 1,3-benzenedimethanethiol stapled (L,D) Axin peptide (top) and unstapled (L,D) Axin peptide (bottom). Five-fold cross-validation was used, where the MSM was trained on 4/5 of the input data, and the GMRQ score computing using the remaining 1/5 of the data, in five separate trials. While the mean GMRQ score for the training data (blue) continues to increase with the number states, the mean score for the testing data reaches a peak at around 50 states (marked with a star). Based on the general consistency of these results for most of the peptide systems, we chose to construct MSMs using 50 states.

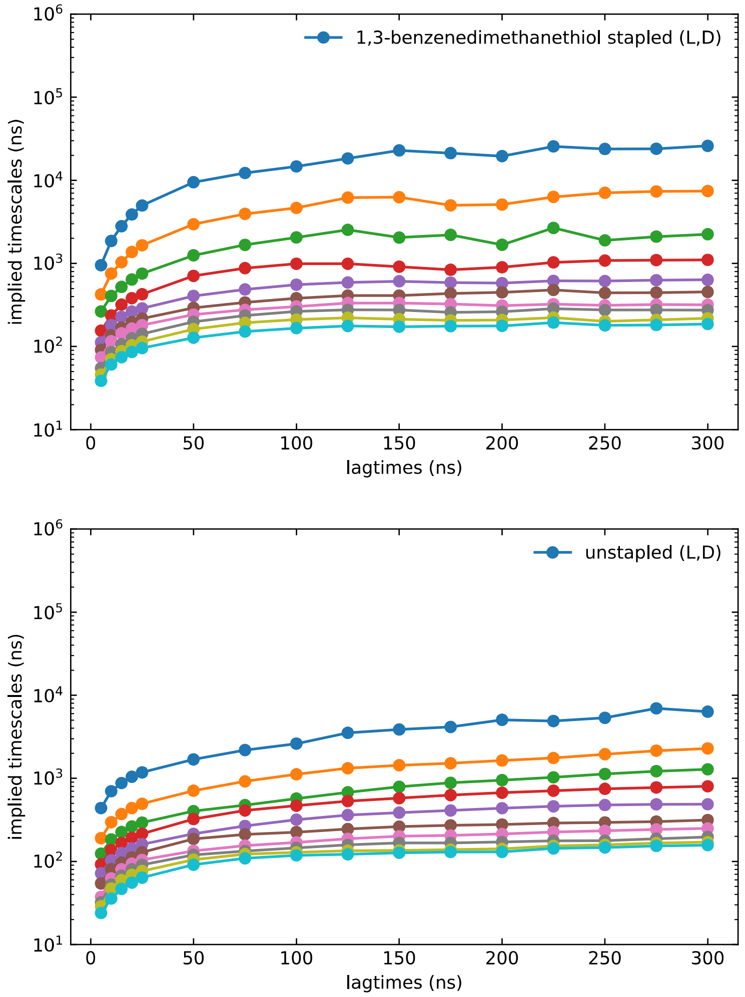


**Figure S5.** Slowest ten implied timescales for MSMs of 1,3-benzenedimethanethiol stapled (L,D) Axin peptide (top) and unstapled (L,D) Axin peptide (bottom), plotted as a function of lagtime, $\tau$. Implied timescales are calculated as $t_i = \tau/\ln \lambda_i$ where $\lambda_i$ are the eigenvalues of the MSM transition matrix. Uncertainties are estimated using a per-trajectory bootstrap procedure.

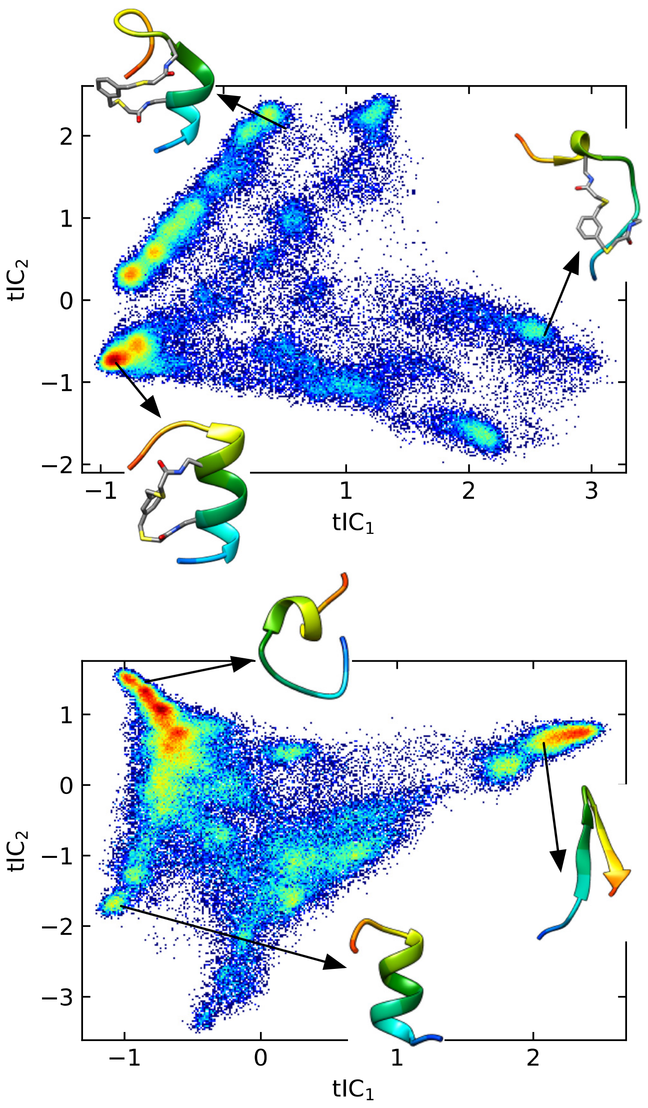


**Figure S6**. Examples of trajectory data projected on the first two tICs, with representative structures, for 1,3-benzenedimethanethiol stapled (L,D) Axin peptide (top), and unstapled (L,D) Axin peptide (bottom). The color scale represents regions of high (red) to low (blue) density.

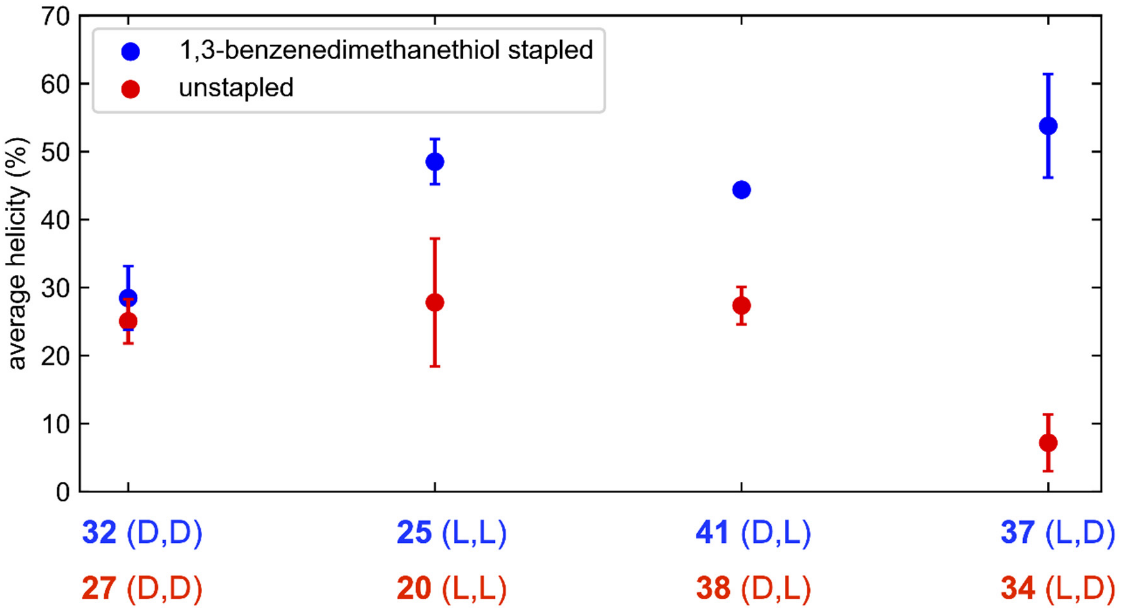


**Figure S7**. Comparison between the helicities of 1,3-benzenedimethanethiol stapled Axin peptides (blue circles) and unstapled Axin peptides (red circle). Uncertainties (vertical bars) were calculated using a bootstrap procedure in which five different MSMs were constructed by sampling the input trajectory data with replacement.

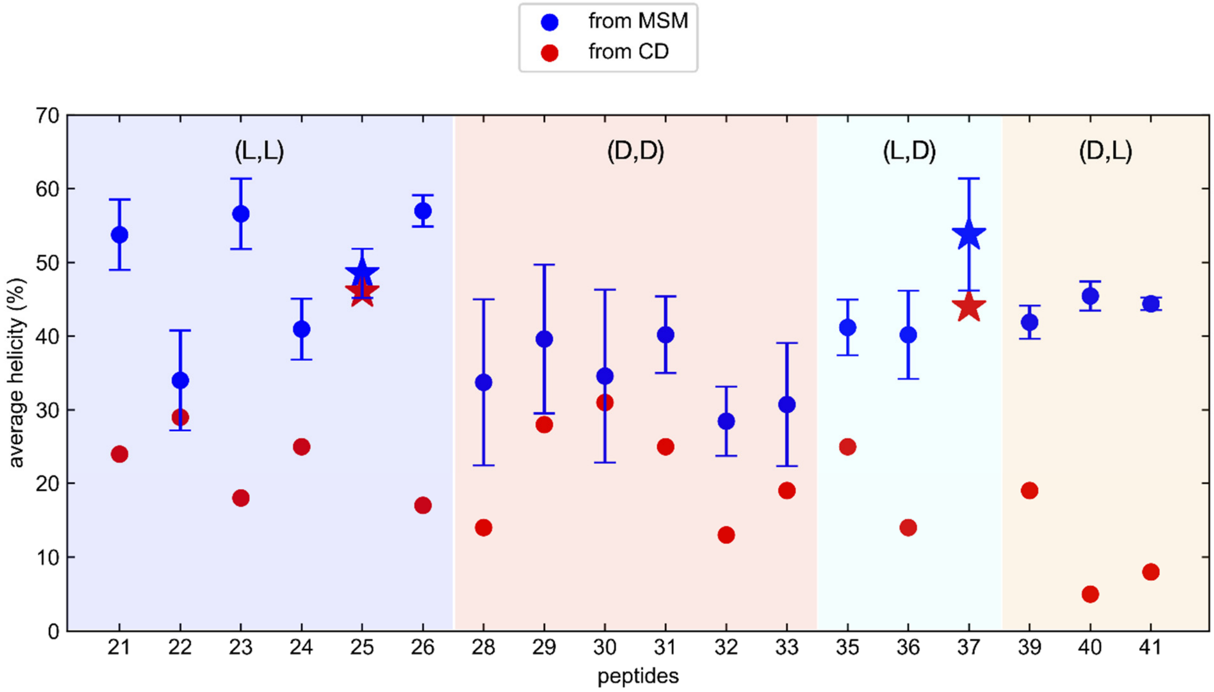


**Figure S8**. Comparison of predicted and experimental helicities for stapled Axin peptides **21-26**, **28-33**, **35-37**, and **39-41**. Uncertainties (vertical bars) were calculated using a bootstrap procedure in which five different MSMs were constructed by sampling the input trajectory data with replacement. Peptides **25** and **37** are labeled with star makers to denote the highest experimentally measured helicities.

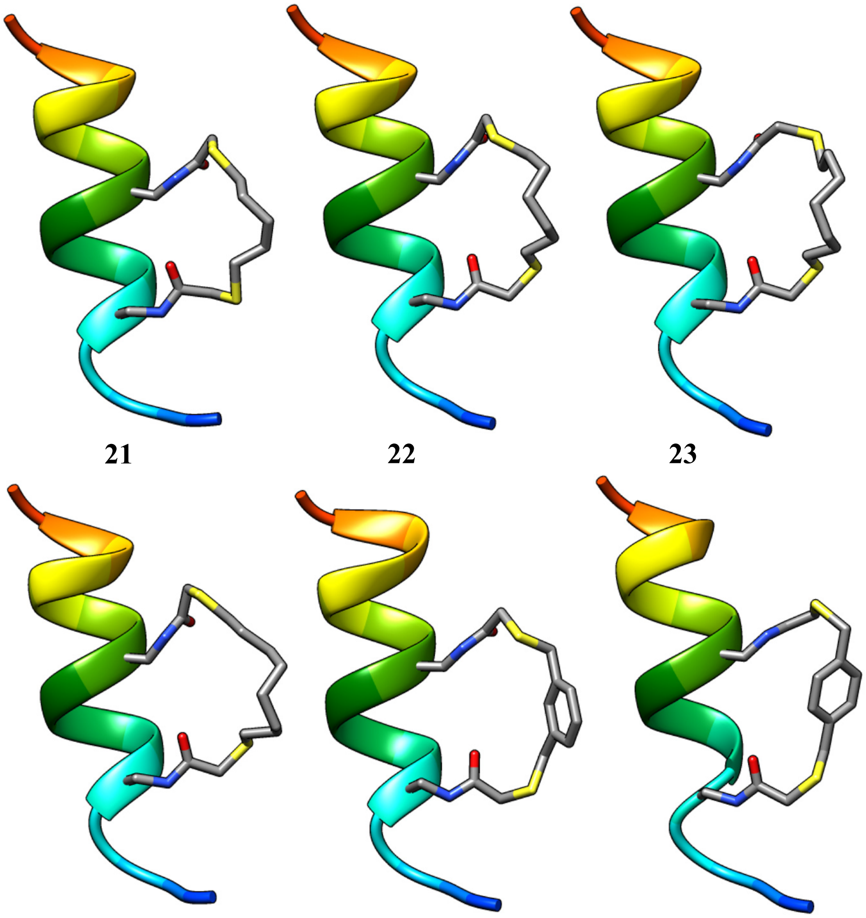
21 22 23 24 25 26

**Figure S9**. Minimized structures of stapled Axin peptides crosslinked at the *i*, *i*+4 positions.

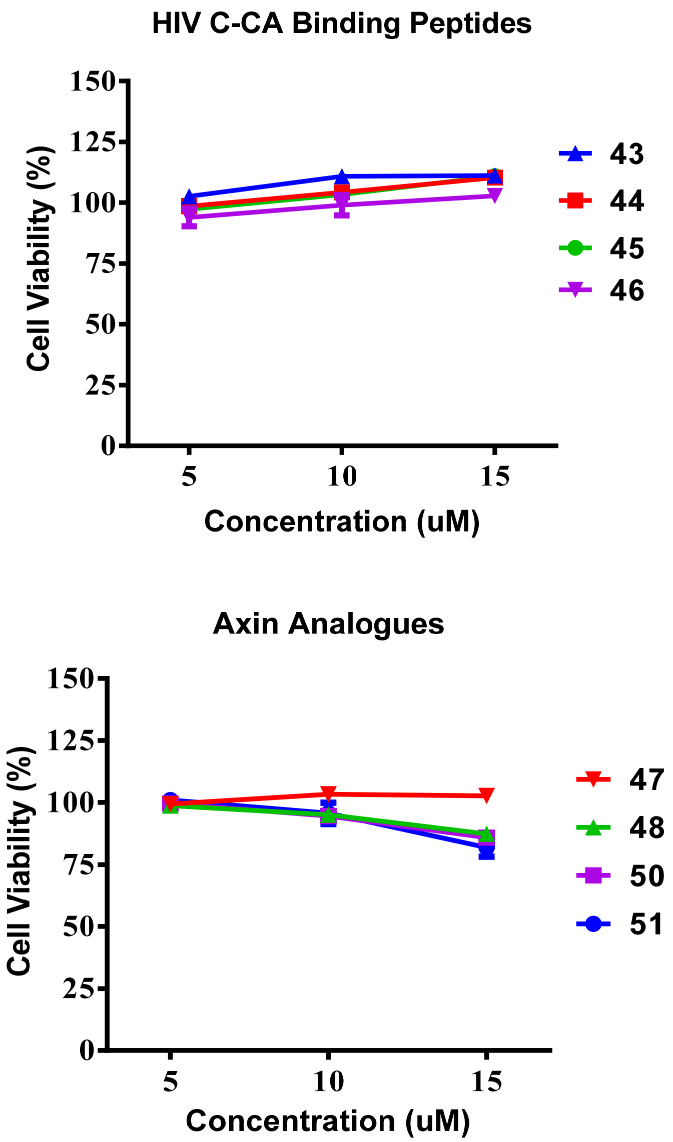


**Figure S10.** Viability of cells after treatment with FITC-labelled peptide analogues. HEK293T cells were treated with HIV C-CA binding peptides, while DLD-1 cells were treated with Axin analogues at doses of 5, 10, or 15 μM. In each set, the cells treated with solvent only were normalized as the 100% control.

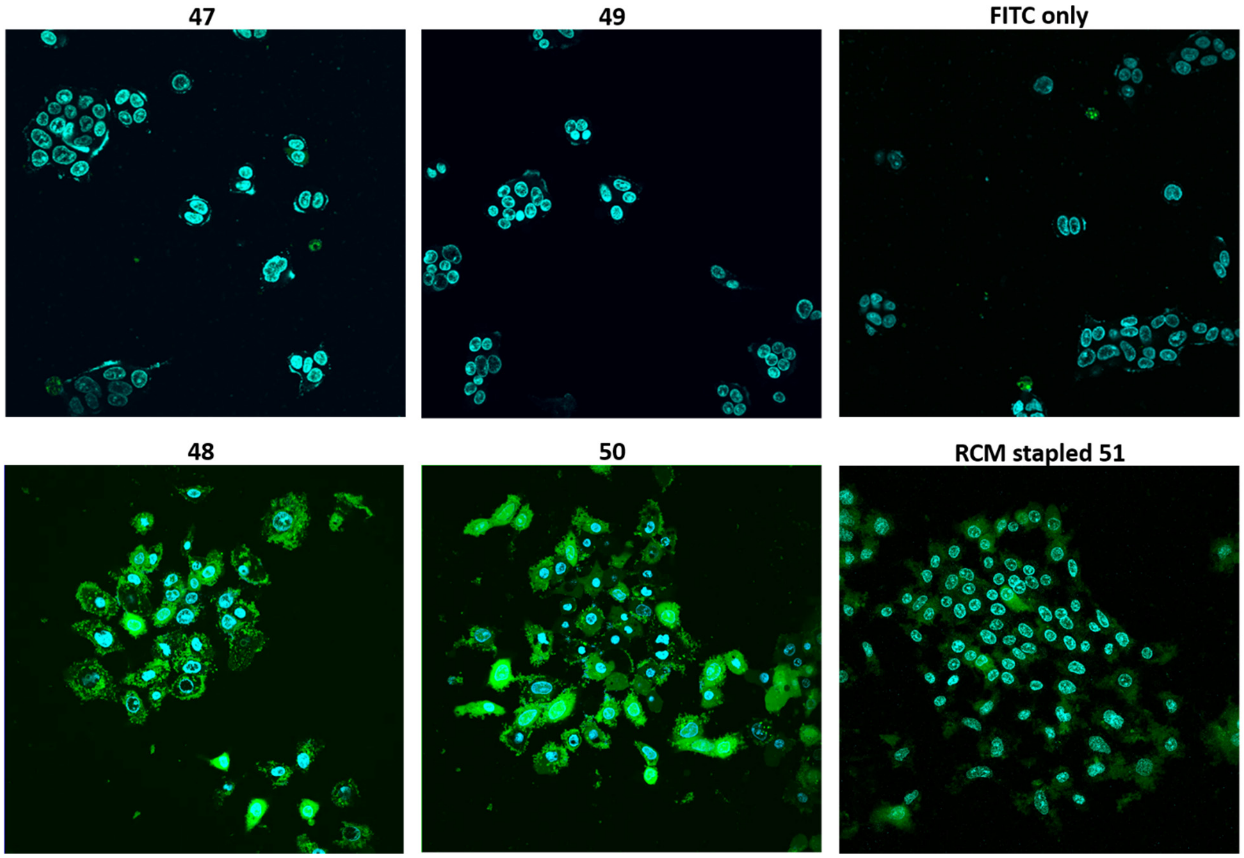


**Figure S11.** Raw imaging data for DLD-1 cells treated with 10 μM of FITC-labelled Axin analogues.

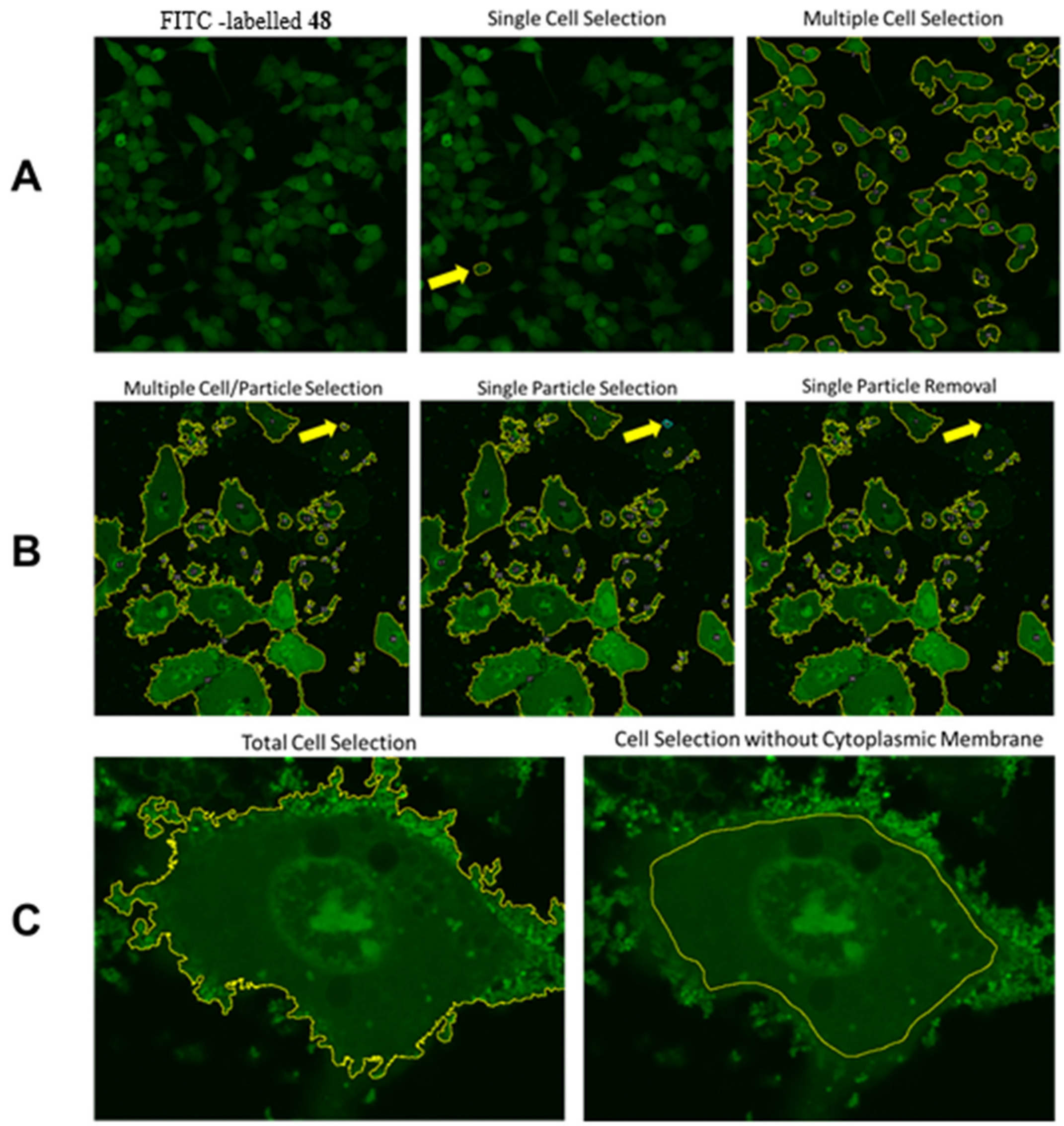


**Figure S12.** Representative cell selection for FITC-labelled Axin analogue **48** using the NIH ImageJ analyze particles function. A) Single and multiple cell selection for the FITC-labeled peptide. B) Extracellular particle selection and removal. C) The removal of the cytoplasmic membrane from the mean intensity measurement.

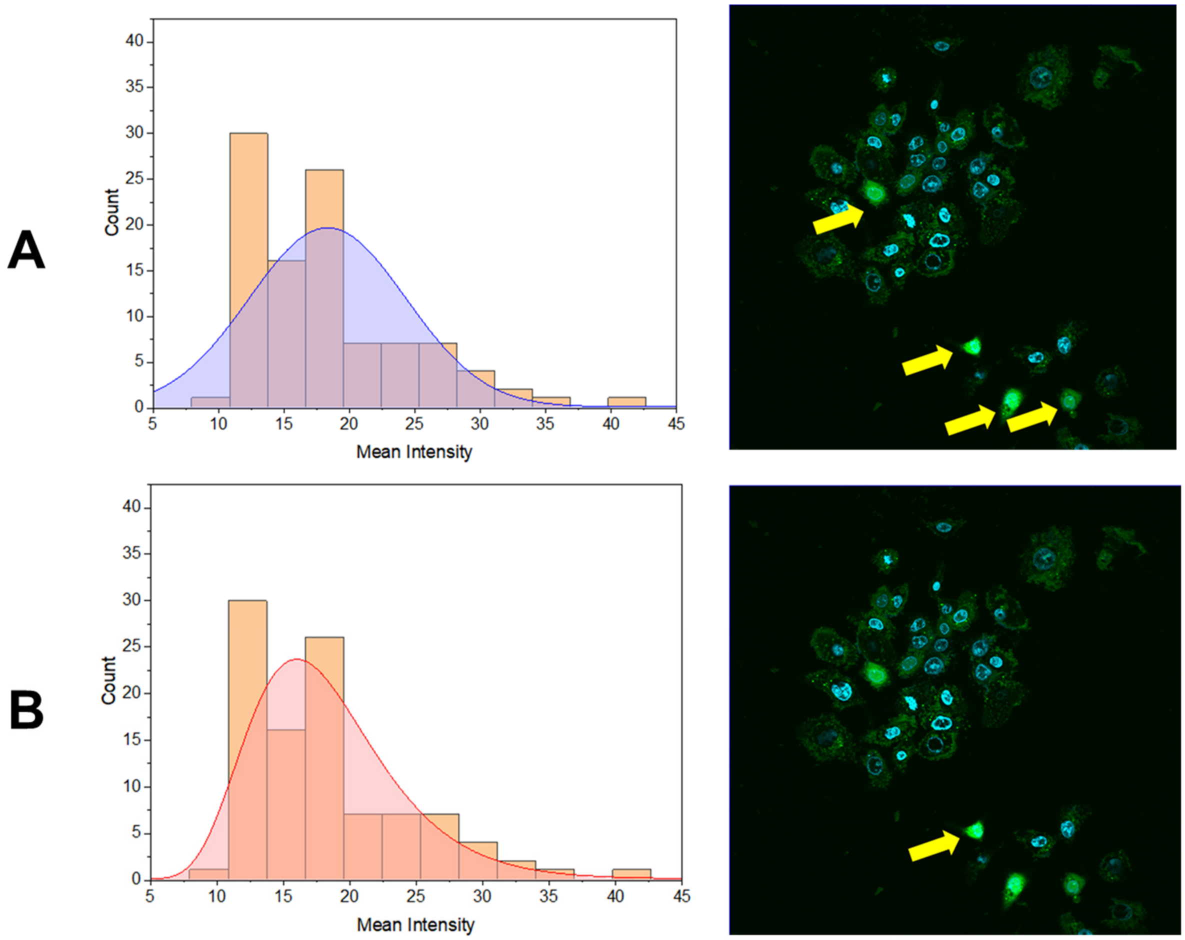


**Figure S13.** Outlier exclusion for cells treated with FITC-labelled Axin analogue **48**. A) Histogram of the mean intensities of the FITC-labeled peptide with a normal distribution curve applied. With a normal distribution, four cells (yellow arrows) with values 31.13, 32.41, 35.64, and 42.38 is excluded from the data set. B) Histogram of the mean intensities of the FITC-labeled peptide with the best fit, lognormal distribution curve applied. With a lognormal distribution, one cell (yellow arrow) with a value 42.38 is excluded from the data set.

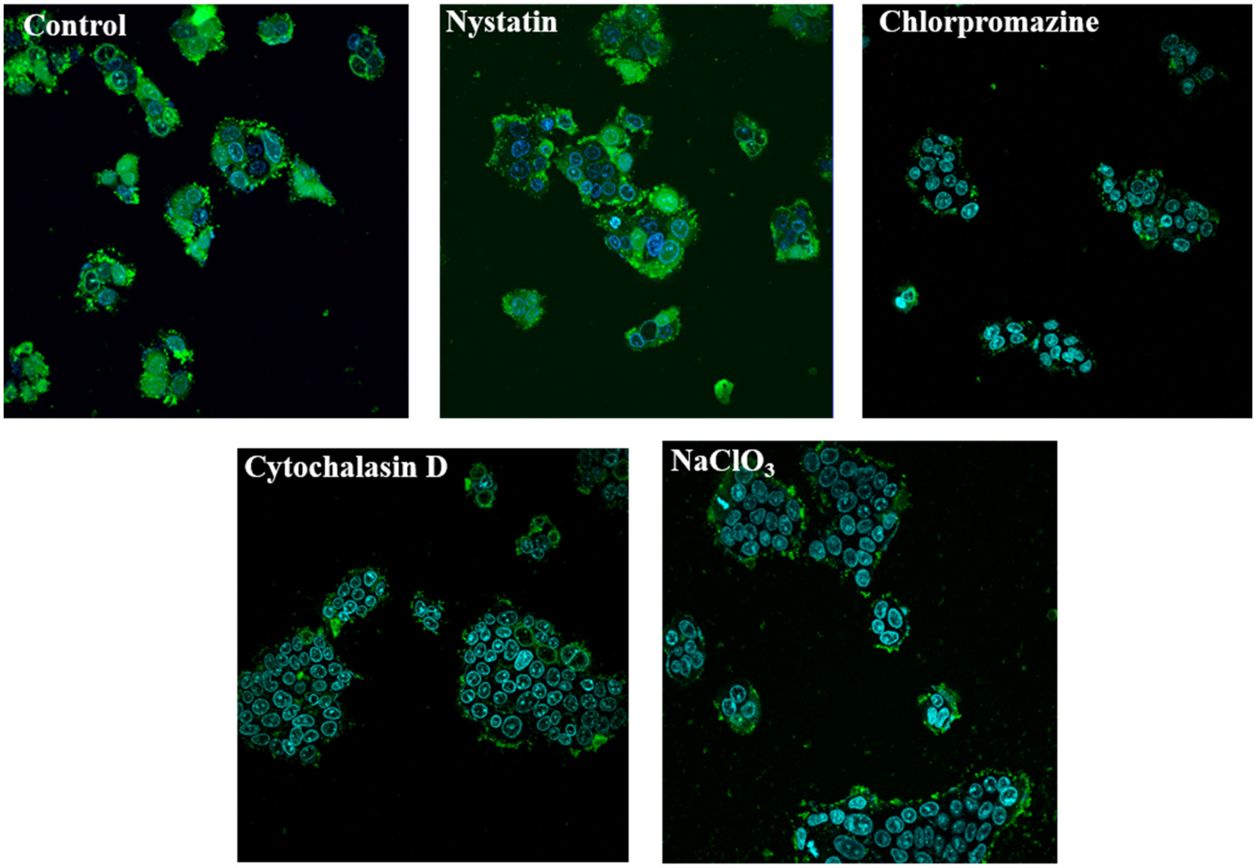


**Figure S14.** Fluorescent confocal microscopy images of the DLD-1 cells treated with the stapled Axin analogue **48** in the presence of different endocytotic pathway blockers (Nystatin, Chlorpromazine, Cytochalasin D, and $NaClO_3$). Green: FITC-labelled peptide **48**; Blue: nucleus stained by Hoechst 33342. Control: cells pretreated with vehicle control and then incubated with peptide **48**.

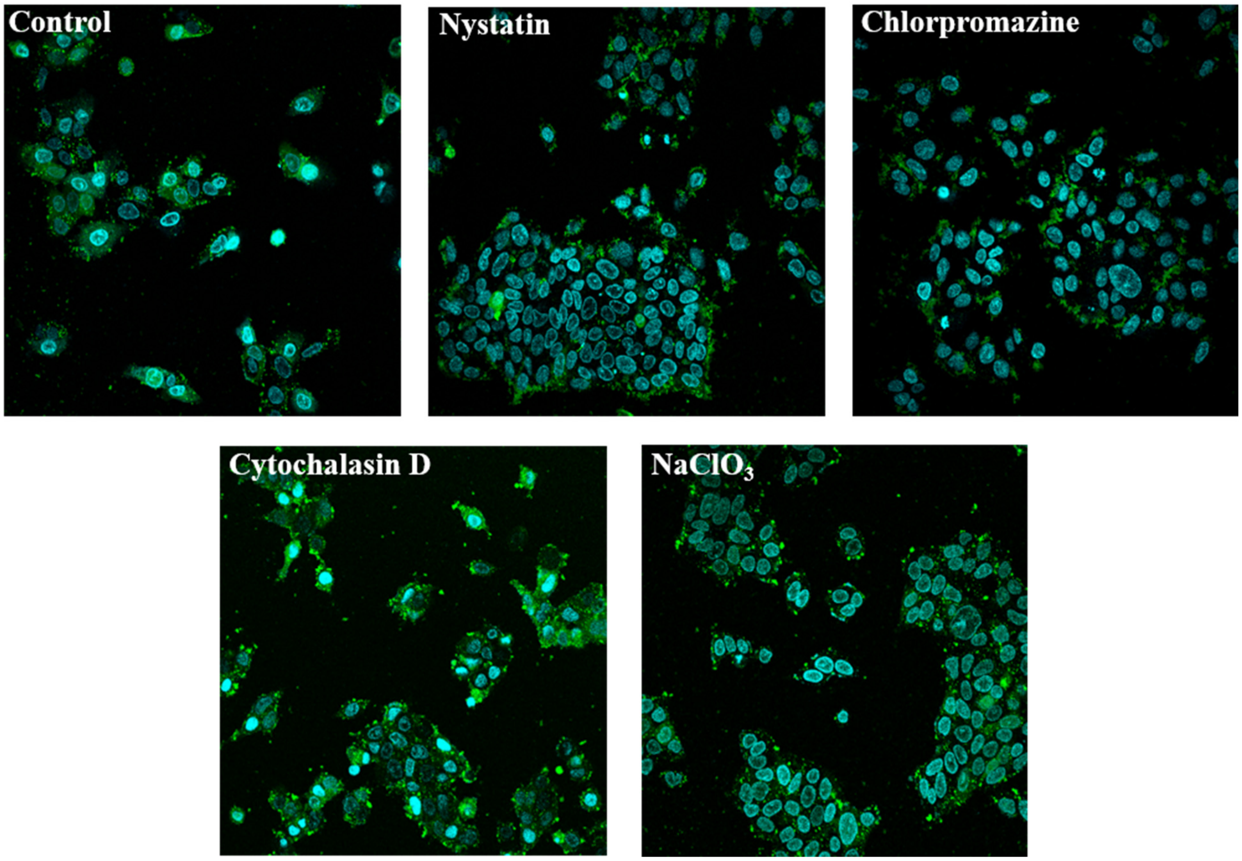


**Figure S15.** Fluorescent confocal microscopy images of the DLD-1 cells treated with the stapled Axin analogue **50** in the presence of different endocytotic pathway blockers (Nystatin, Chlorpromazine, Cytochalasin D, and $NaClO_3$). Green: FITC-labelled peptide **50**; Blue: nucleus stained by Hoechst 33342. Control: cells pretreated with vehicle control and then incubated with peptide **50**.

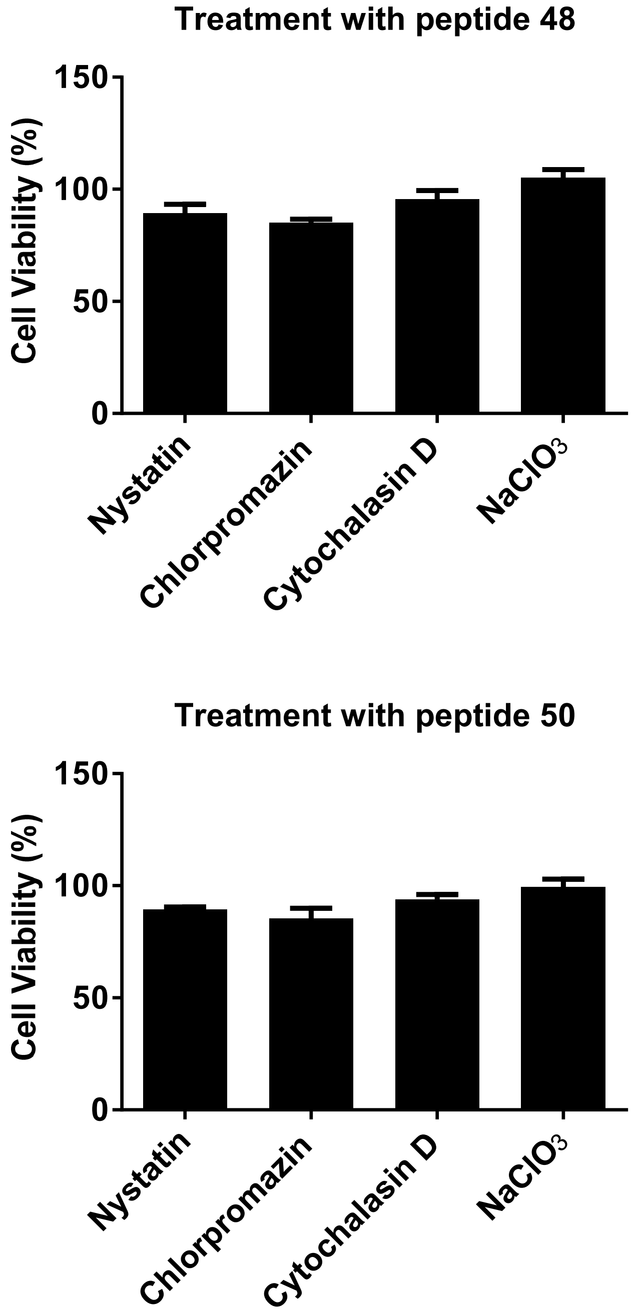


**Figure S16.** Viability of DLD-1 cells after imaging studies with representative stapled Axin analgoues (**48** or **50**). Cells were pre-treated with different small molecule blockers of endocytic pathways before imaging studies. The viability of cells treated with only vehicle control were normalized as 100%.

### NMR Spectra

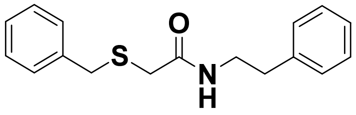


**2**

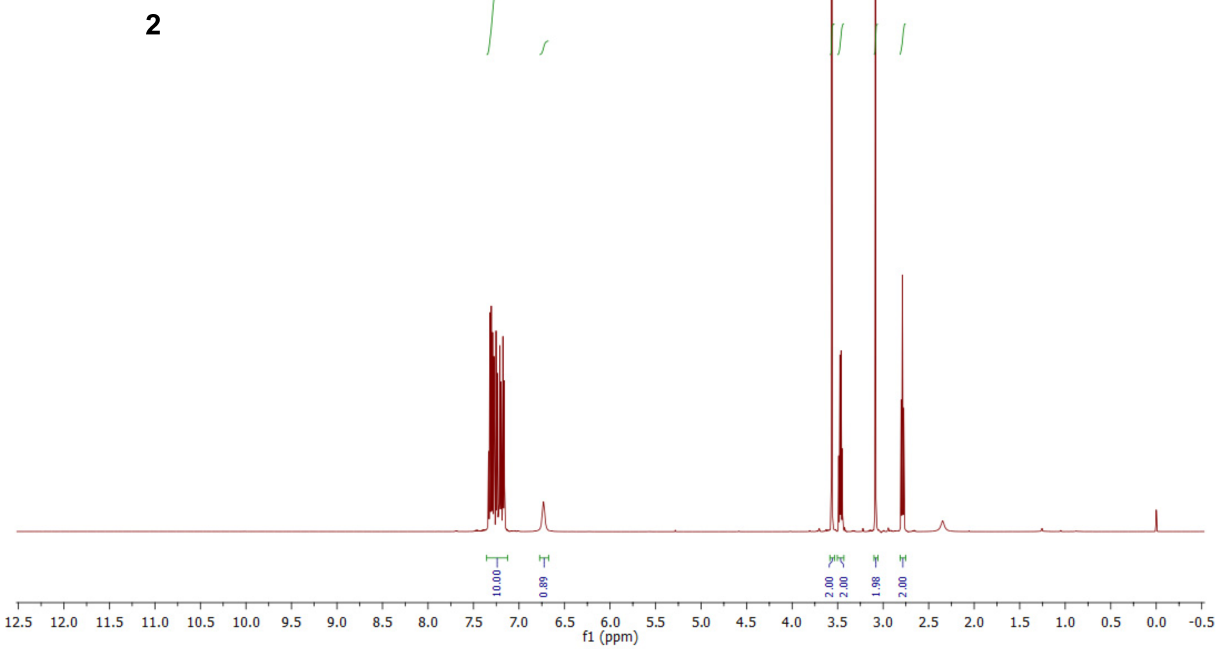


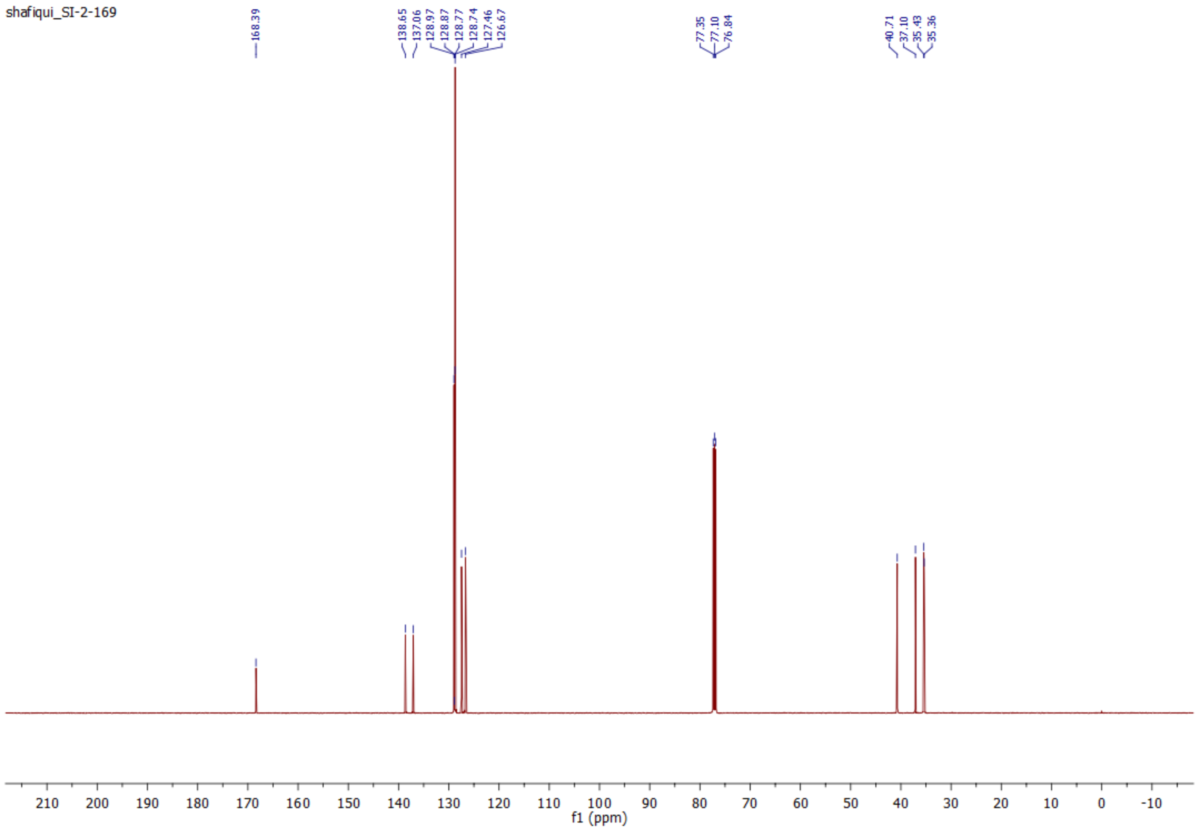

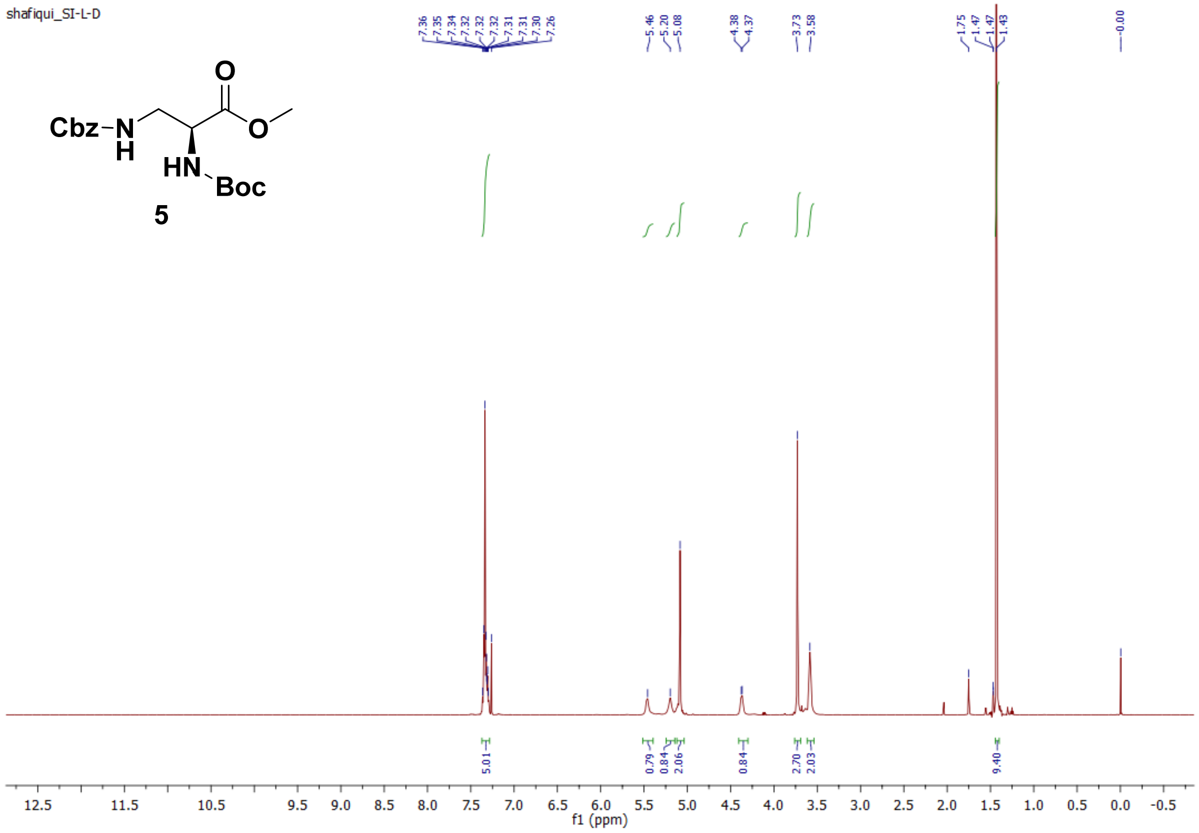
shafiqui_SI-L-D
Cbz–N
H
O
O
HN
Boc
5
7.36
7.35
7.34
7.32
7.32
7.32
7.31
7.31
7.30
7.26
5.46
5.20
5.08
4.38
4.37
3.73
3.58
1.75
1.47
1.47
1.43
0.00
5.01
0.79
0.84
2.06
0.84
2.70
2.03
9.40
12.5 11.5 10.5 9.5 9.0 8.5 8.0 7.5 7.0 6.5 6.0 5.5 5.0 4.5 4.0 3.5 3.0 2.5 2.0 1.5 1.0 0.5 0.0 -0.5
f1 (ppm)

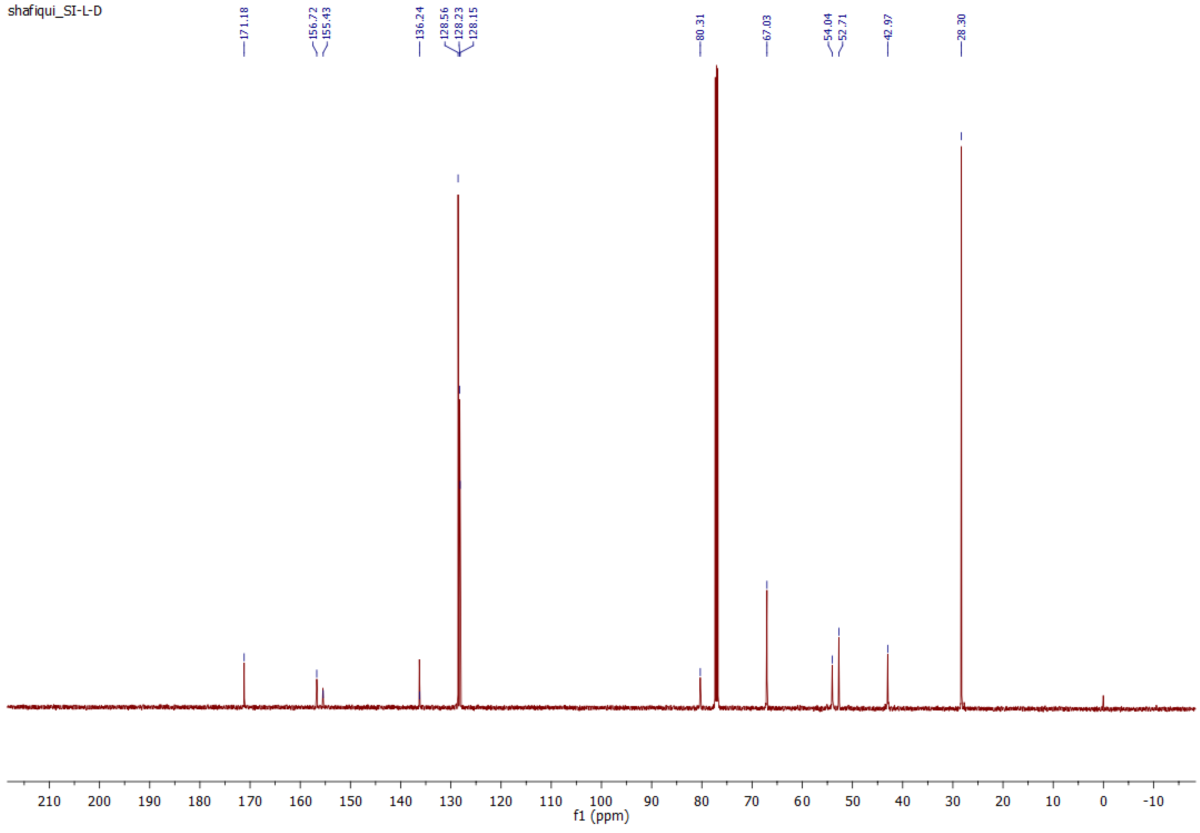
shafiqui_SI-L-D
171.18
156.72
155.43
136.24
128.56
128.23
128.15
80.31
67.03
54.04
52.71
42.97
28.30
210 200 190 180 170 160 150 140 130 120 110 100 90 80 70 60 50 40 30 20 10 0 -10
f1 (ppm)

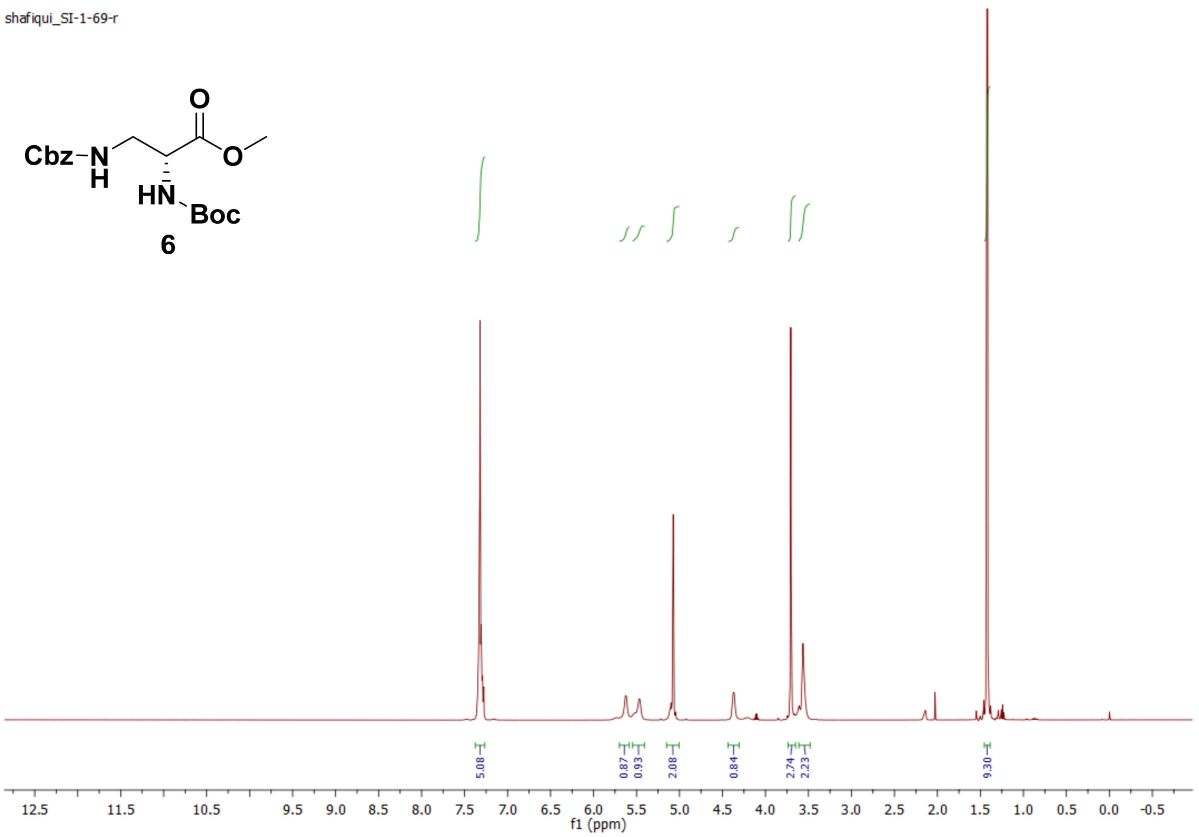
shafiqui_SI-1-69-r
Cbz
Boc
6
5.08
0.87
0.93
2.08
0.84
2.74
2.23
9.30
f1 (ppm)

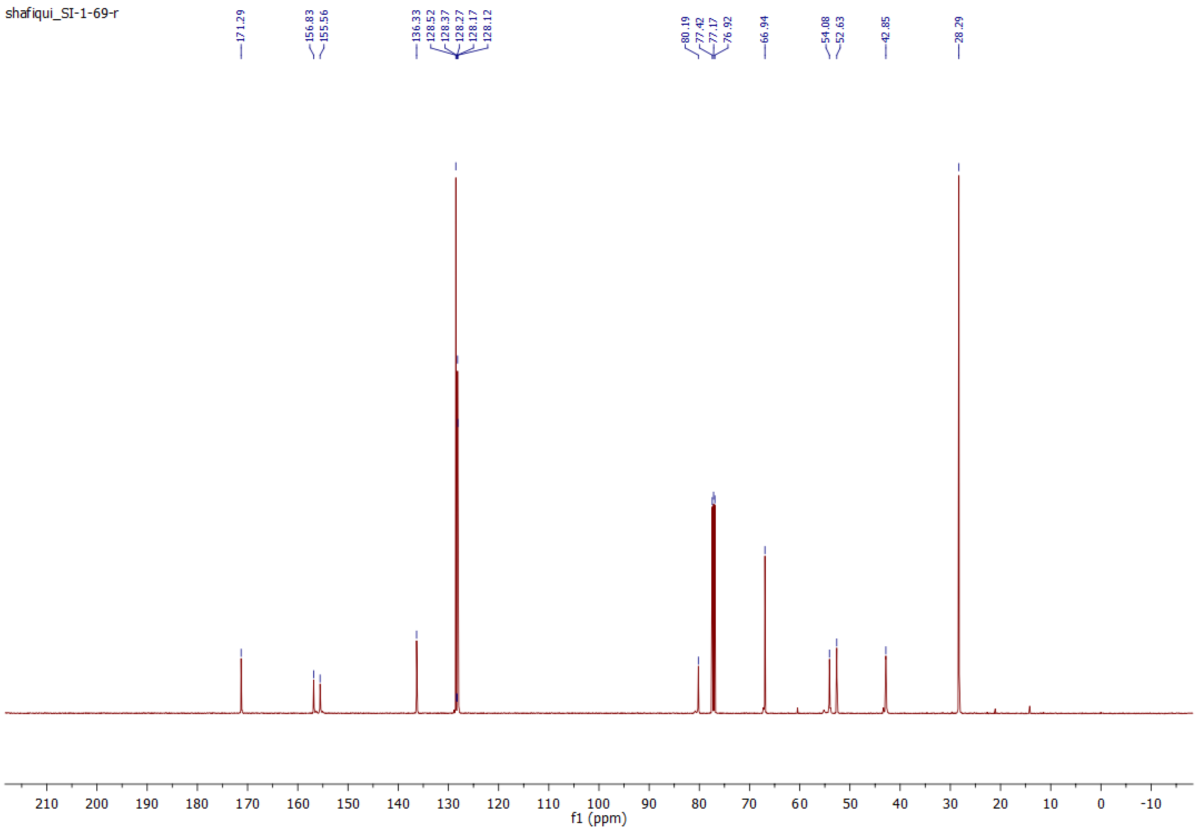
shafiqui_SI-1-69-r
171.29
156.83
155.56
136.33
128.52
128.37
128.27
128.17
128.12
80.19
77.42
77.17
76.92
66.94
54.08
52.63
42.85
28.29
f1 (ppm)

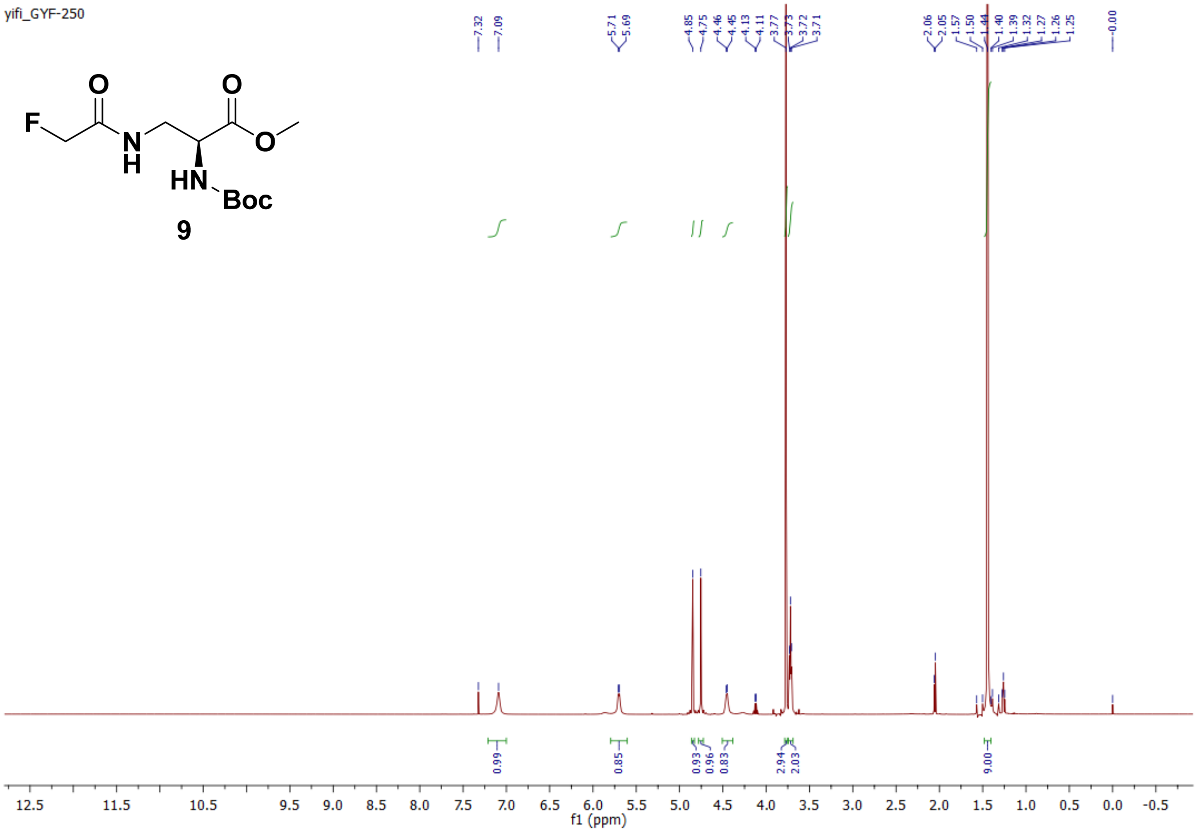
yifi_GYF-250
9
f1 (ppm)

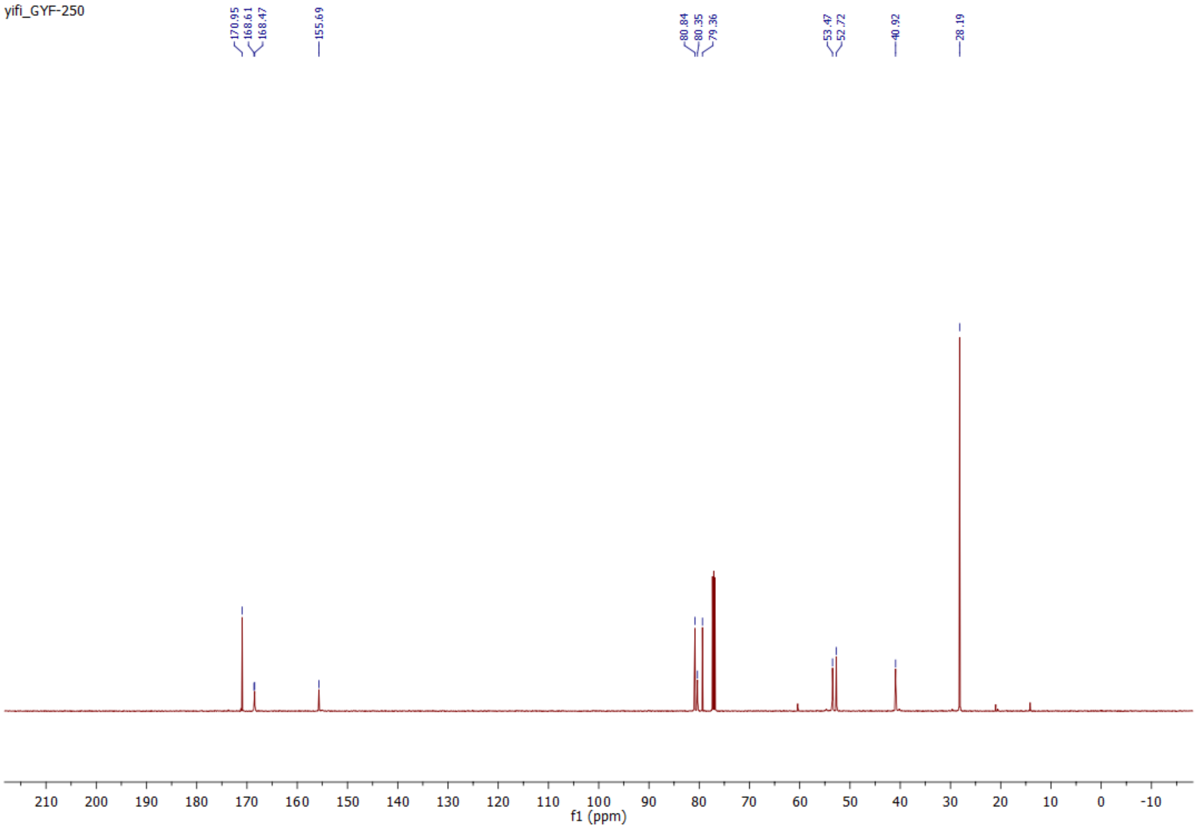
yifi_GYF-250
f1 (ppm)

shafiqui_SI-D-F

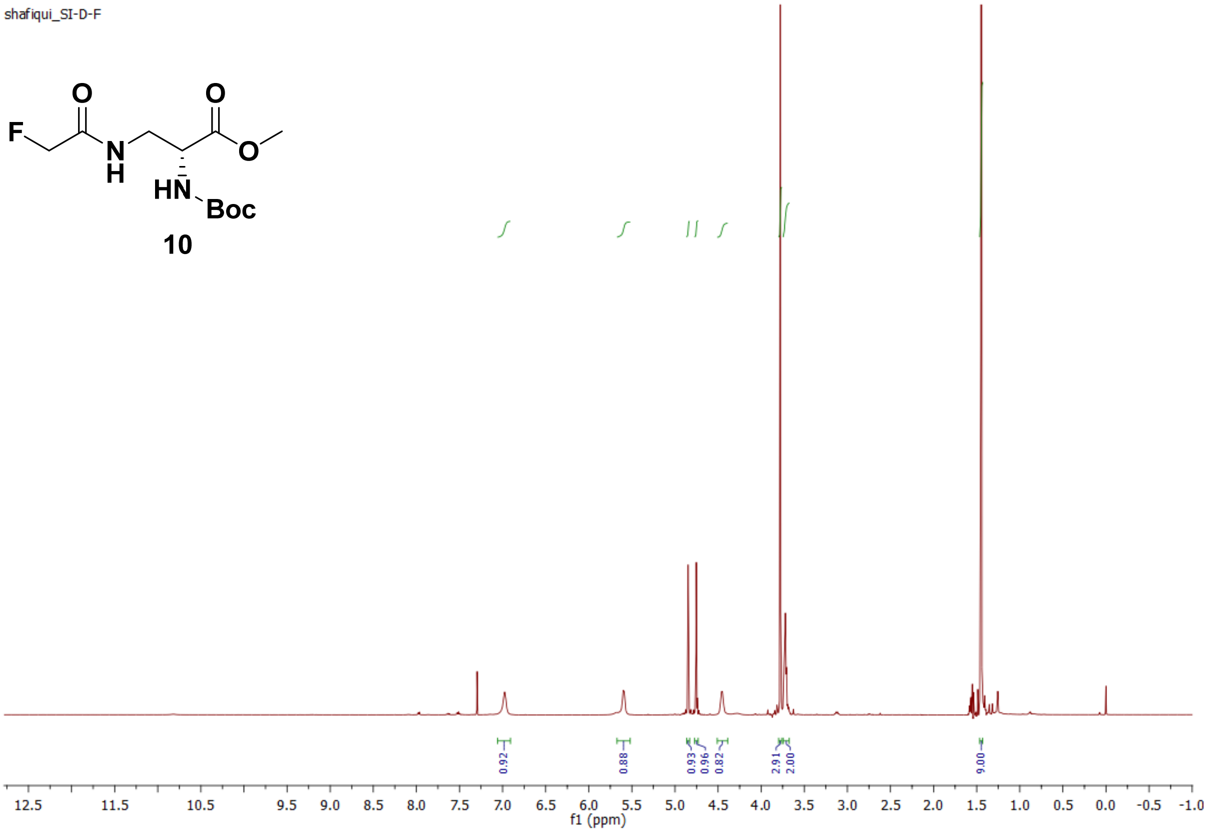


shafiqui_SI-D-F

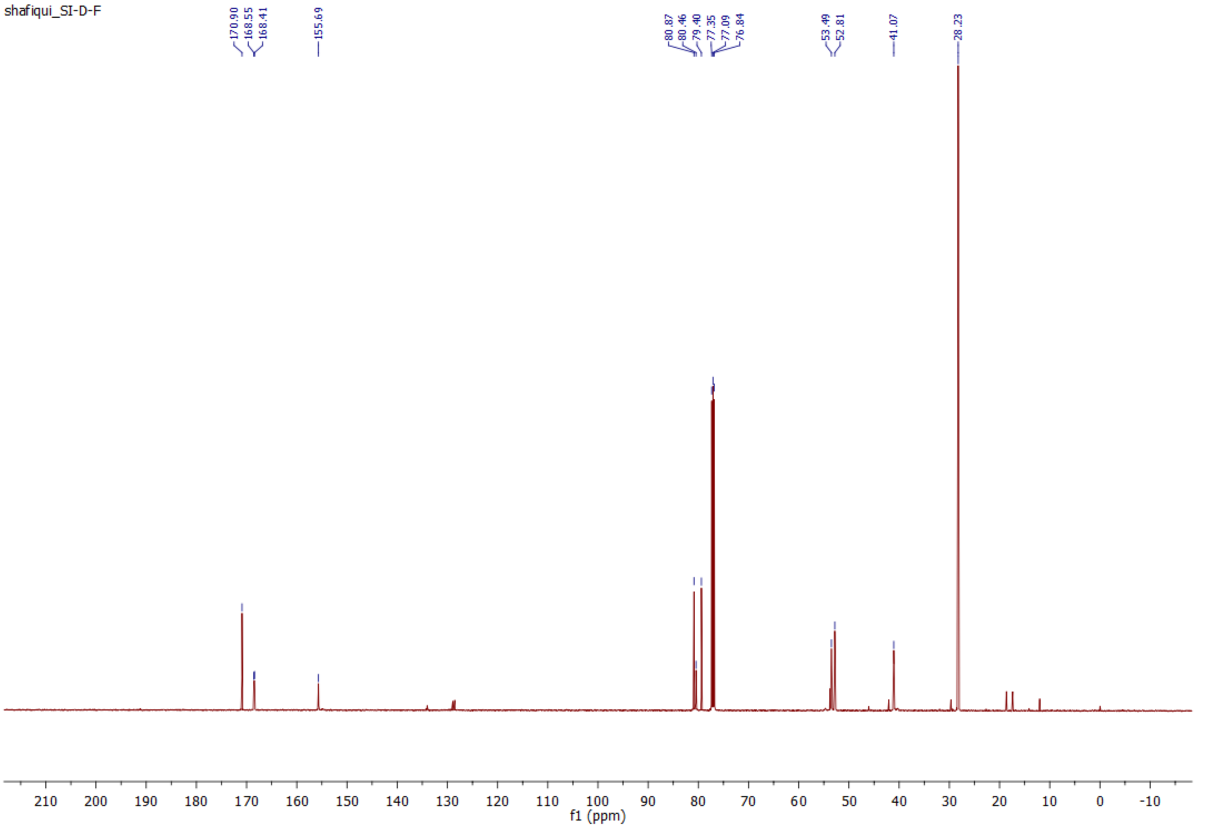

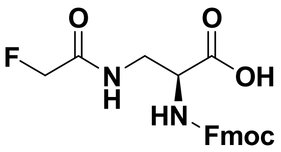
F
O
N
H
O
OH
HN
Fmoc

**15**

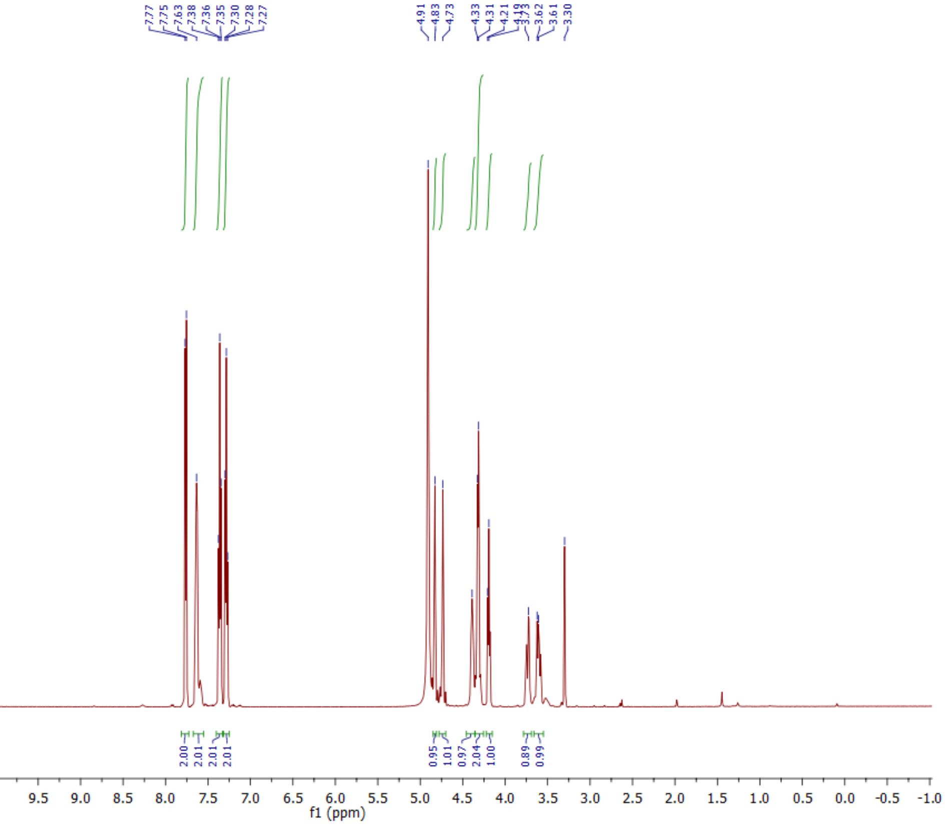
shafiqui_SI-L
7.77
7.75
7.63
7.38
7.36
7.35
7.30
7.28
7.27
4.91
4.83
4.73
4.33
4.31
4.21
4.19
3.73
3.62
3.61
3.30
2.00
2.01
2.01
2.01
0.95
1.01
0.97
2.04
1.00
0.89
0.99
f1 (ppm)

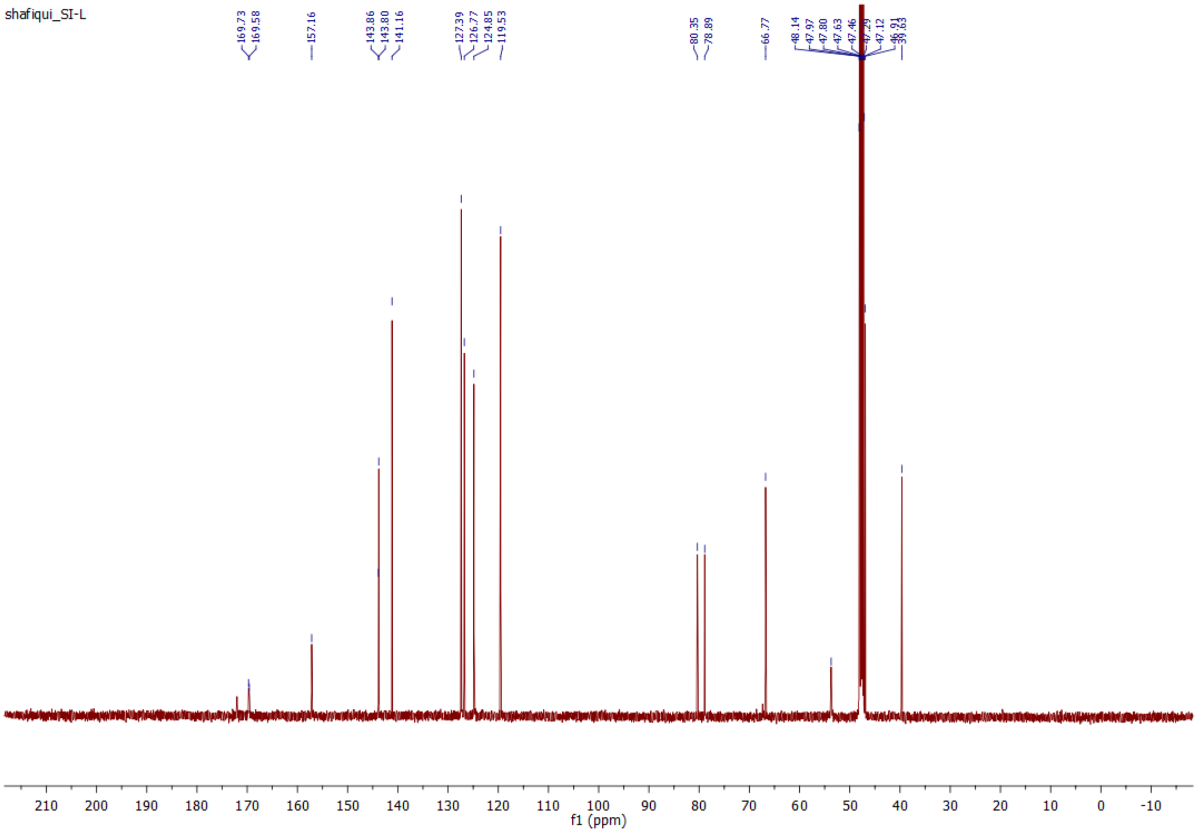
shafiqui_SI-L
169.73
169.58
157.16
143.86
143.80
141.16
127.39
126.77
124.85
119.53
80.35
78.89
66.77
48.14
47.97
47.80
47.63
47.46
47.29
47.12
46.91
39.63
f1 (ppm)

shafiqui_SI-1-94

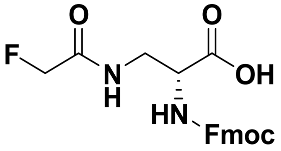


**16**

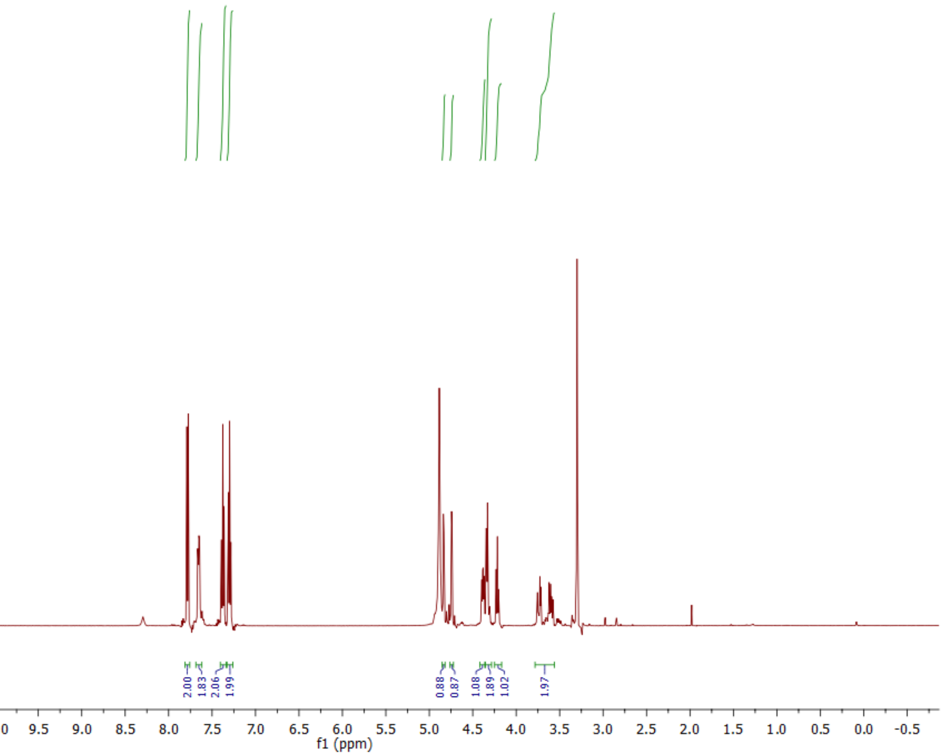


shafiqui_SI-1-94

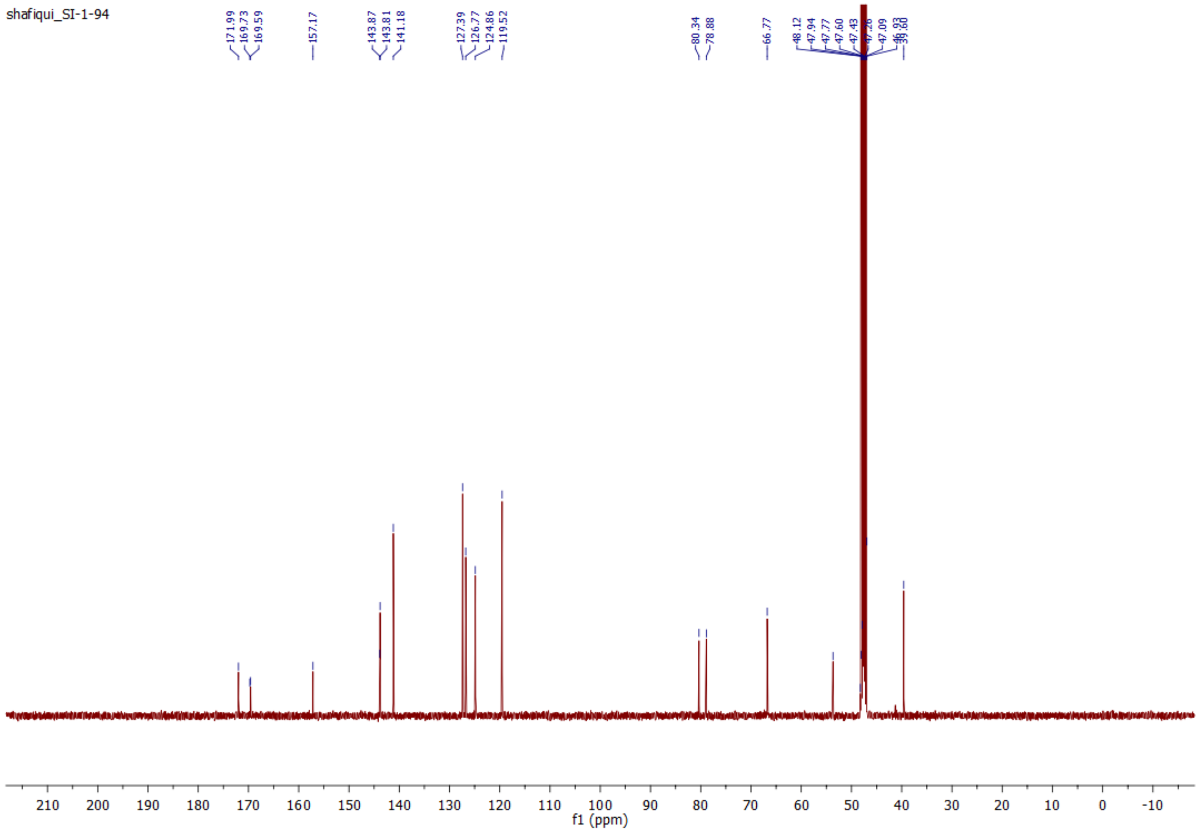

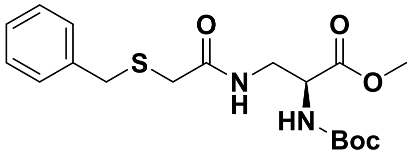
O
O
S
N
H
O
HN
Boc

**17**

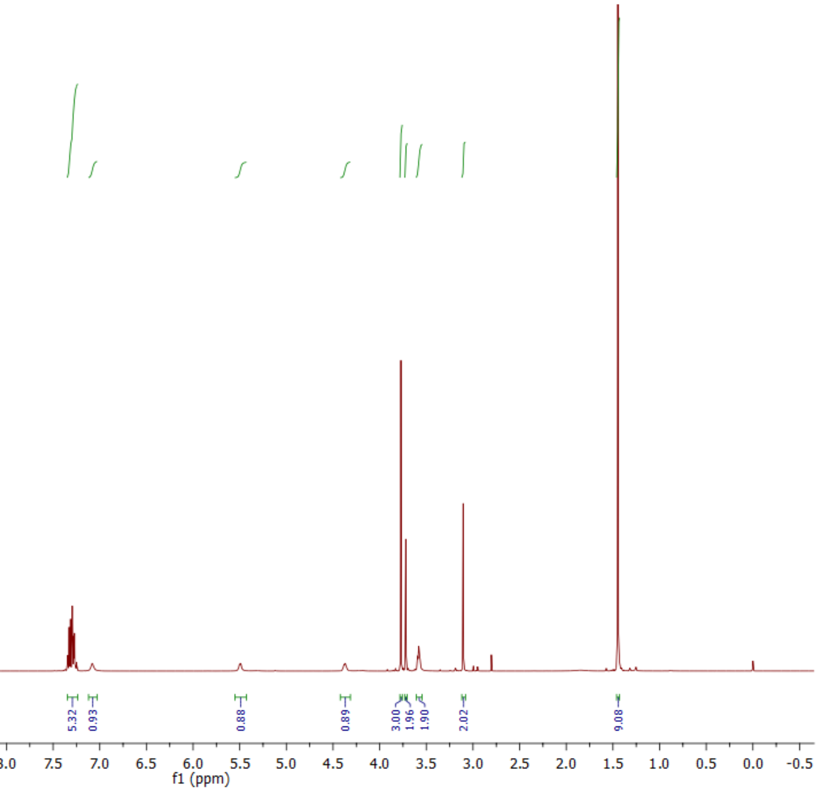
5.32
0.93
0.88
0.89
3.00
1.96
1.90
2.02
9.08
f1 (ppm)

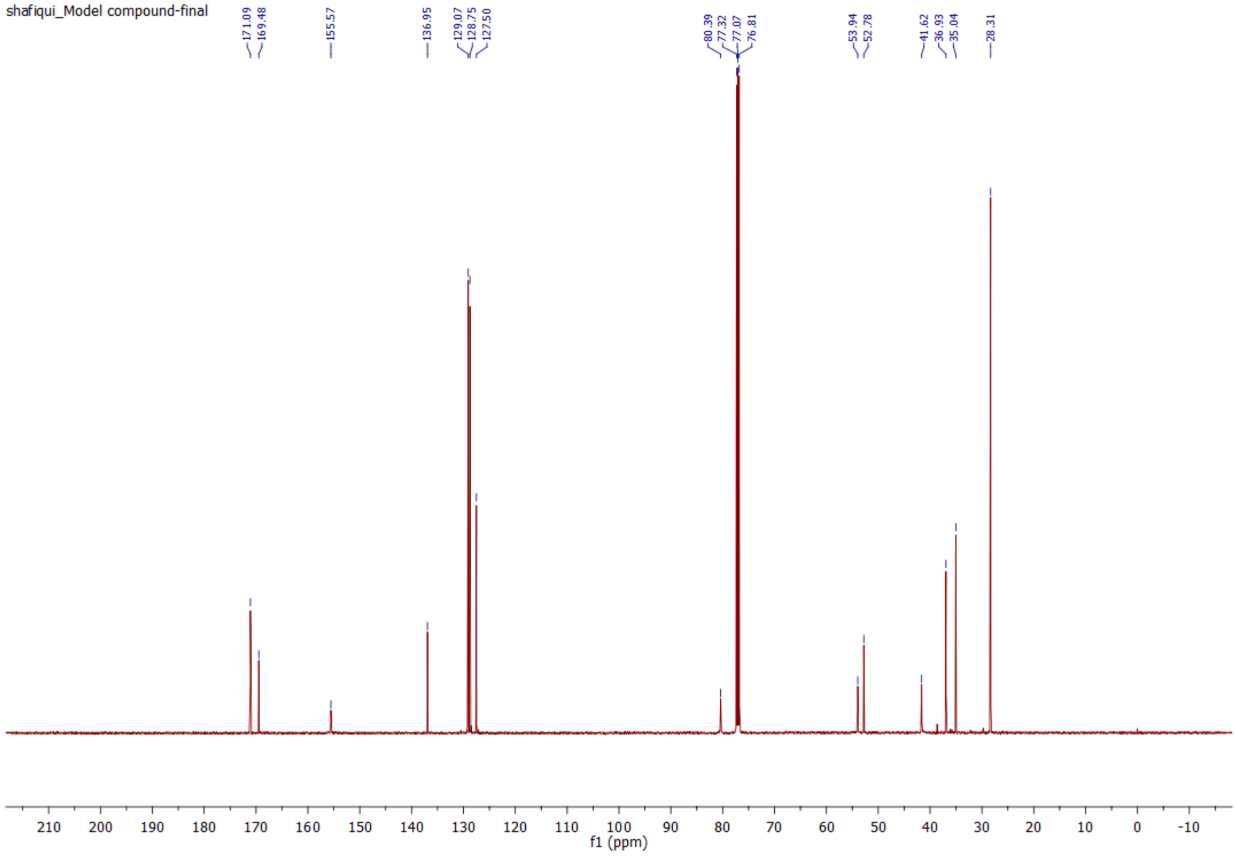
shafiqui_Model compound-final
171.09
169.48
155.57
136.95
129.07
128.75
127.50
80.39
77.32
77.07
76.81
53.94
52.78
41.62
36.93
35.04
28.31
f1 (ppm)

### LC-Chromatogram and MS-Spectra for Peptides

Peptide **21**.

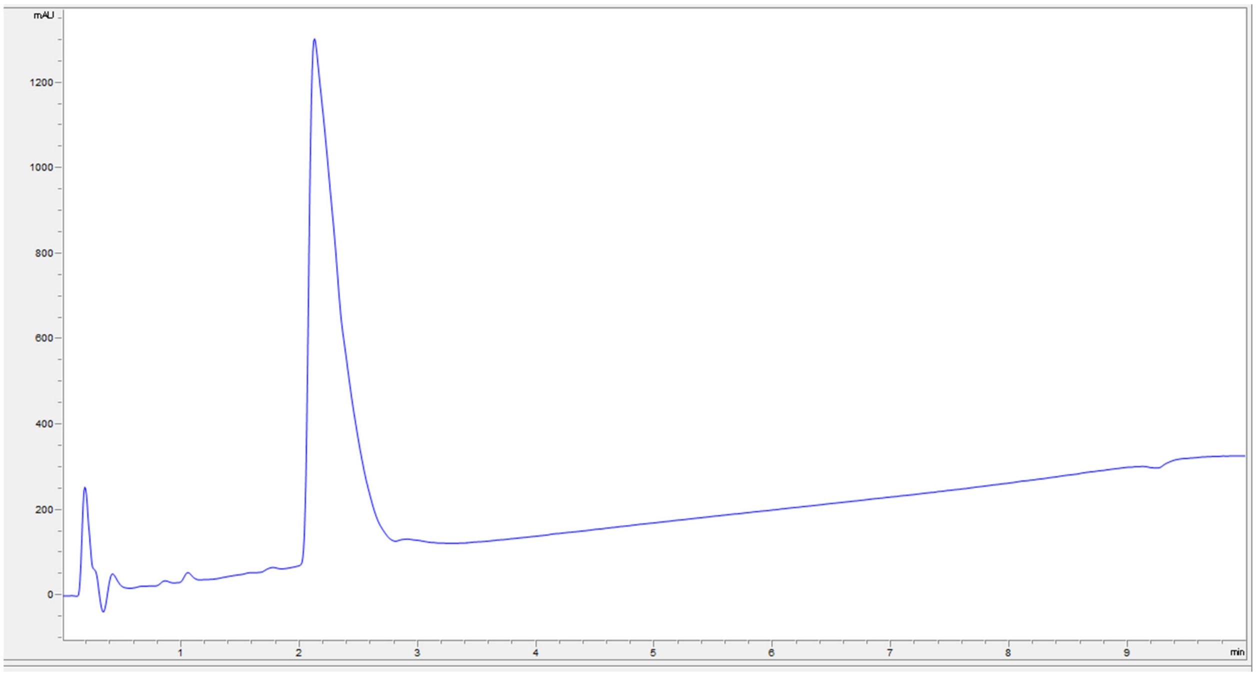


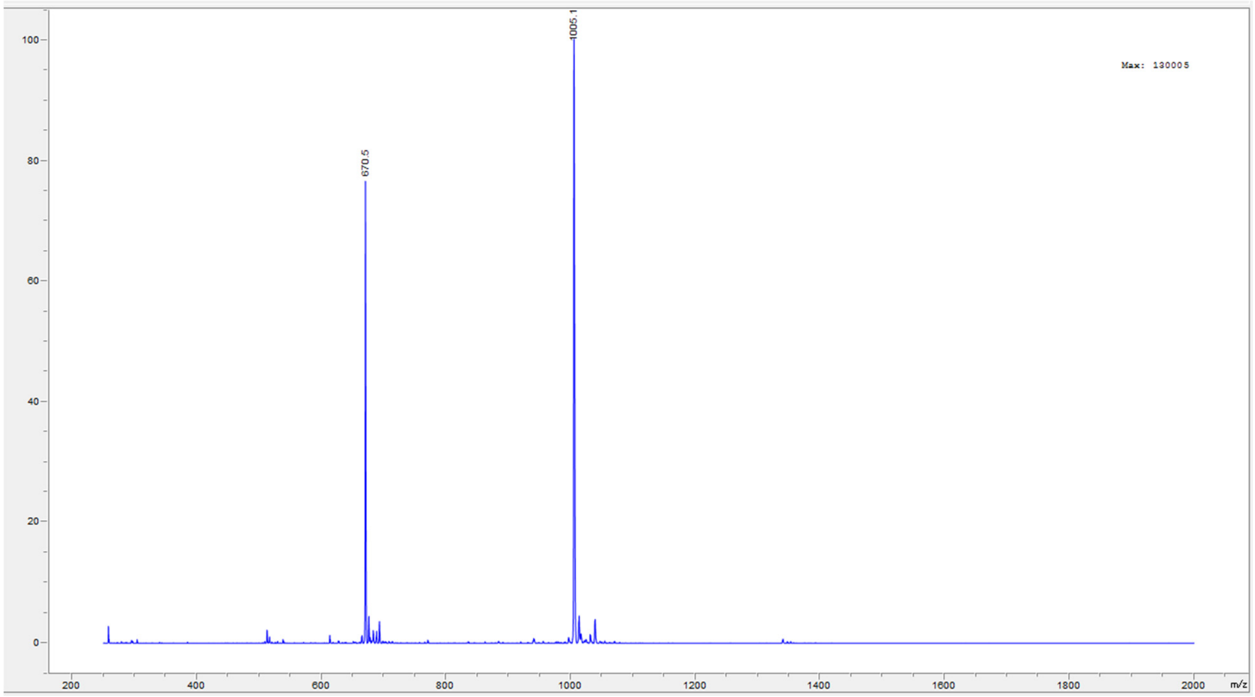

Peptide **22**.

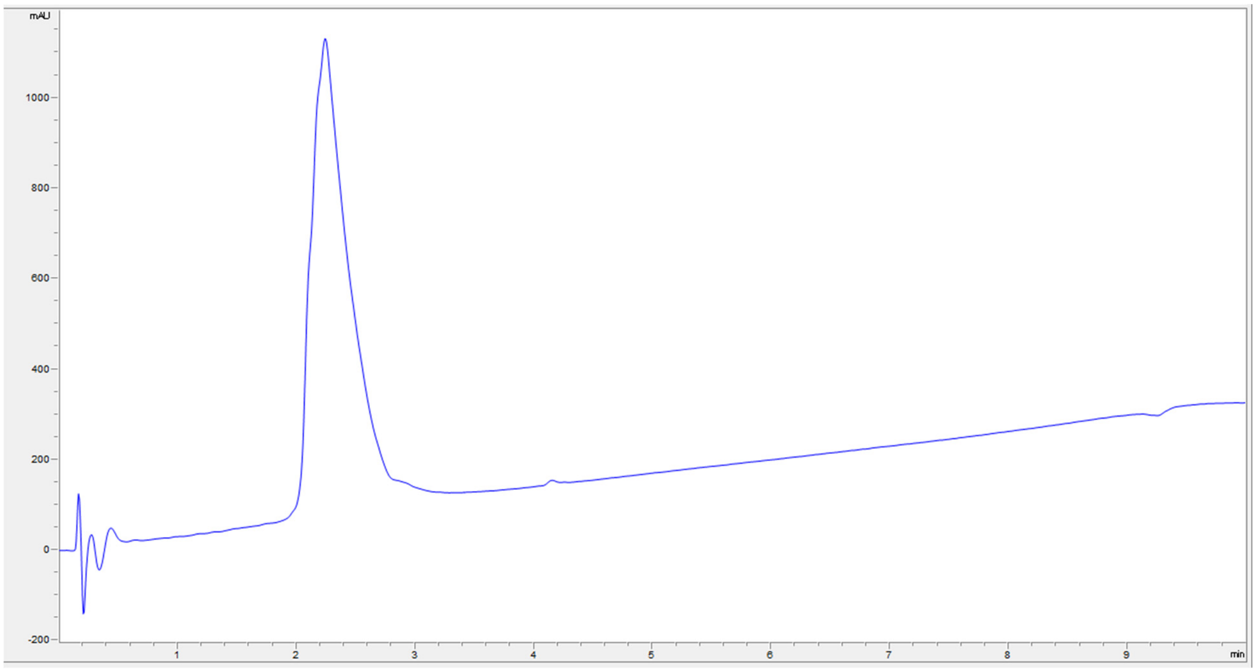
mAU
1000
800
600
400
200
0
-200
1
2
3
4
5
6
7
8
9
min

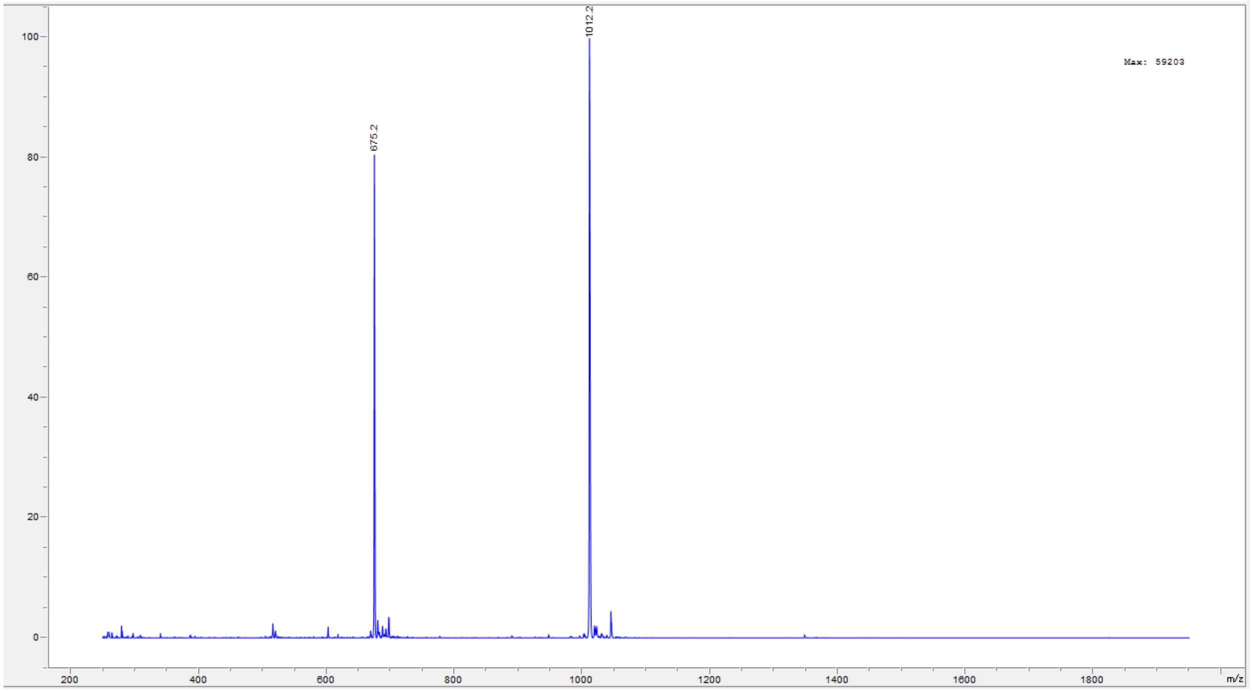
1012.2
675.2
Max: 59203
100
80
60
40
20
0
200
400
600
800
1000
1200
1400
1600
1800
m/z

Peptide **23**.

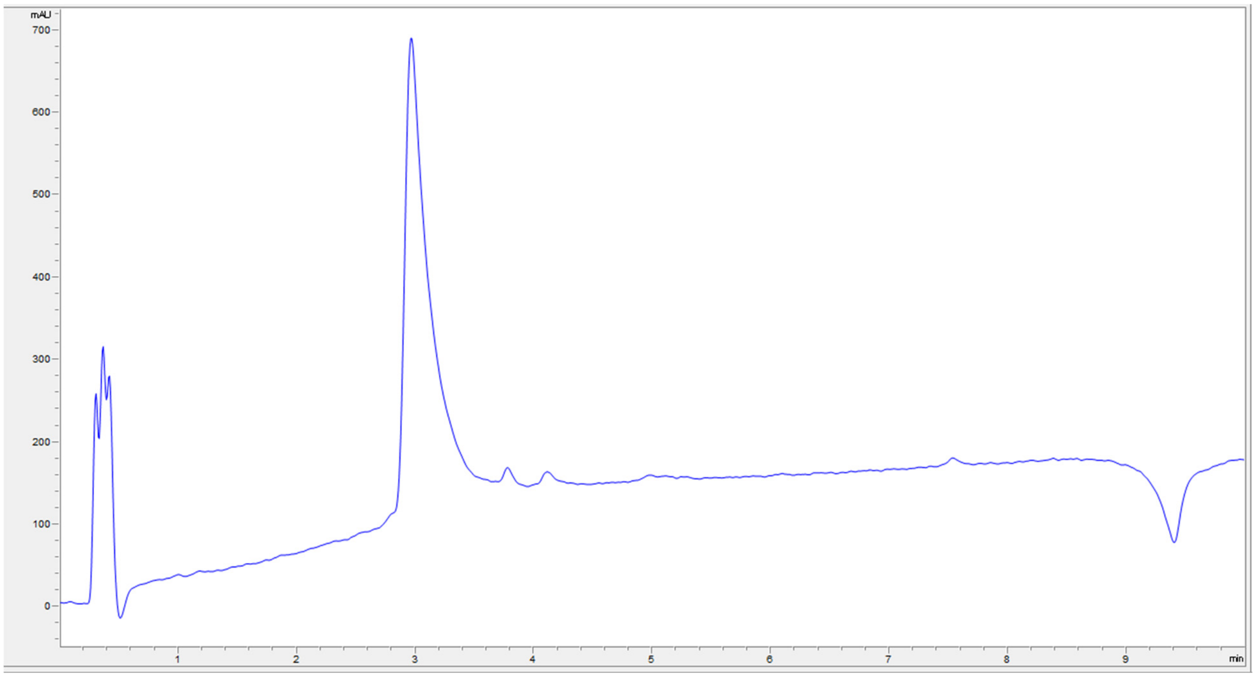

mAU
700
600
500
400
300
200
100
0
1
2
3
4
5
6
7
8
9
min


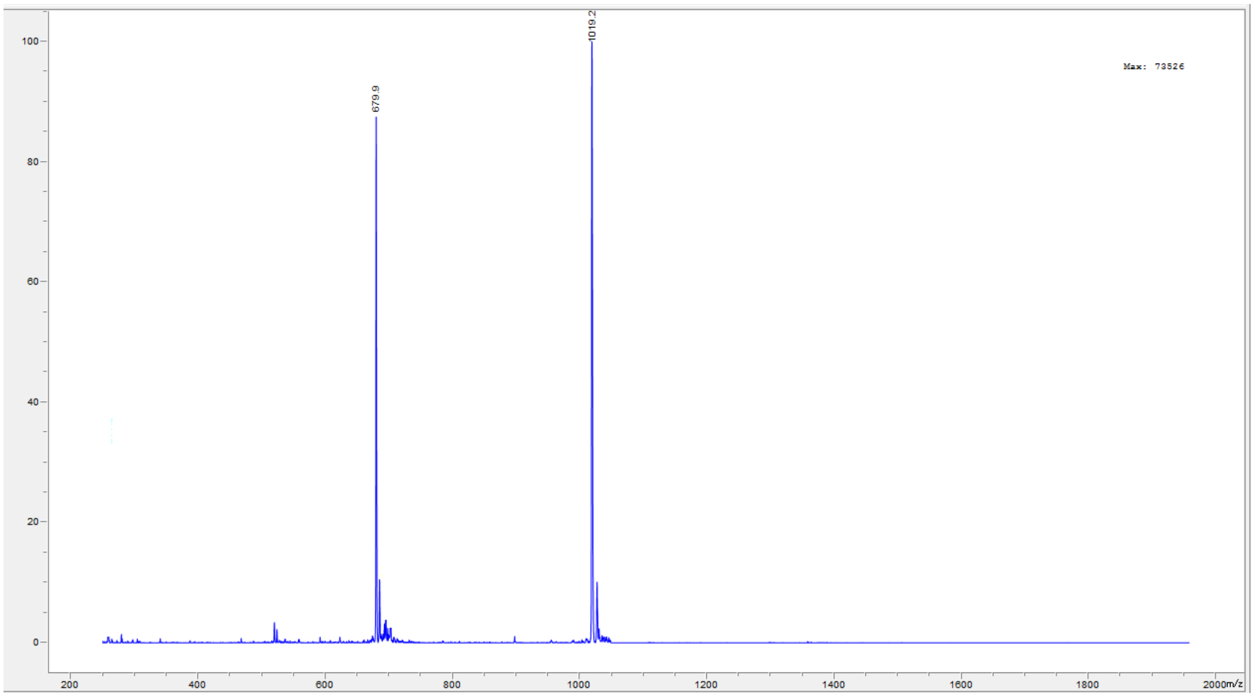

1019.2
679.9
Max: 73526
100
80
60
40
20
0
200
400
600
800
1000
1200
1400
1600
1800
2000m/z

Peptide **24**.

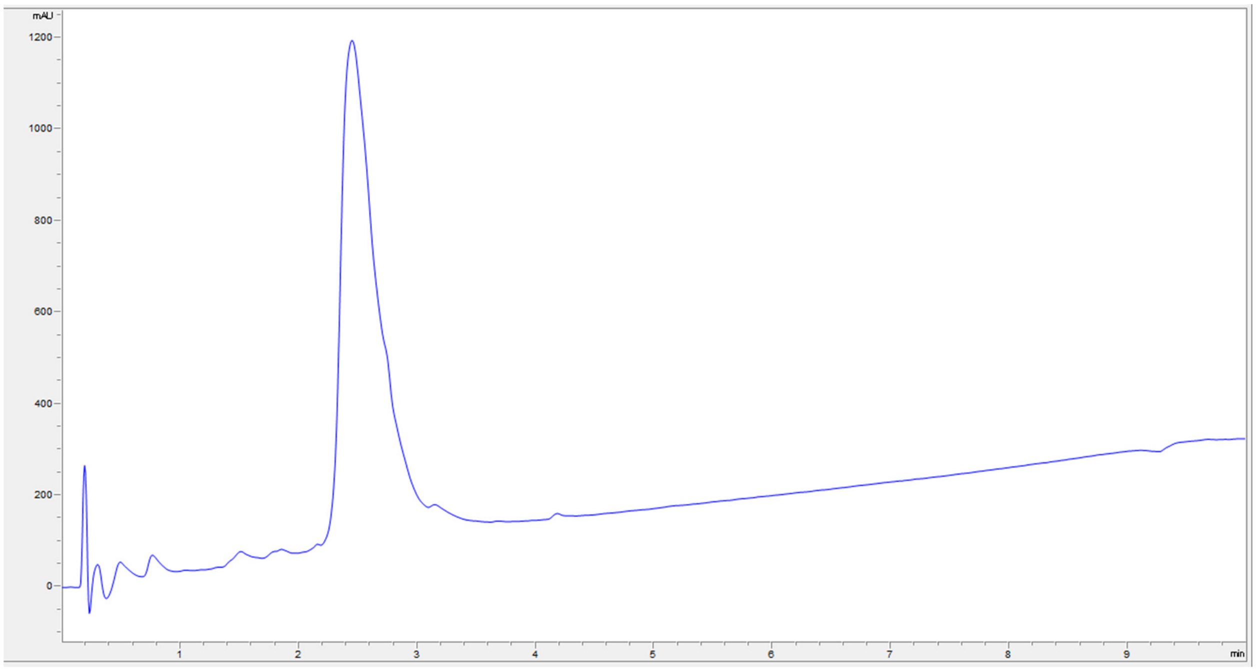

mAU
1200
1000
800
600
400
200
0
1
2
3
4
5
6
7
8
9
min


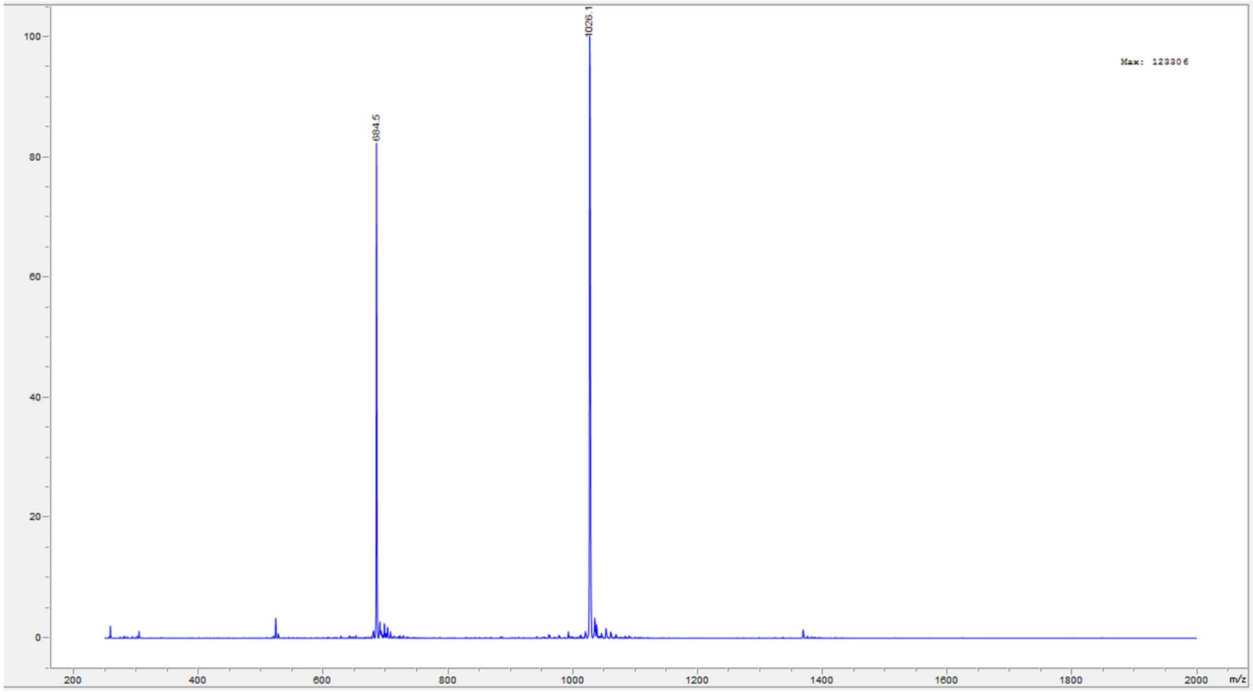

1026.1
684.5
Max: 123306
100
80
60
40
20
0
200
400
600
800
1000
1200
1400
1600
1800
2000
m/z

Peptide **25**.

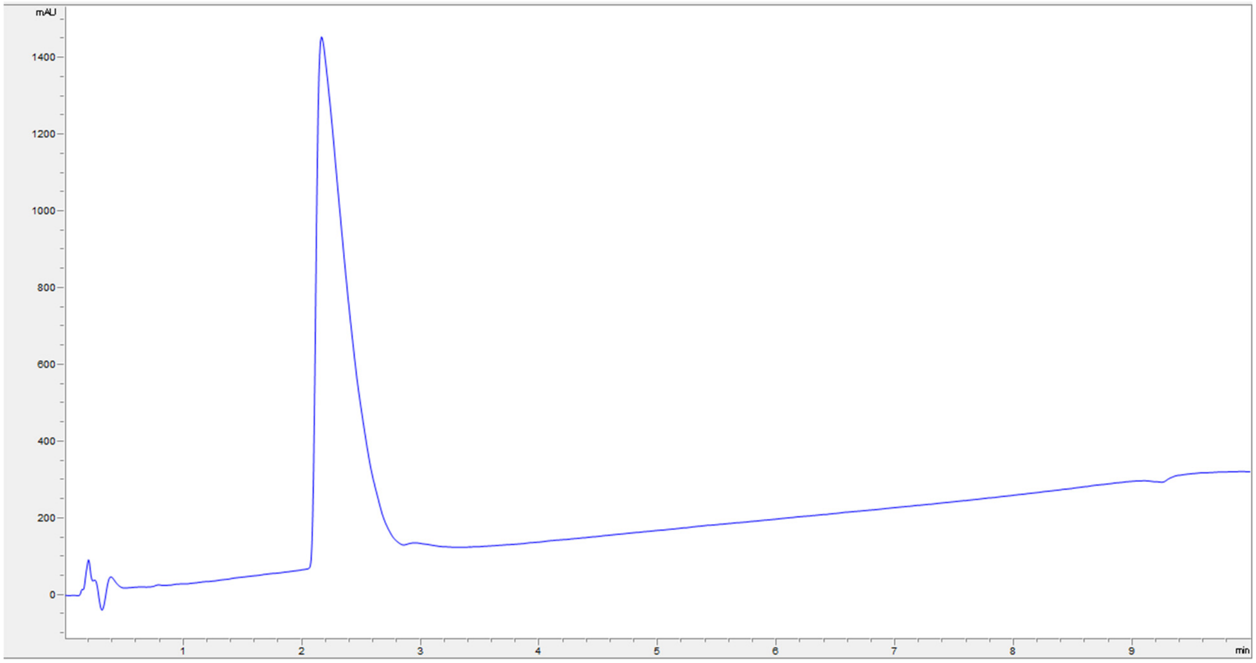
mAU
1400
1200
1000
800
600
400
200
0
1
2
3
4
5
6
7
8
9
min

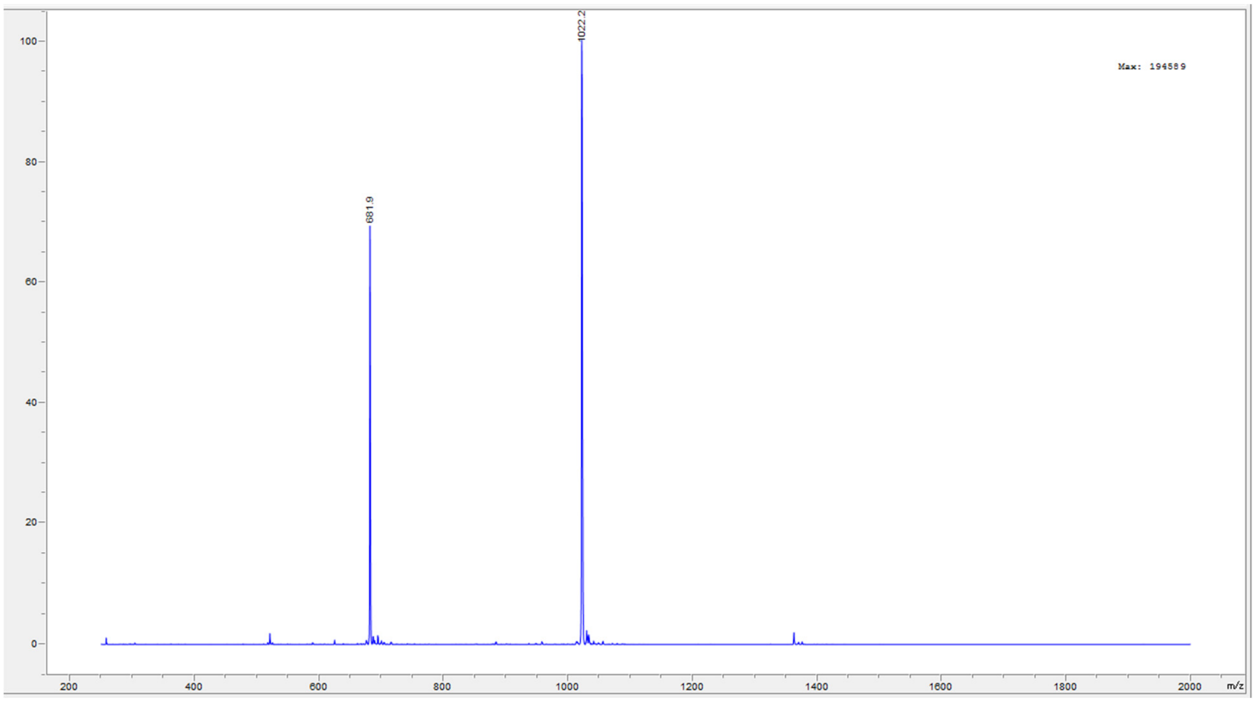
1022.2
681.9
Max: 194589
100
80
60
40
20
0
200
400
600
800
1000
1200
1400
1600
1800
2000
m/z

Peptide **26**.

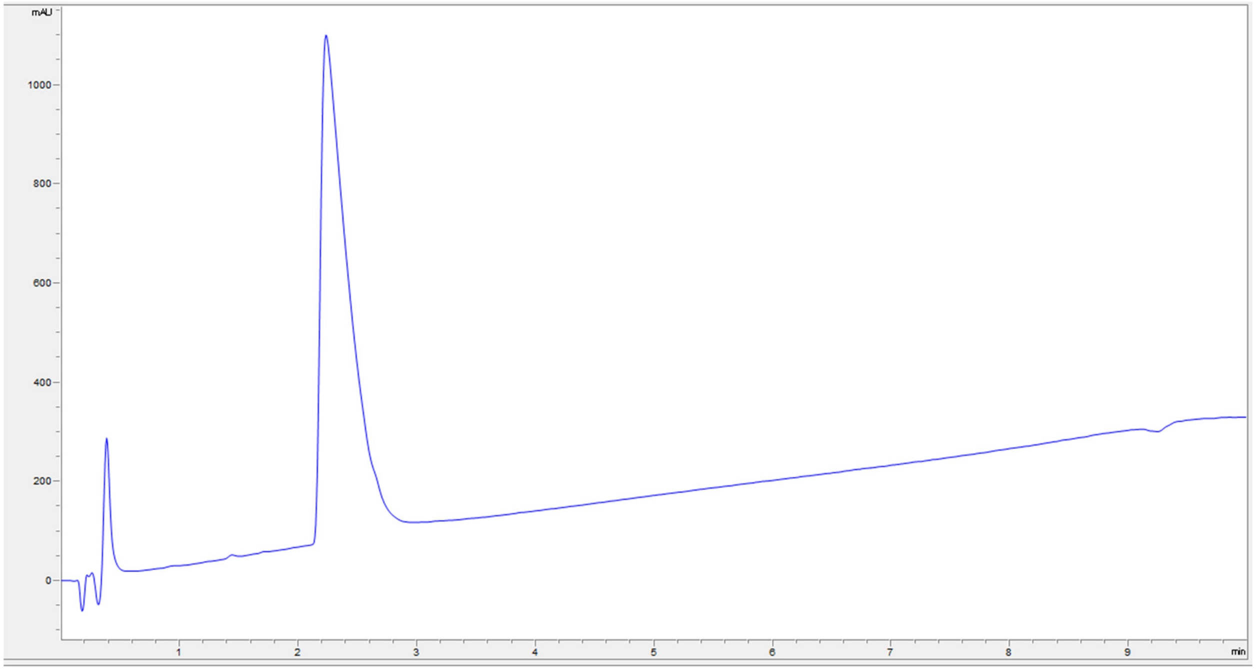

mAU
1000
800
600
400
200
0
1
2
3
4
5
6
7
8
9
min


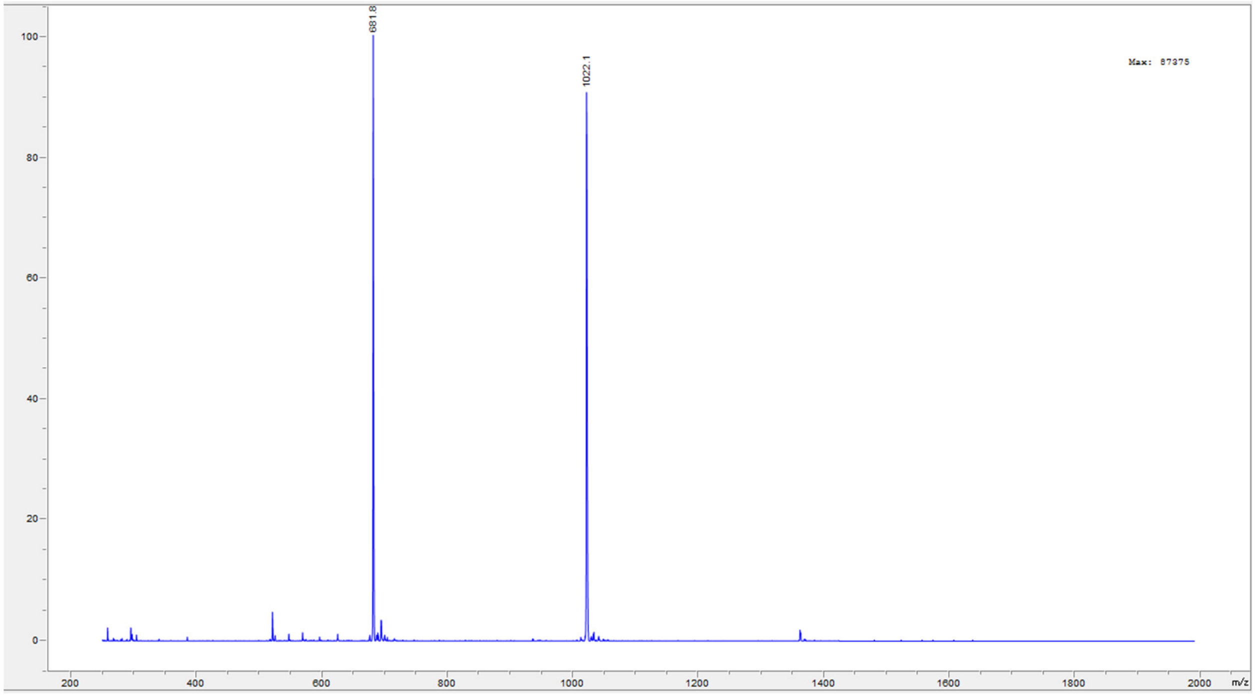

681.8
1022.1
Max: 87375
100
80
60
40
20
0
200
400
600
800
1000
1200
1400
1600
1800
2000
m/z

Peptide **28**.

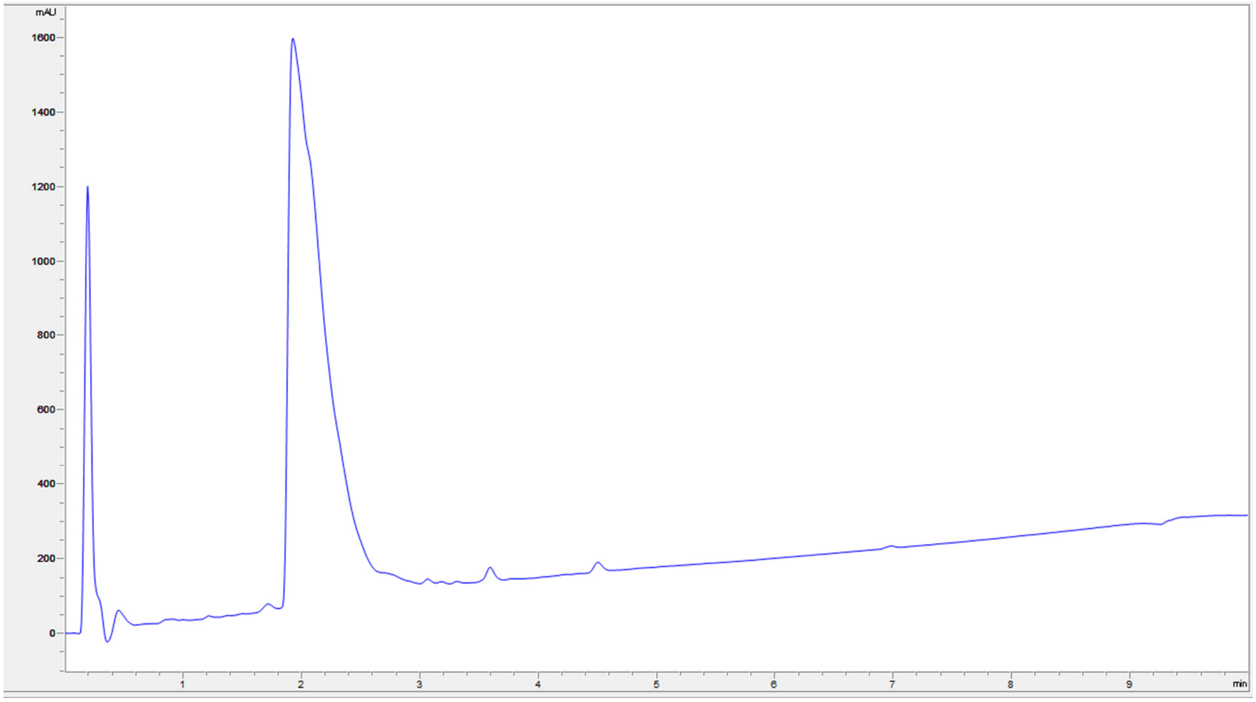

mAU
1600
1400
1200
1000
800
600
400
200
0
1
2
3
4
5
6
7
8
9
min


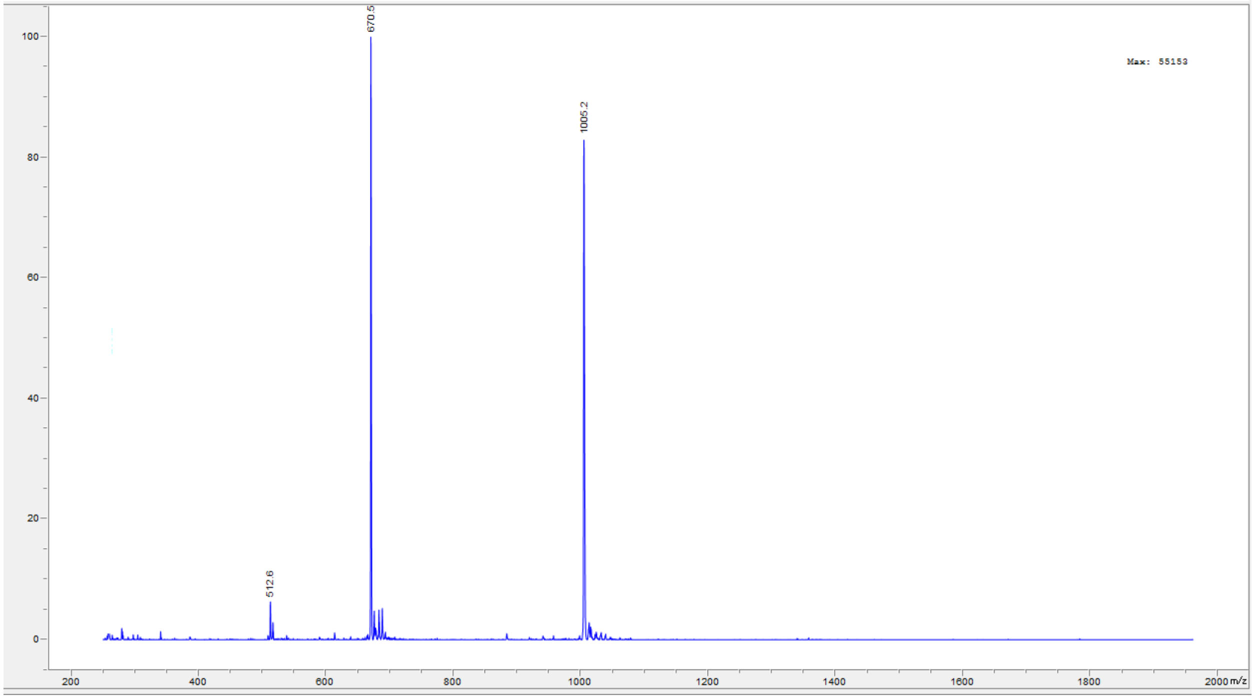

670.5
1005.2
512.6
Max: 55153
100
80
60
40
20
0
200
400
600
800
1000
1200
1400
1600
1800
2000 m/z

Peptide **29**.

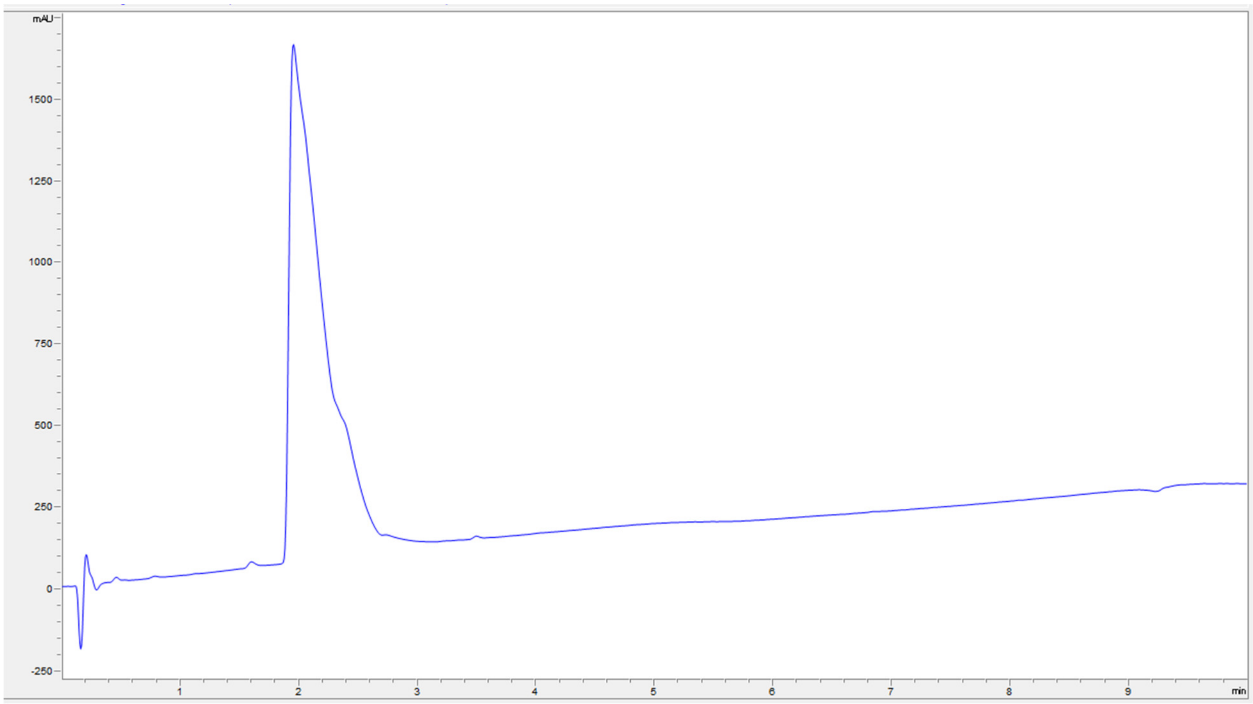

mAU
1500
1250
1000
750
500
250
0
-250
1
2
3
4
5
6
7
8
9
min


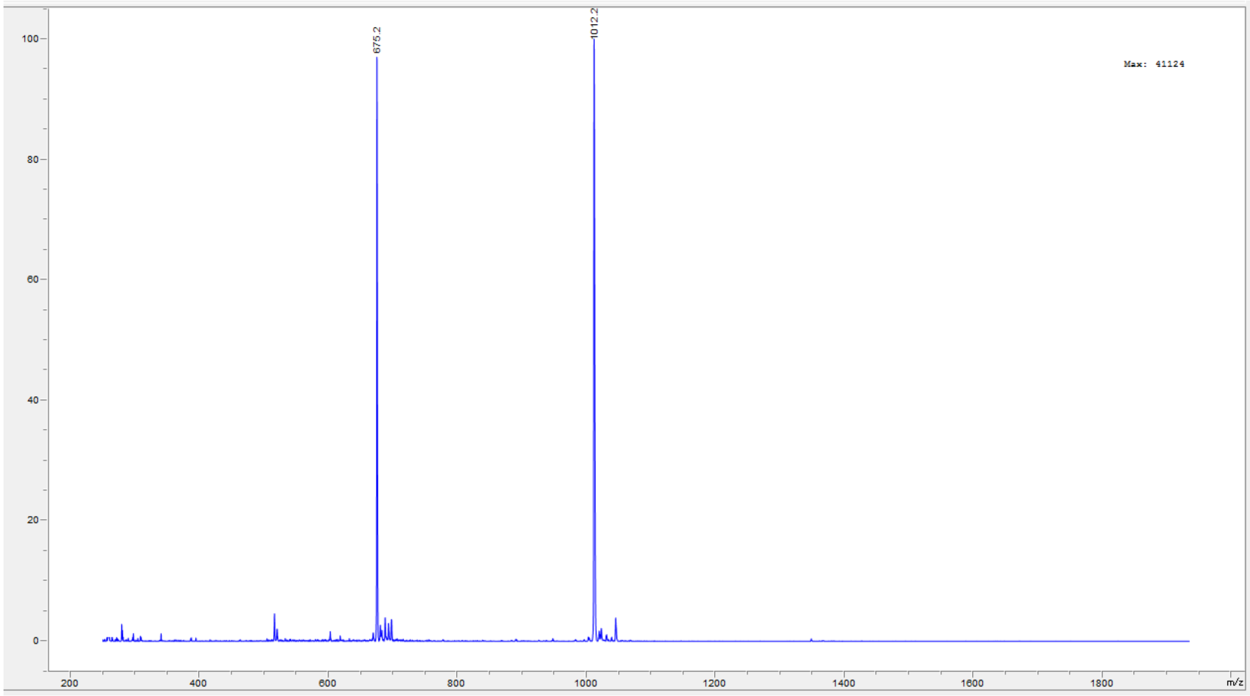

675.2
1012.2
Max: 41124
100
80
60
40
20
0
200
400
600
800
1000
1200
1400
1600
1800
m/z

Peptide **30**.

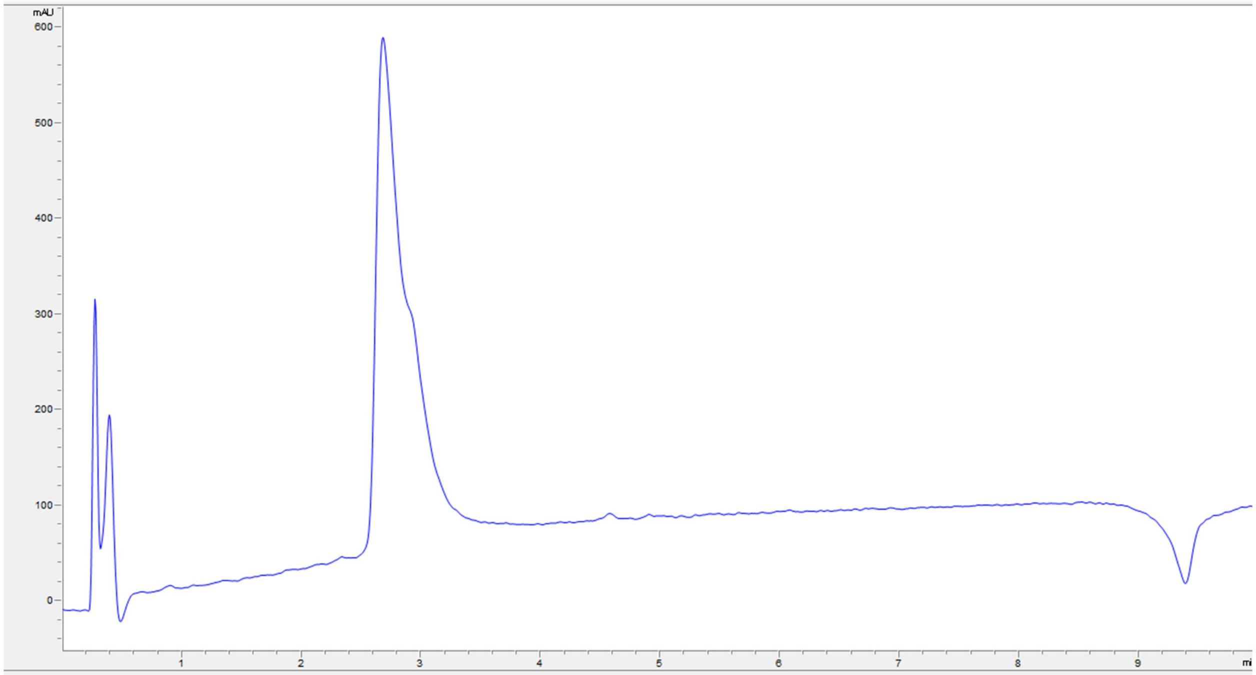
mAU
600
500
400
300
200
100
0
1
2
3
4
5
6
7
8
9

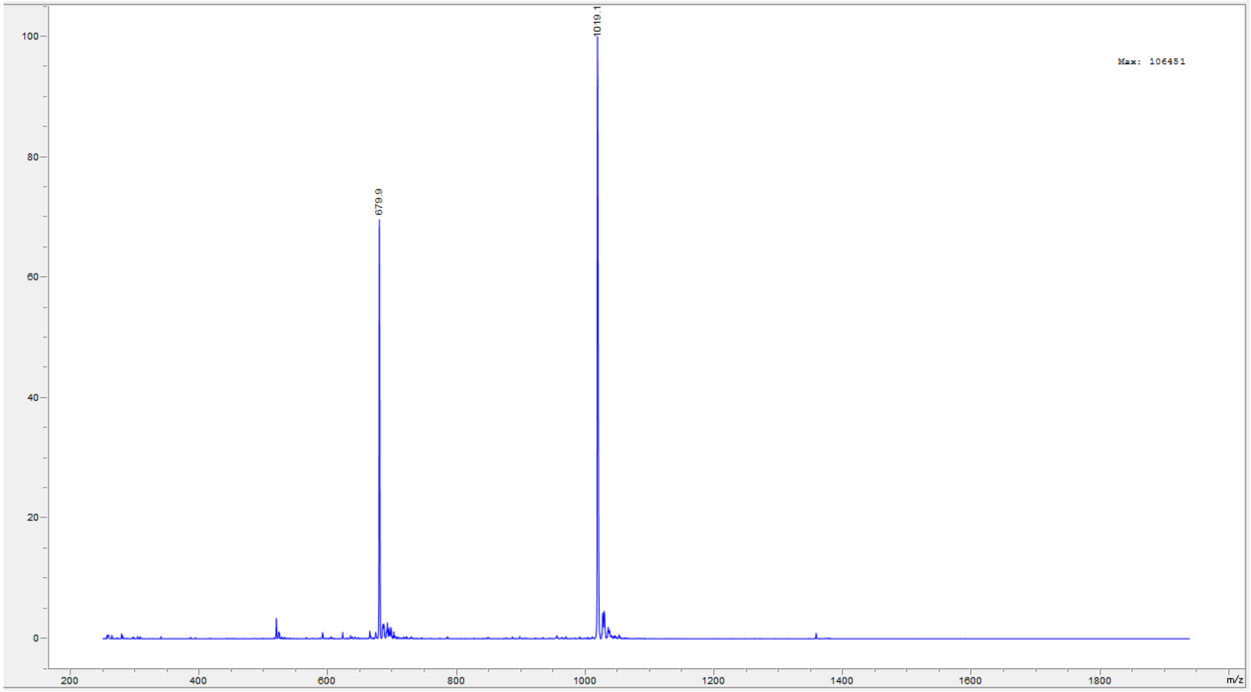
1019.1
679.9
Max: 106451
100
80
60
40
20
0
200
400
600
800
1000
1200
1400
1600
1800
m/z

Peptide **31**.

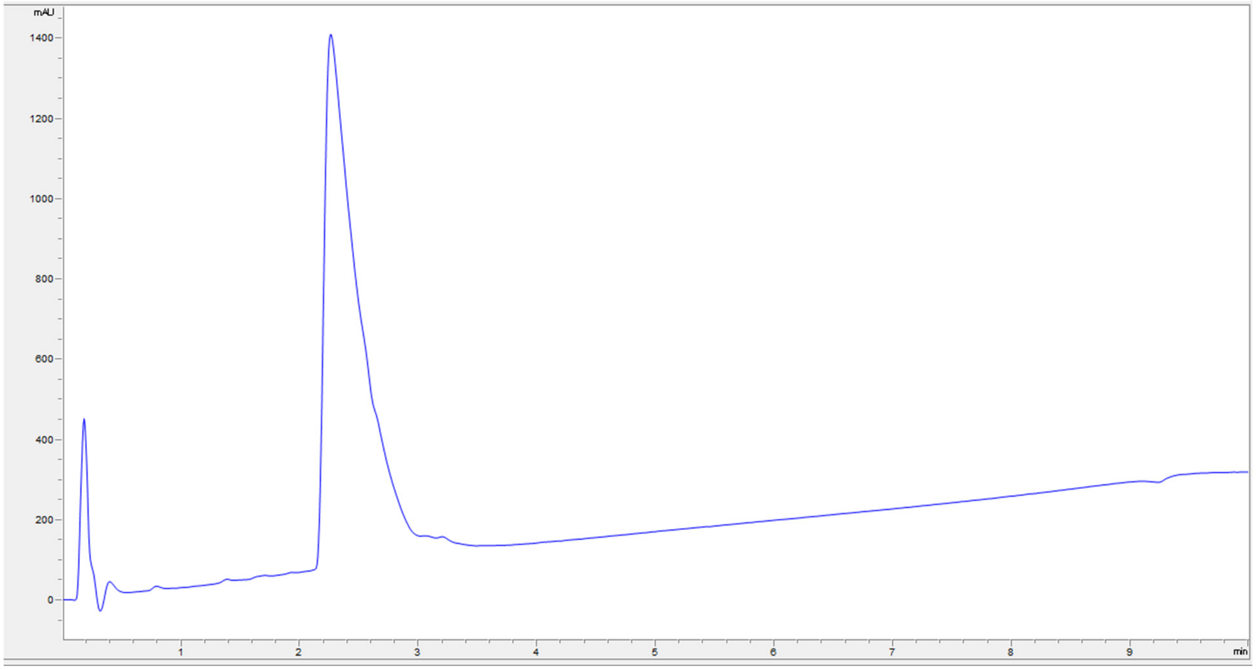
mAU
1400
1200
1000
800
600
400
200
0
1
2
3
4
5
6
7
8
9
min

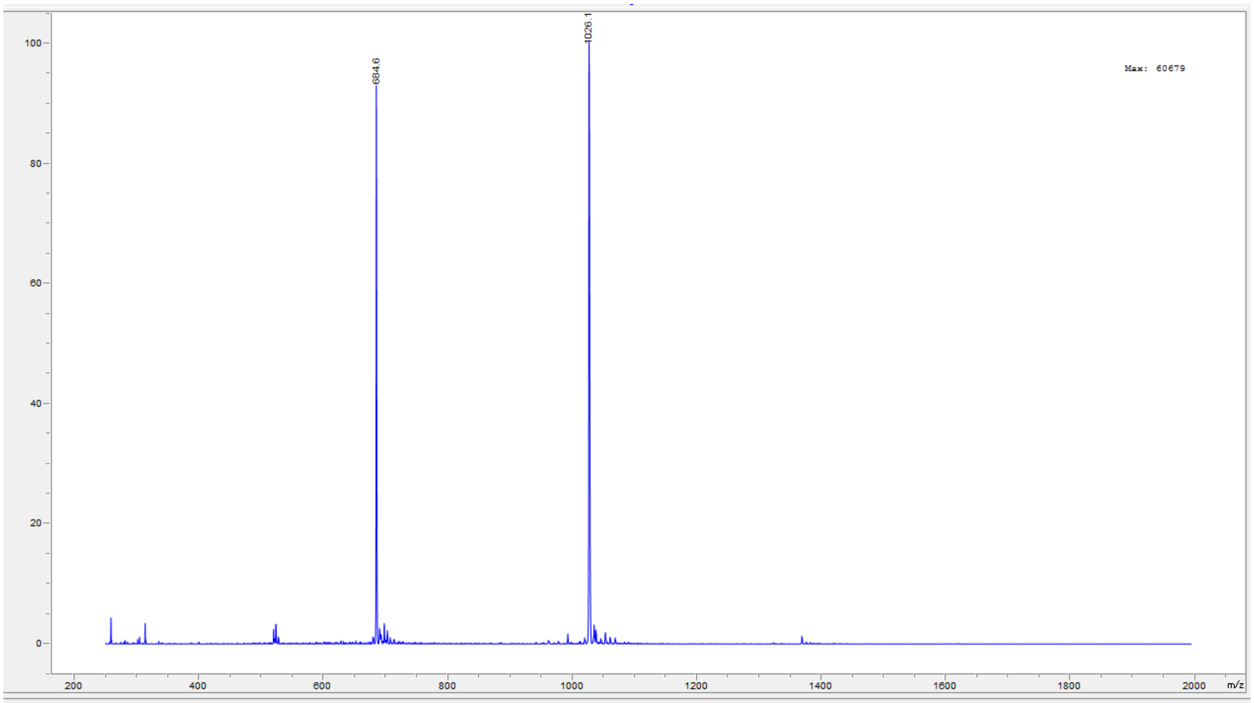
100
80
60
40
20
0
684.6
1026.1
Max: 60679
200
400
600
800
1000
1200
1400
1600
1800
2000
m/z

Peptide **32**.

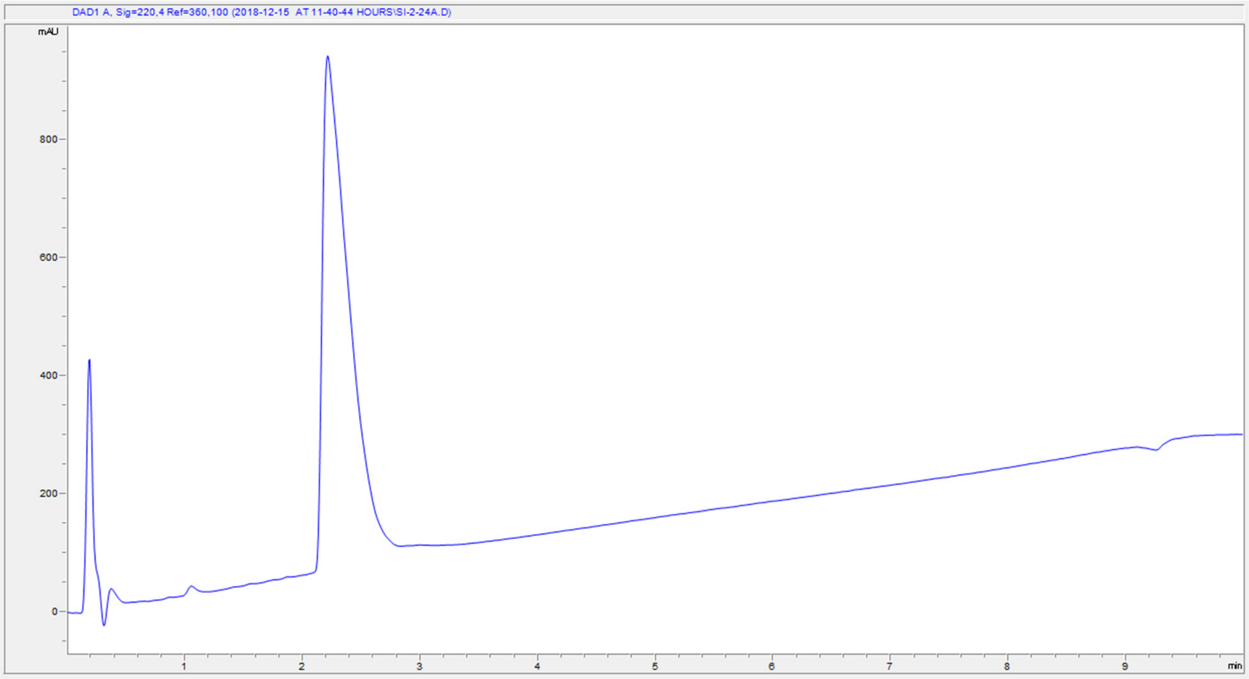
DAD1 A, Sig=220,4 Ref=360,100 (2018-12-15 AT 11-40-44 HOURS\SI-2-24A.D)
mAU
800
600
400
200
0
1
2
3
4
5
6
7
8
9
min

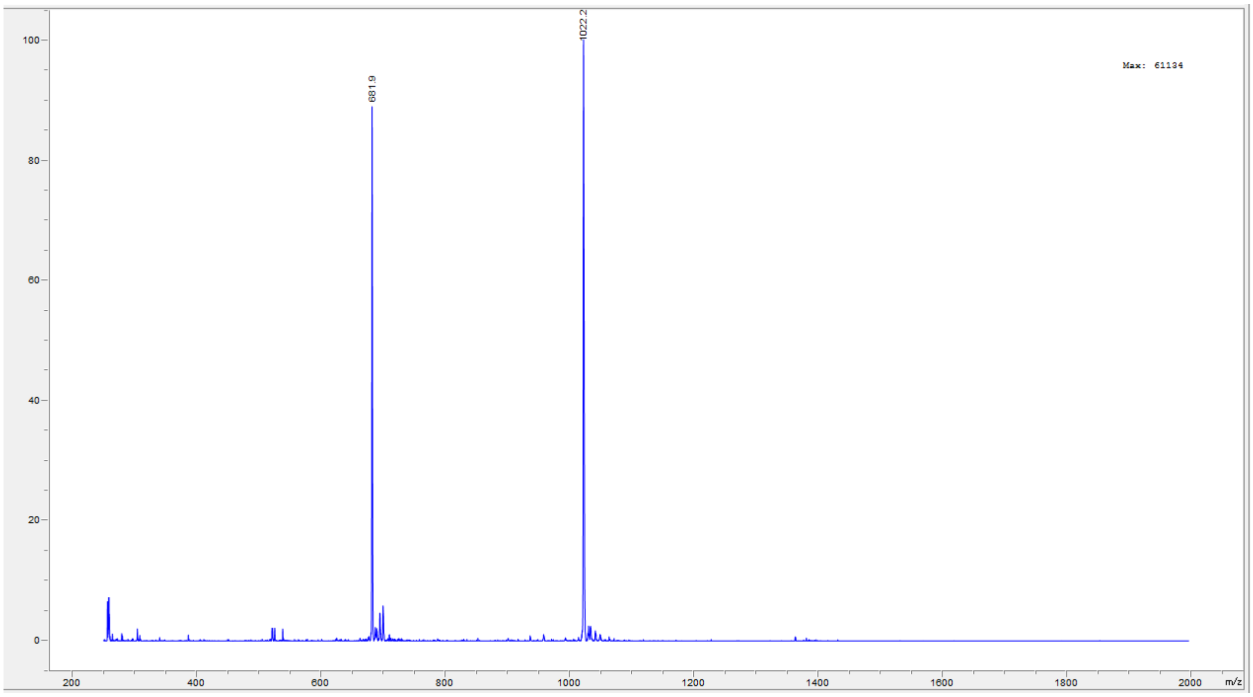
1022.2
681.9
Max: 61134
100
80
60
40
20
0
200
400
600
800
1000
1200
1400
1600
1800
2000
m/z

Peptide **33**.

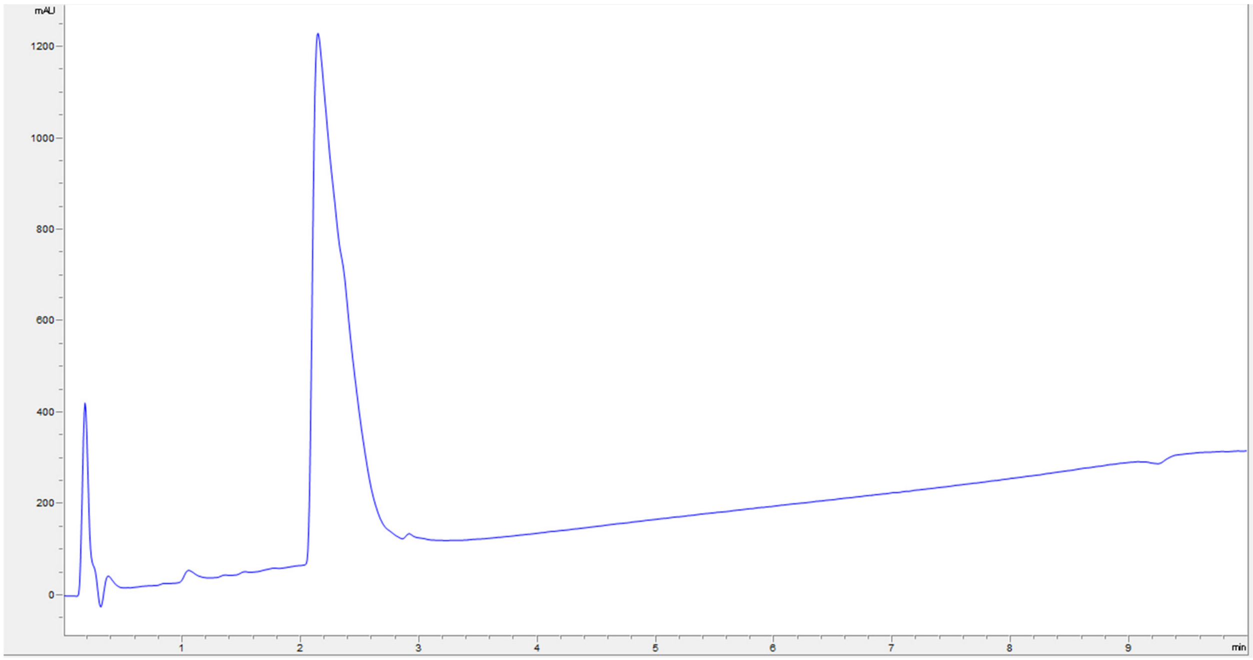

mAU
1200
1000
800
600
400
200
0
1
2
3
4
5
6
7
8
9
min


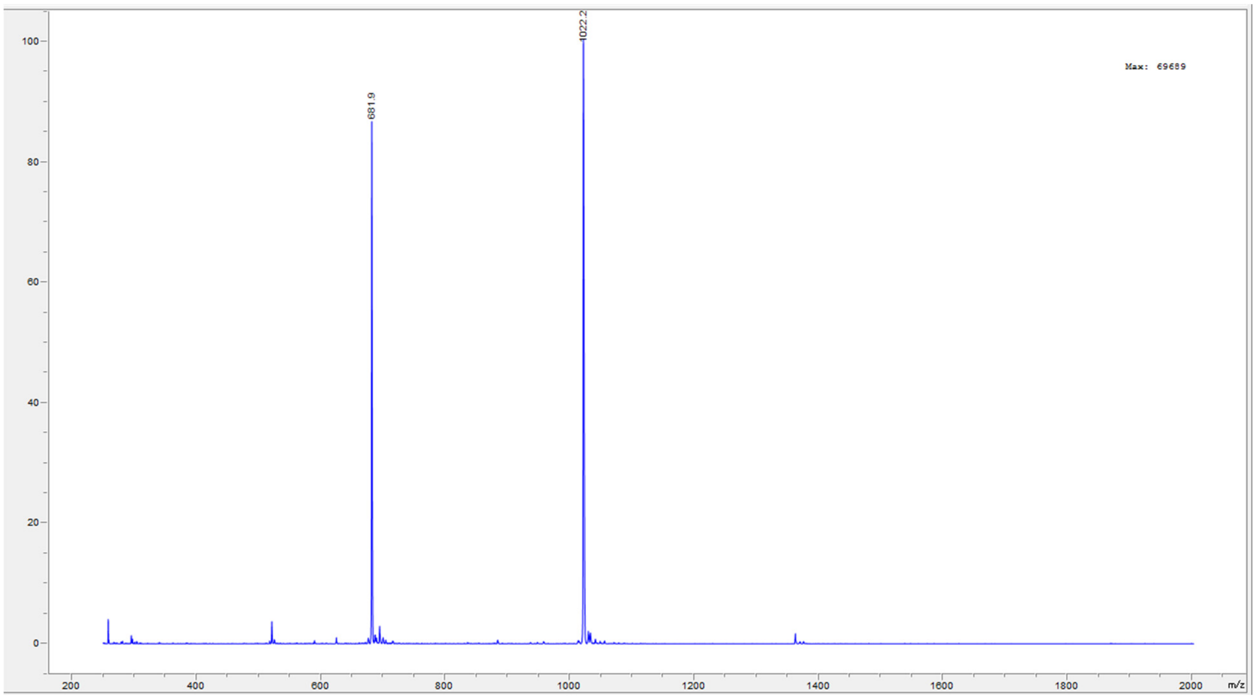

100
80
60
40
20
0
1022.2
681.9
Max: 69689
200
400
600
800
1000
1200
1400
1600
1800
2000
m/z

Peptide **35**.

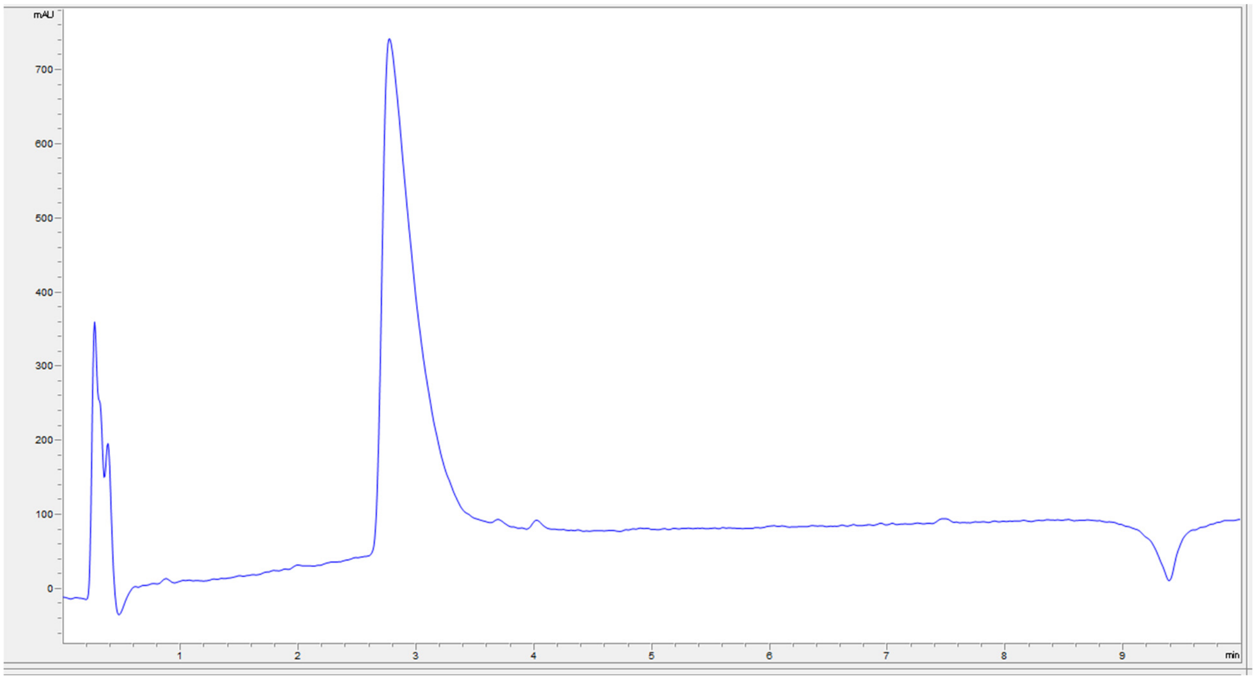
mAU
700
600
500
400
300
200
100
0
1
2
3
4
5
6
7
8
9
min

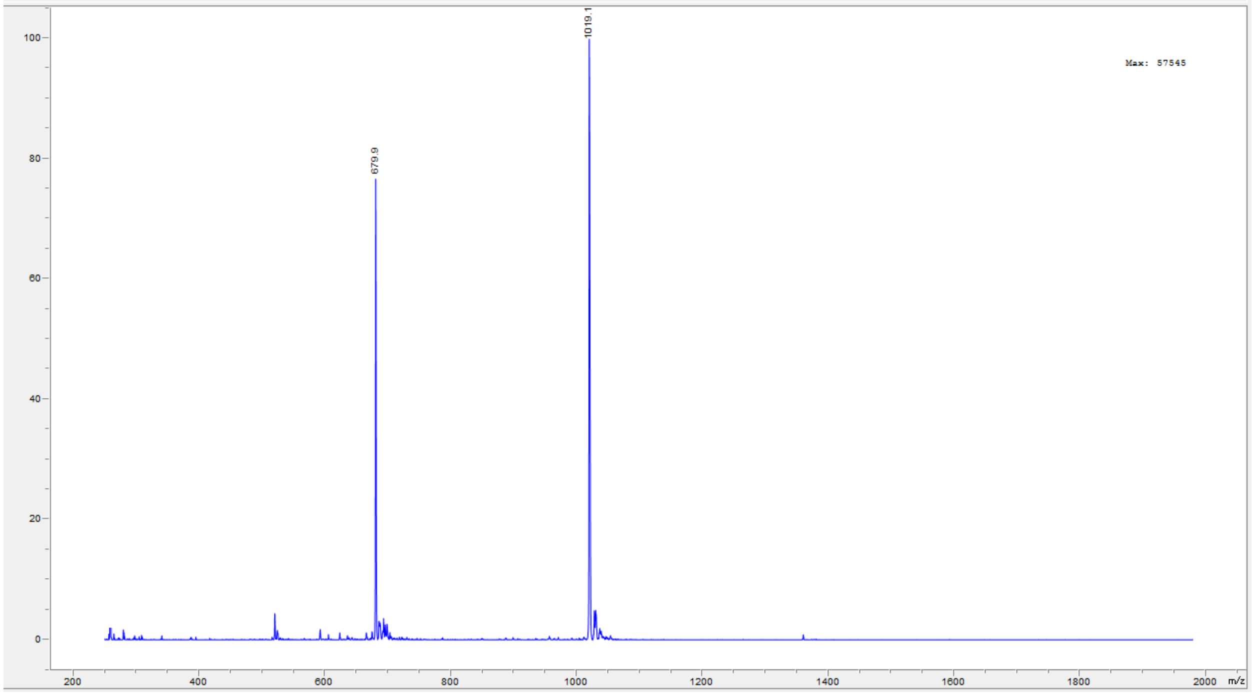
1019.1
679.9
Max: 57545
100
80
60
40
20
0
200
400
600
800
1000
1200
1400
1600
1800
2000
m/z

Peptide **36**.

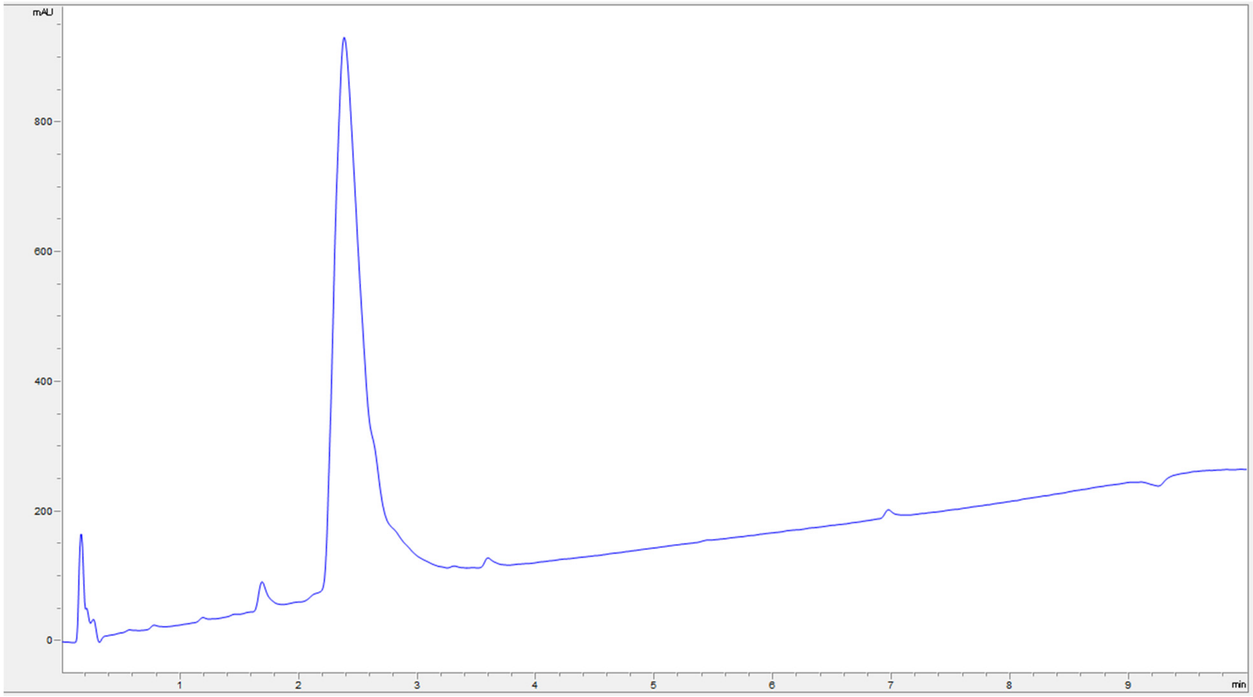
mAU
800
600
400
200
0
1
2
3
4
5
6
7
8
9
min

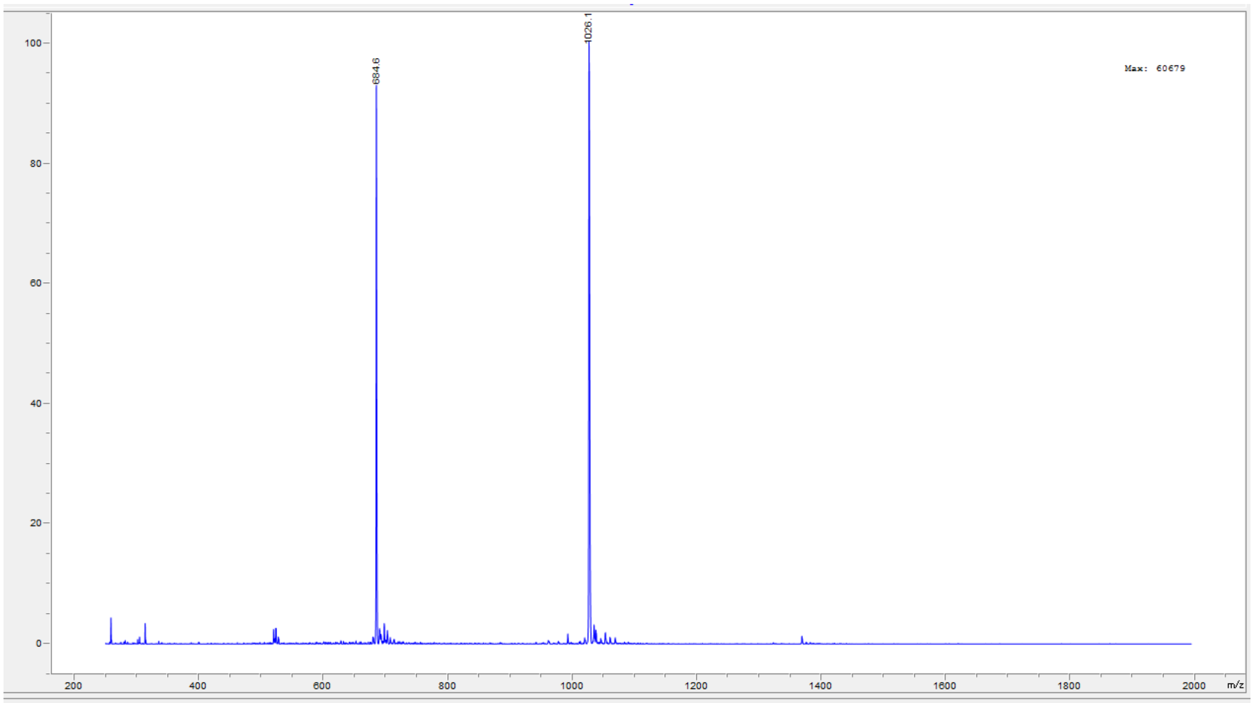
100
80
60
40
20
0
684.6
1026.1
Max: 60679
200
400
600
800
1000
1200
1400
1600
1800
2000
m/z

Peptide **37**.

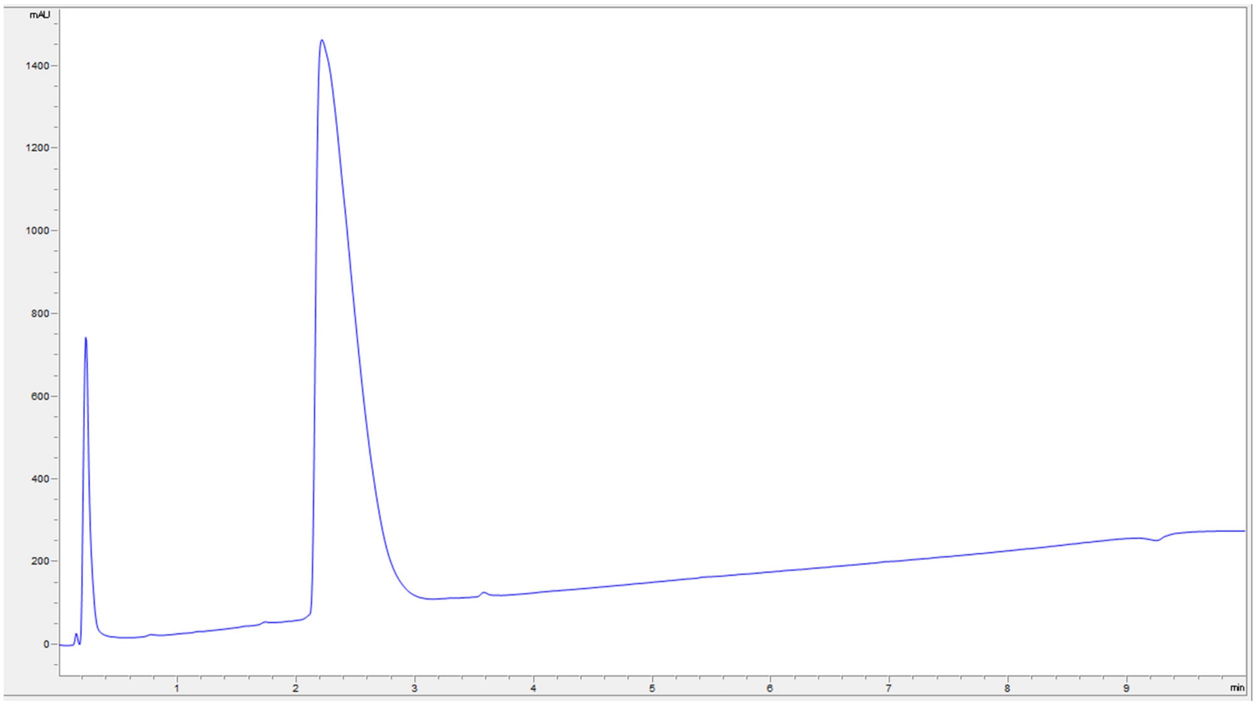

mAU
1400
1200
1000
800
600
400
200
0
1
2
3
4
5
6
7
8
9
min


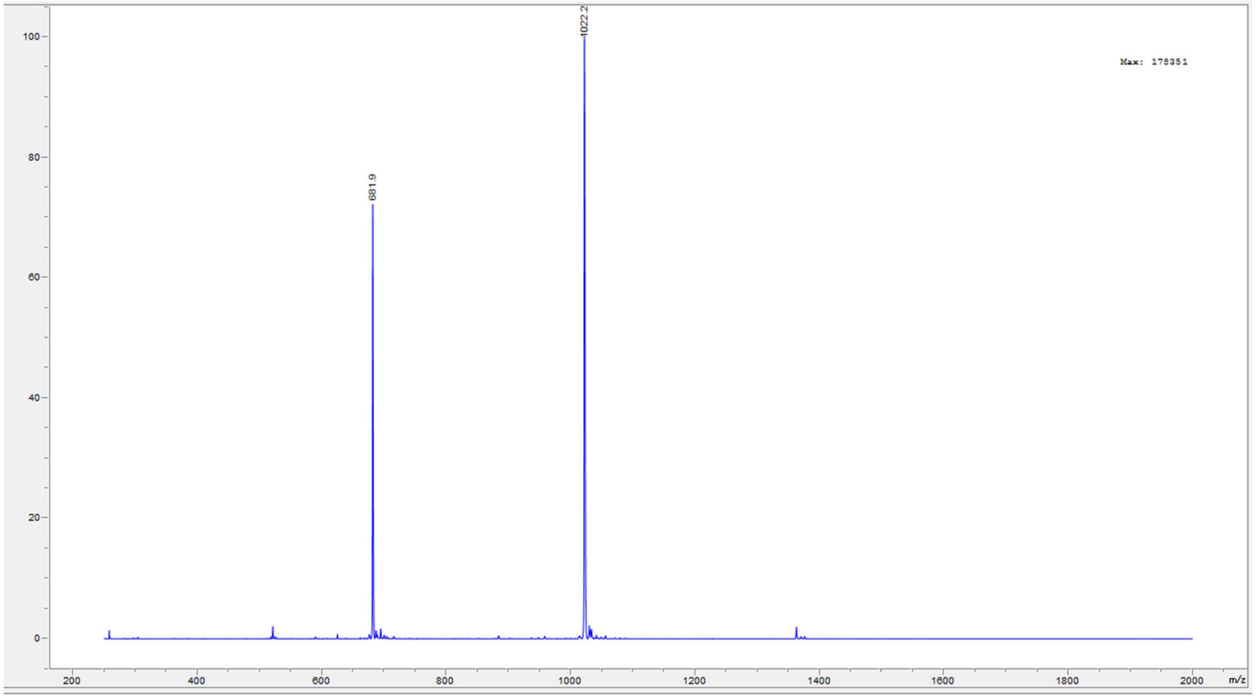

1022.2
681.9
Max: 178351
100
80
60
40
20
0
200
400
600
800
1000
1200
1400
1600
1800
2000
m/z

Peptide **39**.

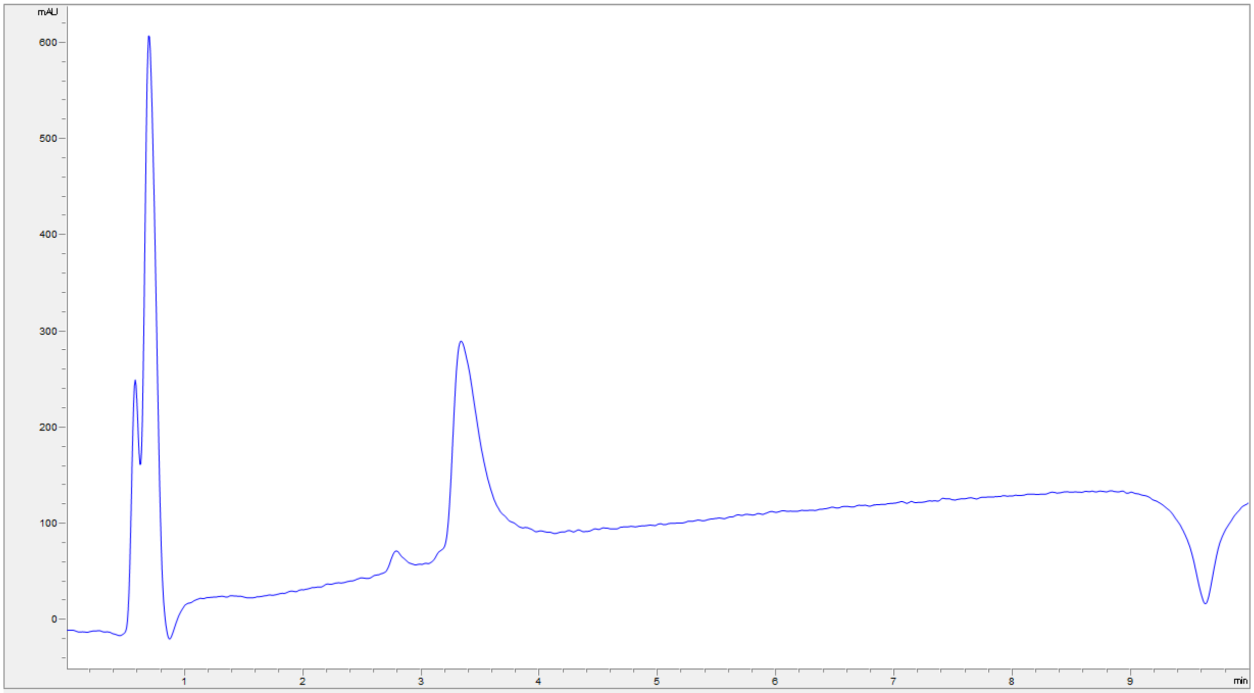
mAU
600
500
400
300
200
100
0
1
2
3
4
5
6
7
8
9
min

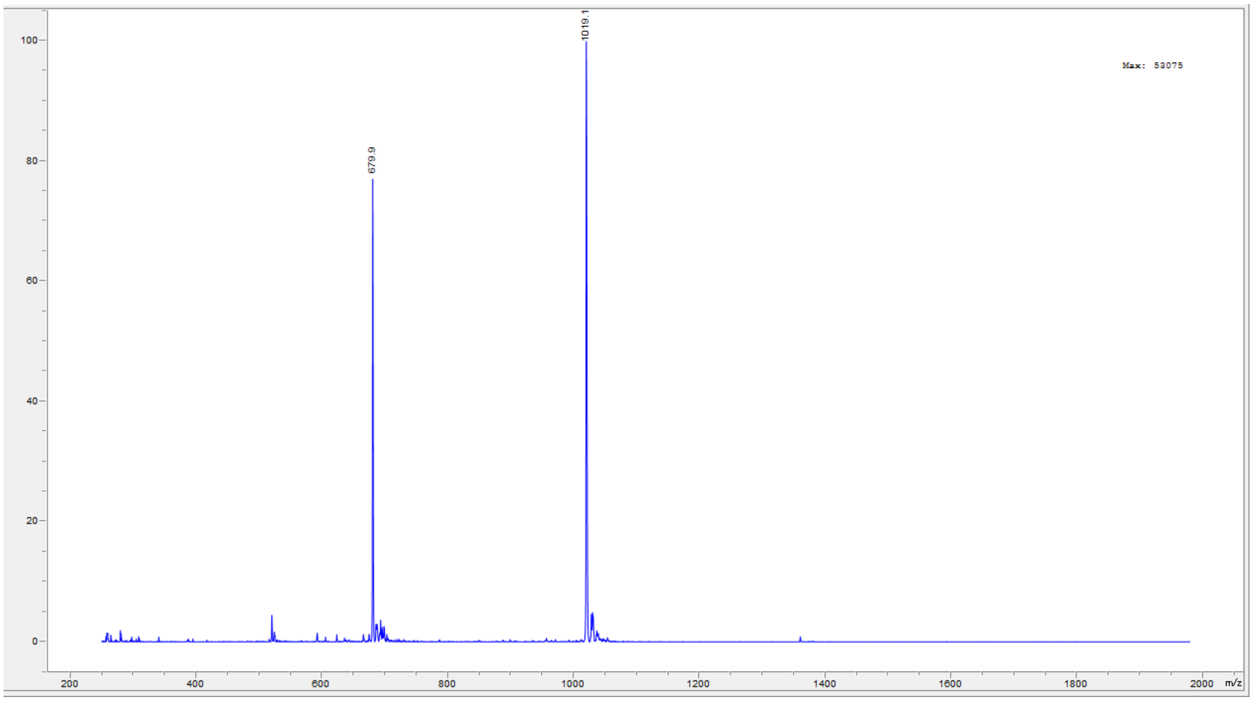
1019.1
679.9
Max: 53075
100
80
60
40
20
0
200
400
600
800
1000
1200
1400
1600
1800
2000
m/z

Peptide **40**.

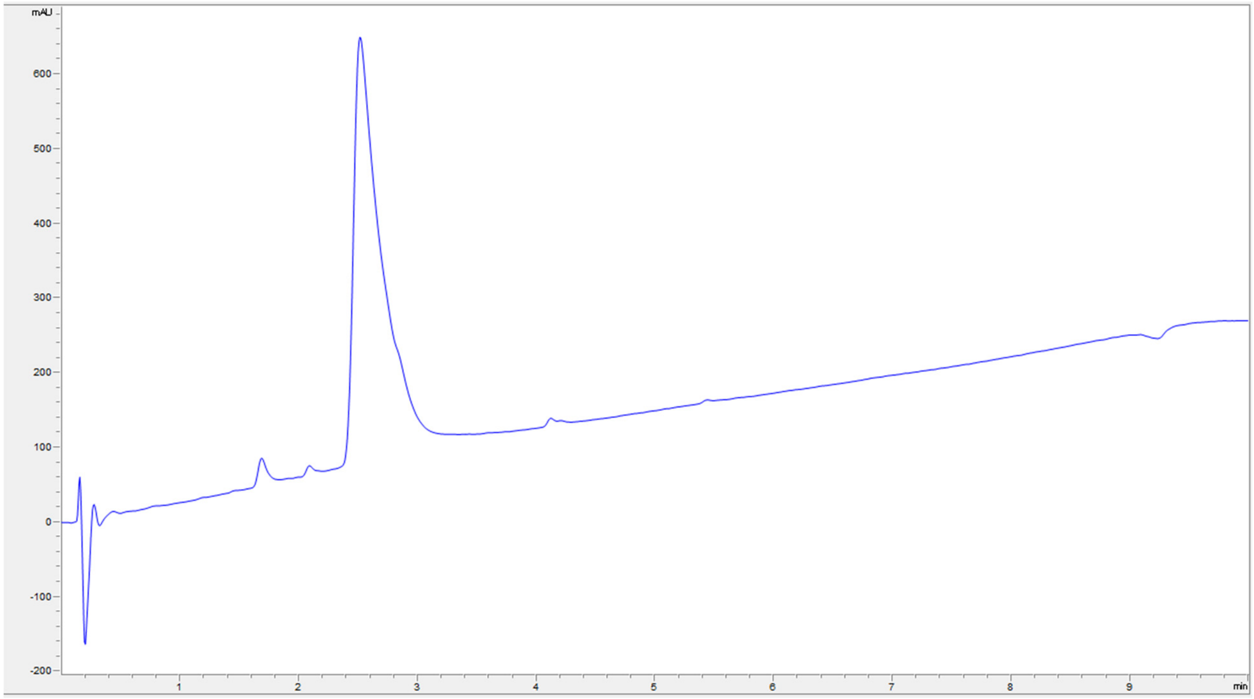
mAU
600
500
400
300
200
100
0
-100
-200
1
2
3
4
5
6
7
8
9
min

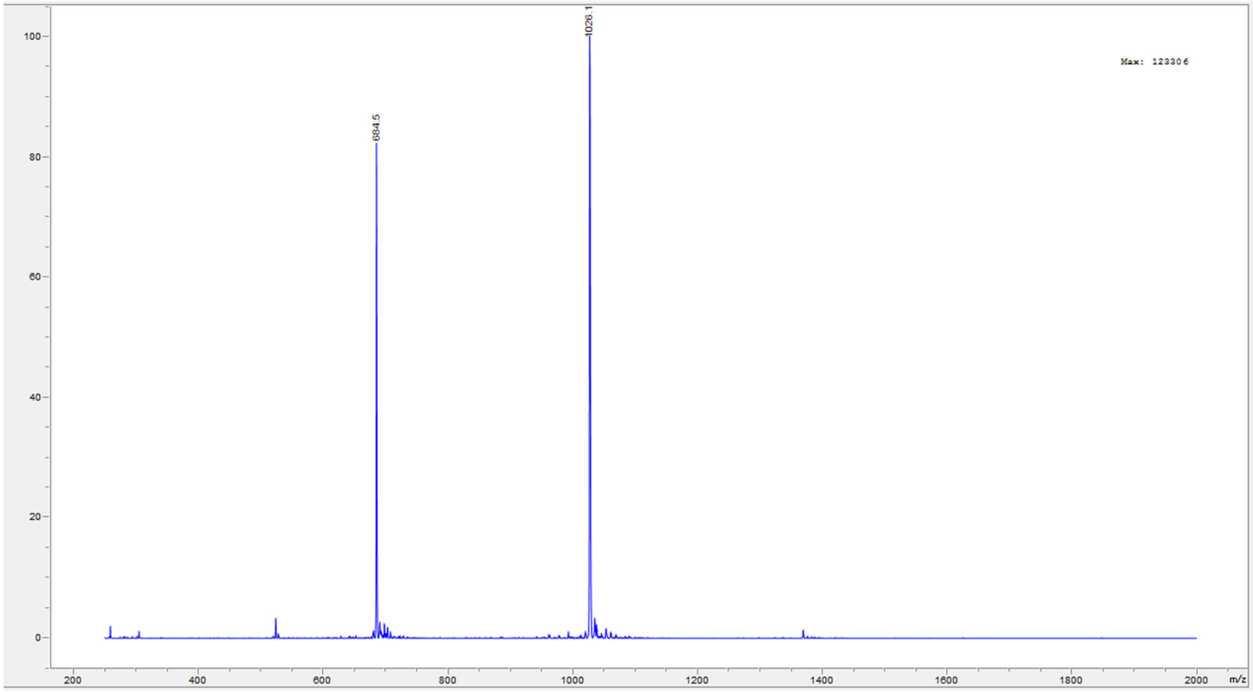
1026.1
684.5
Max: 123306
100
80
60
40
20
0
200
400
600
800
1000
1200
1400
1600
1800
2000
m/z

Peptide **41**.

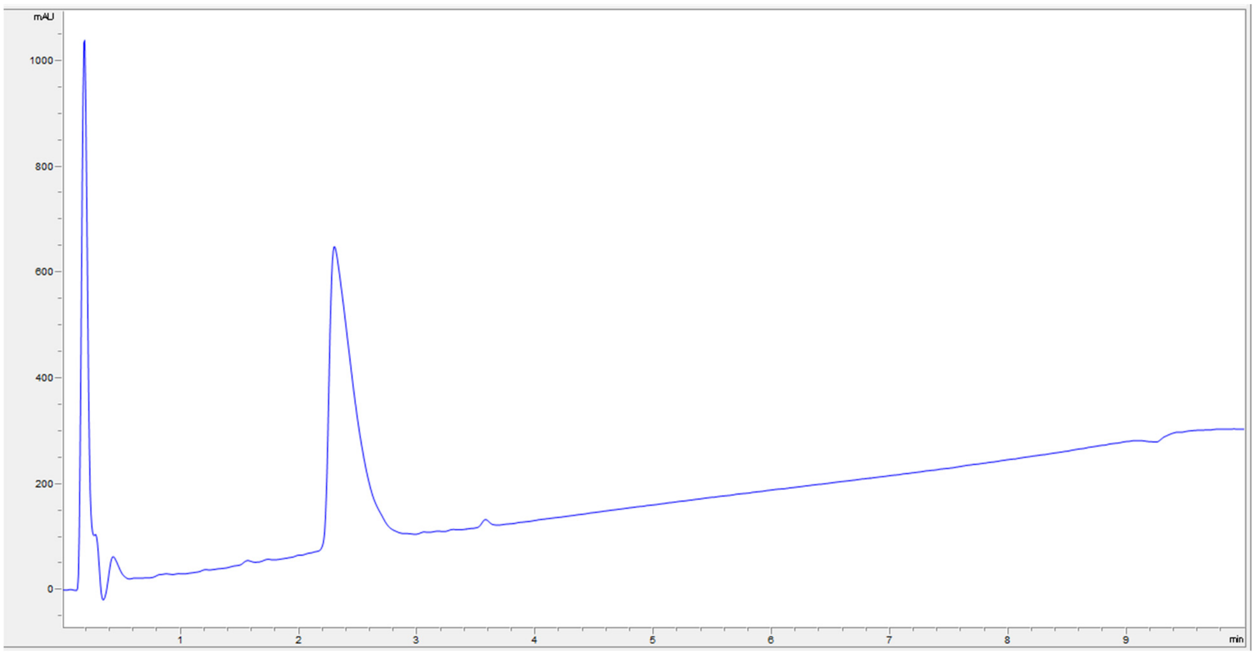
mAU
1000
800
600
400
200
0
1
2
3
4
5
6
7
8
9
min

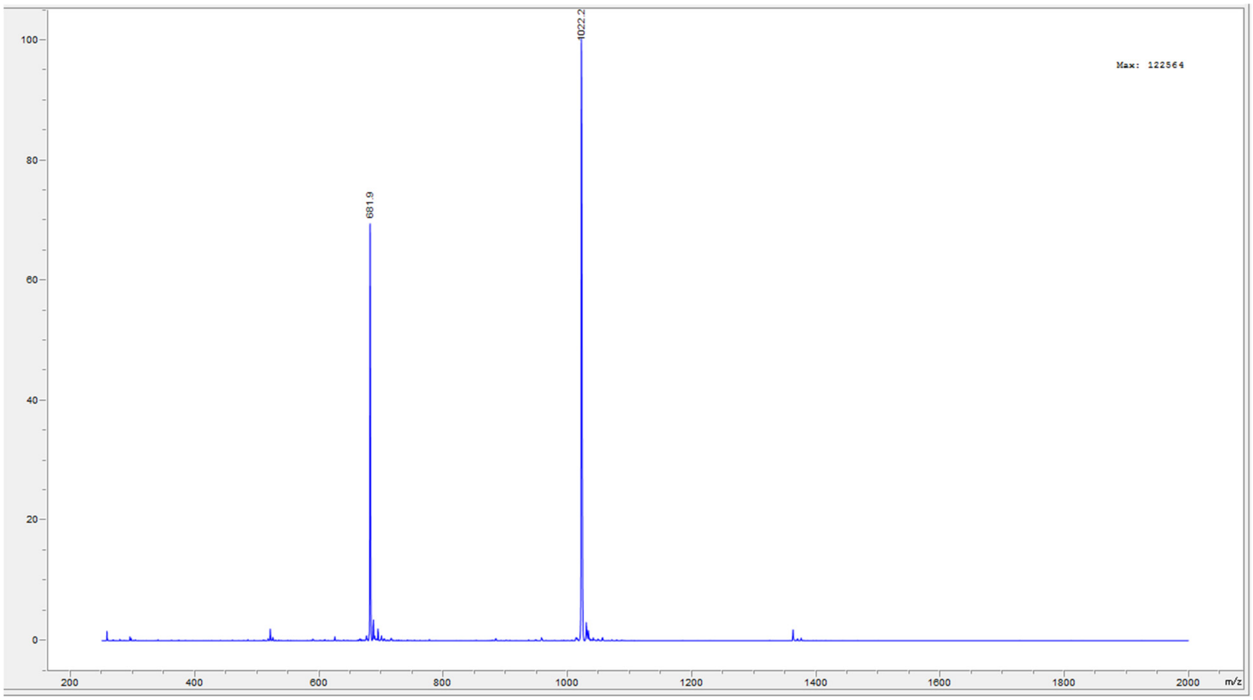
100
80
60
40
20
0
200
400
600
800
1000
1200
1400
1600
1800
2000
m/z
681.9
1022.2
Max: 122564

Peptide **42**.

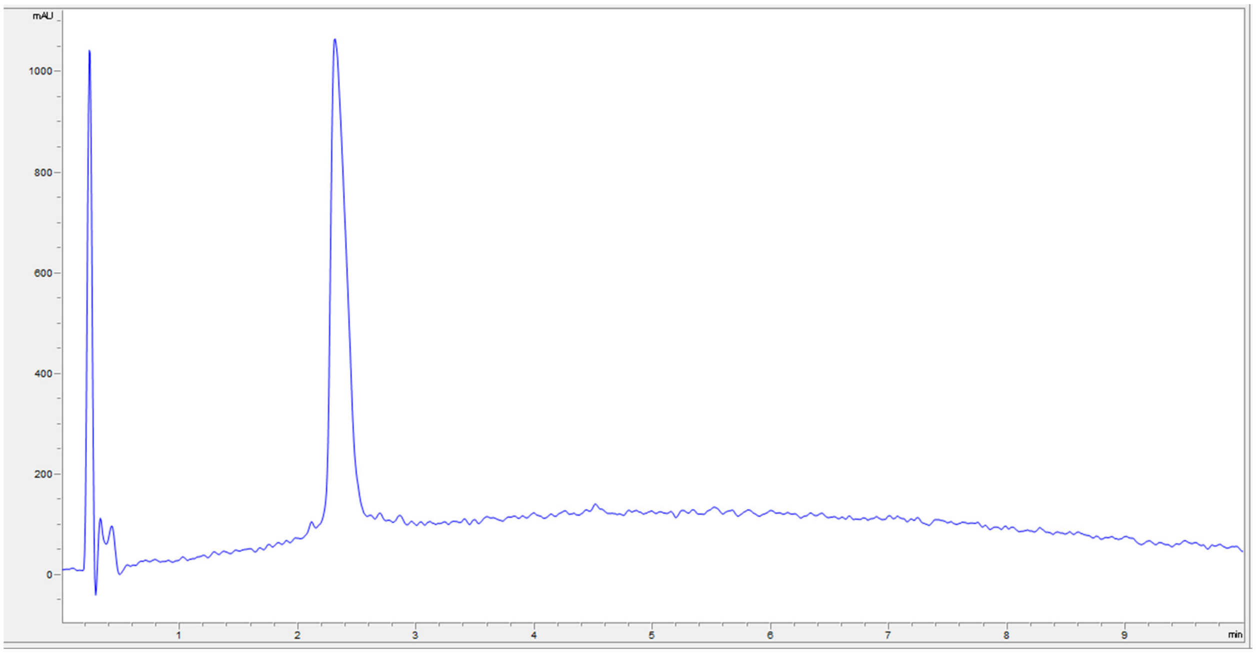
mAU
1000
800
600
400
200
0
1
2
3
4
5
6
7
8
9
min

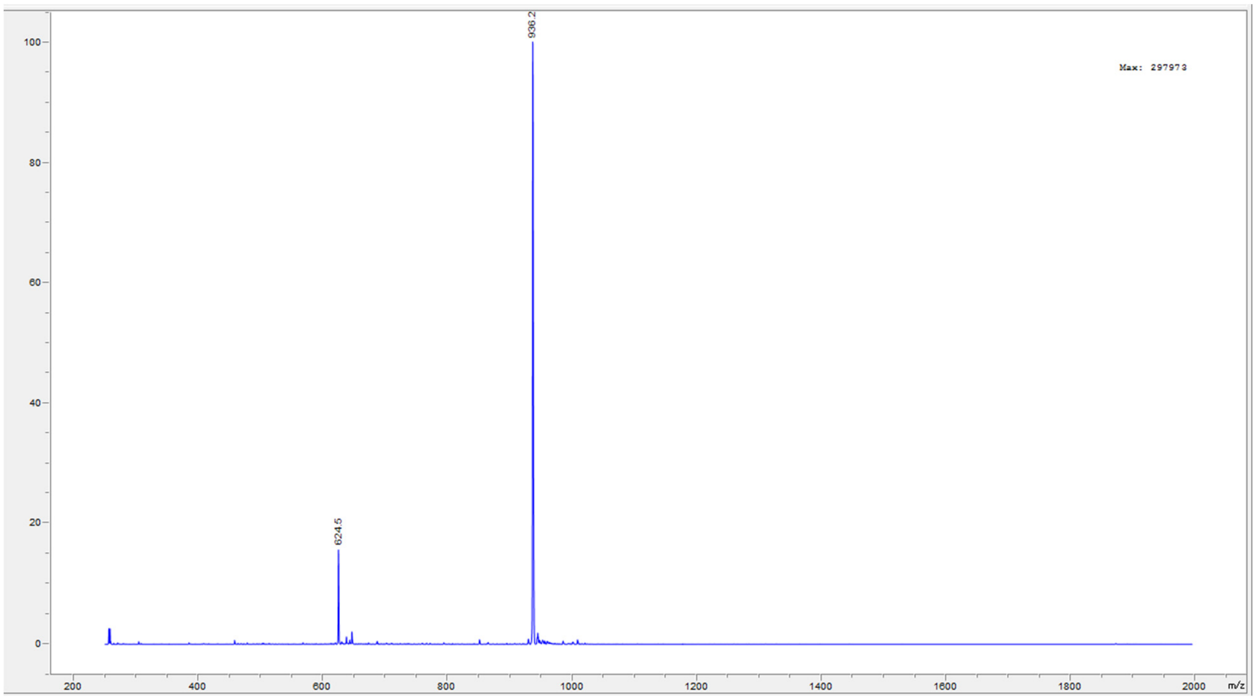
936.2
Max: 297973
624.5
100
80
60
40
20
0
200
400
600
800
1000
1200
1400
1600
1800
2000
m/z

Peptide **43**.

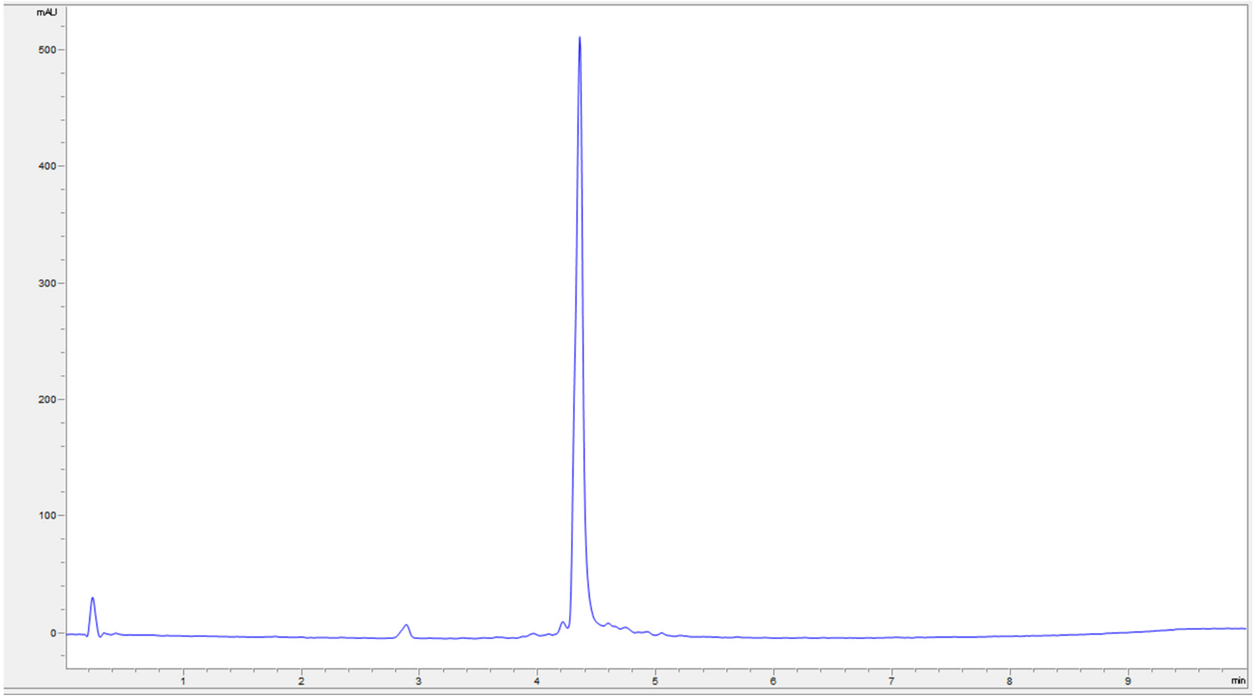
mAU
500
400
300
200
100
0
1
2
3
4
5
6
7
8
9
min

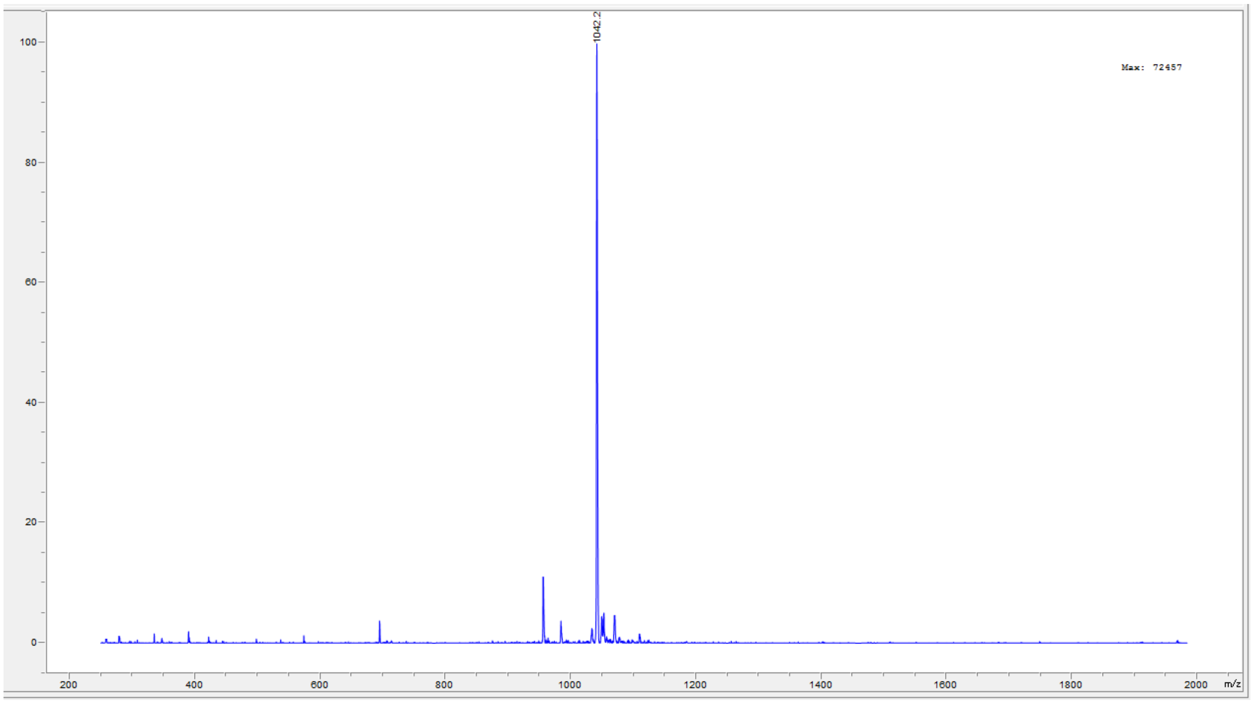
1042.2
Max: 72457
100
80
60
40
20
0
200
400
600
800
1000
1200
1400
1600
1800
2000
m/z

Peptide **44**.

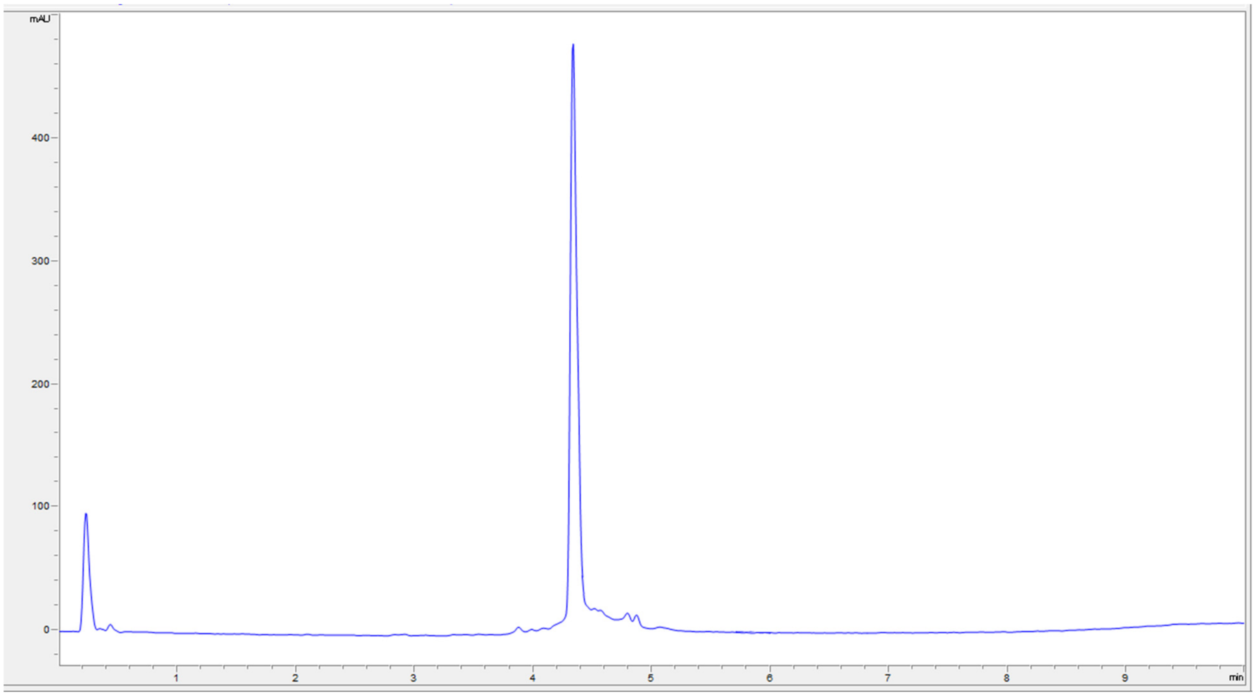
mAU
400
300
200
100
0
1
2
3
4
5
6
7
8
9
min

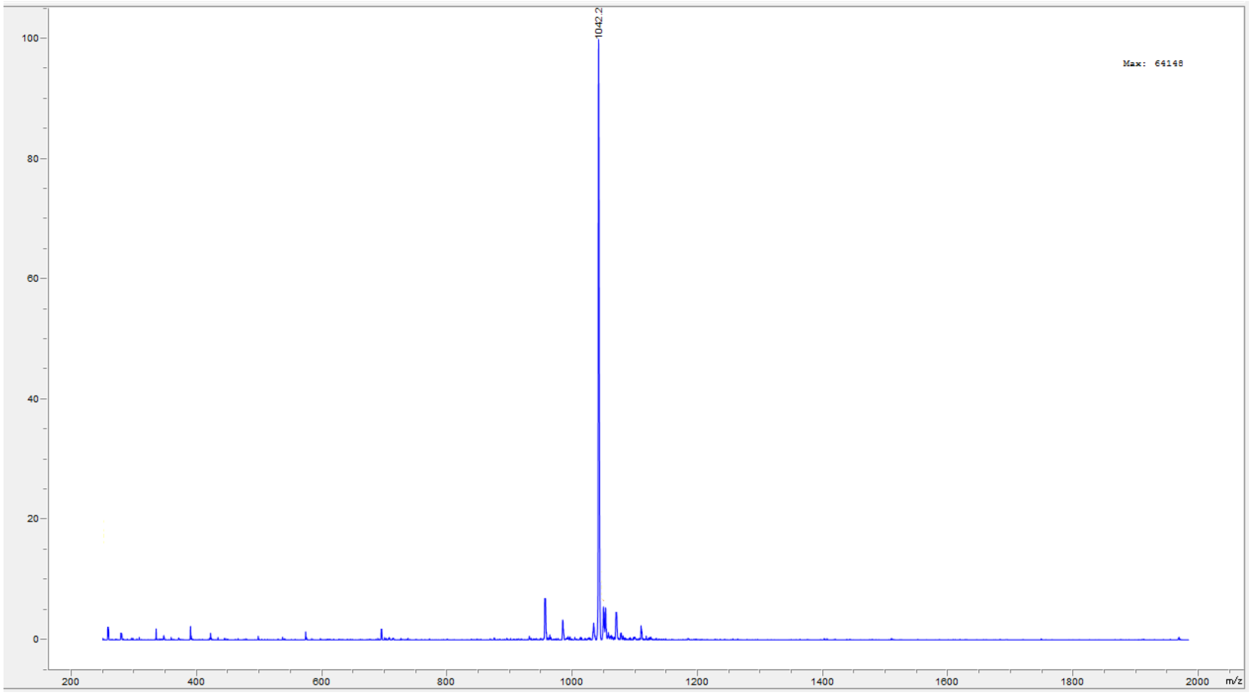
1042.2
Max: 64148
100
80
60
40
20
0
200
400
600
800
1000
1200
1400
1600
1800
2000
m/z

Peptide **45**.

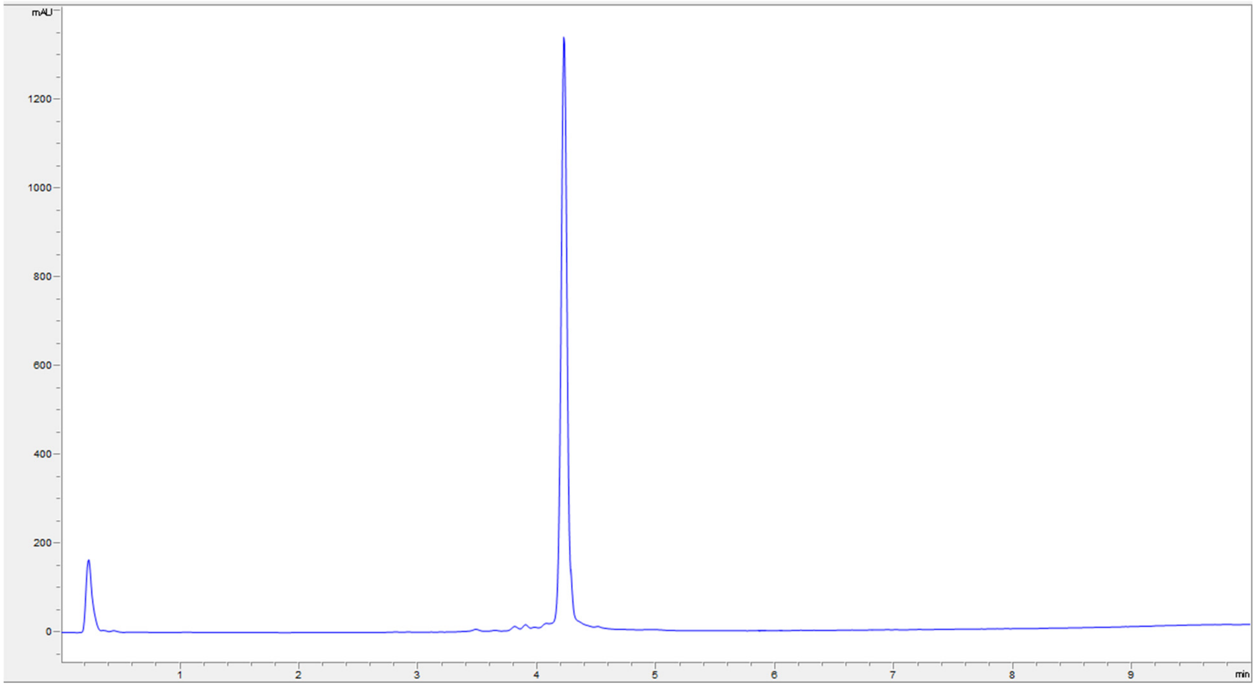

mAU
1200
1000
800
600
400
200
0
1
2
3
4
5
6
7
8
9
min


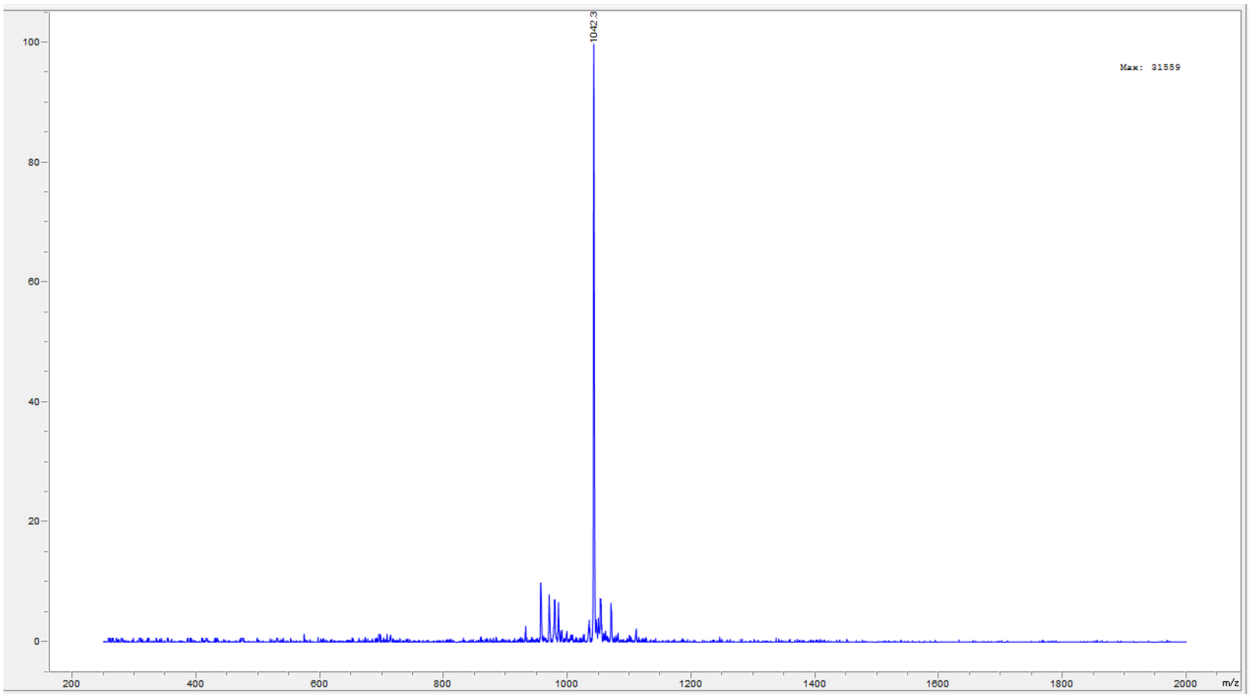

1042.3
Max: 31559
100
80
60
40
20
0
200
400
600
800
1000
1200
1400
1600
1800
2000
m/z

Peptide **46**.

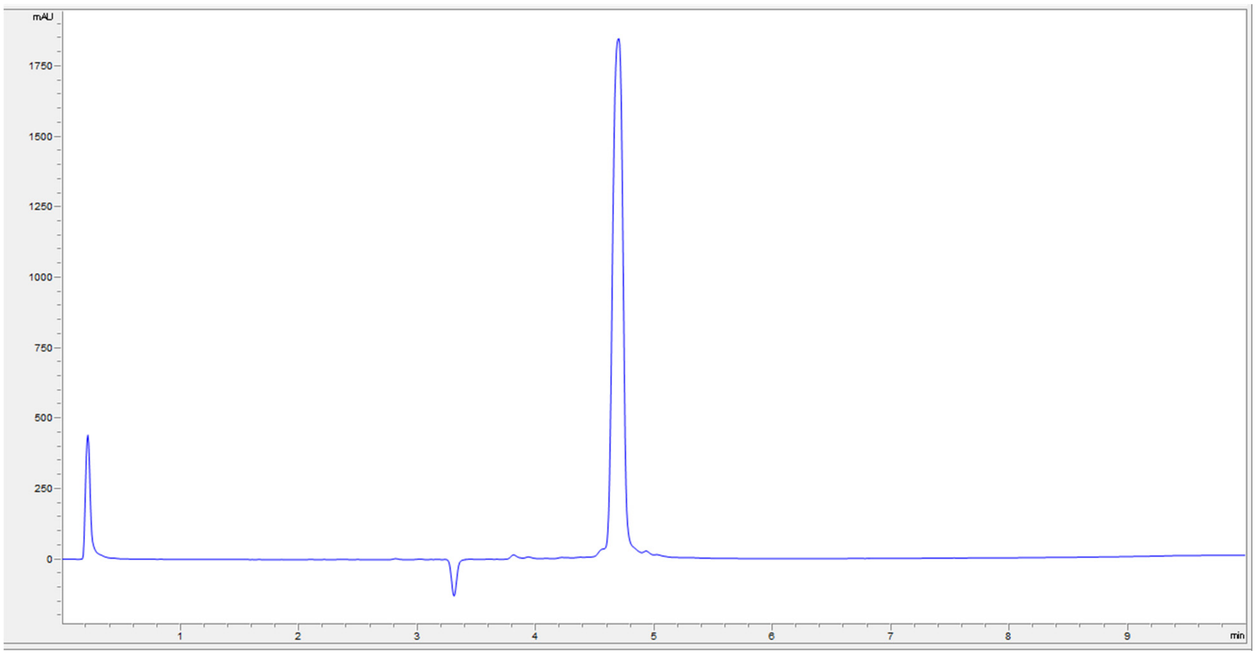

mAU
1750
1500
1250
1000
750
500
250
0
1
2
3
4
5
6
7
8
9
min


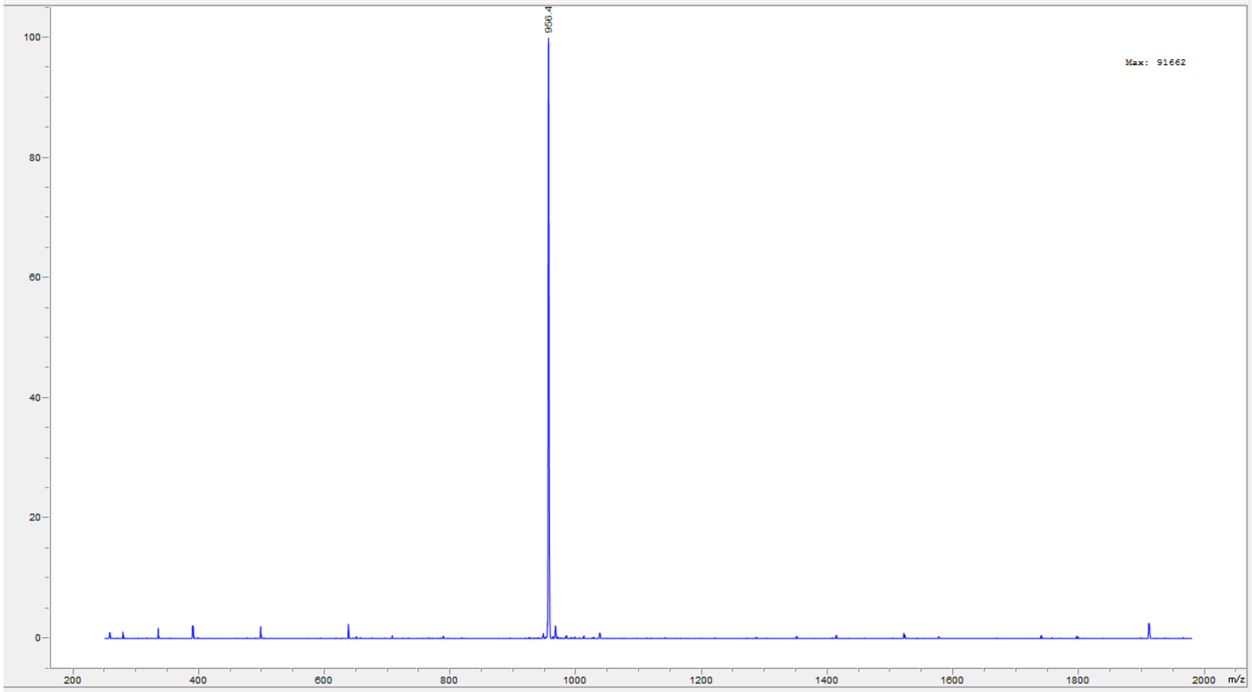

956.4
Max: 91662
100
80
60
40
20
0
200
400
600
800
1000
1200
1400
1600
1800
2000
m/z

Peptide **48**.

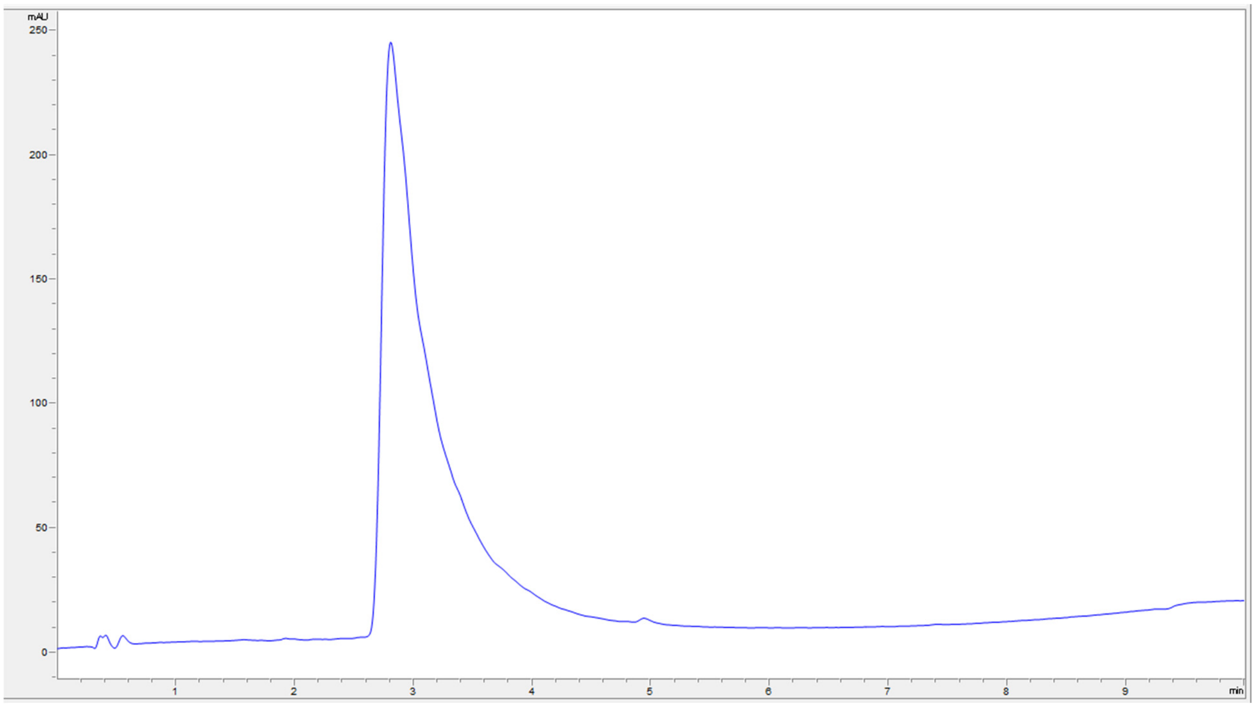

mAU
250
200
150
100
50
0
1
2
3
4
5
6
7
8
9
min


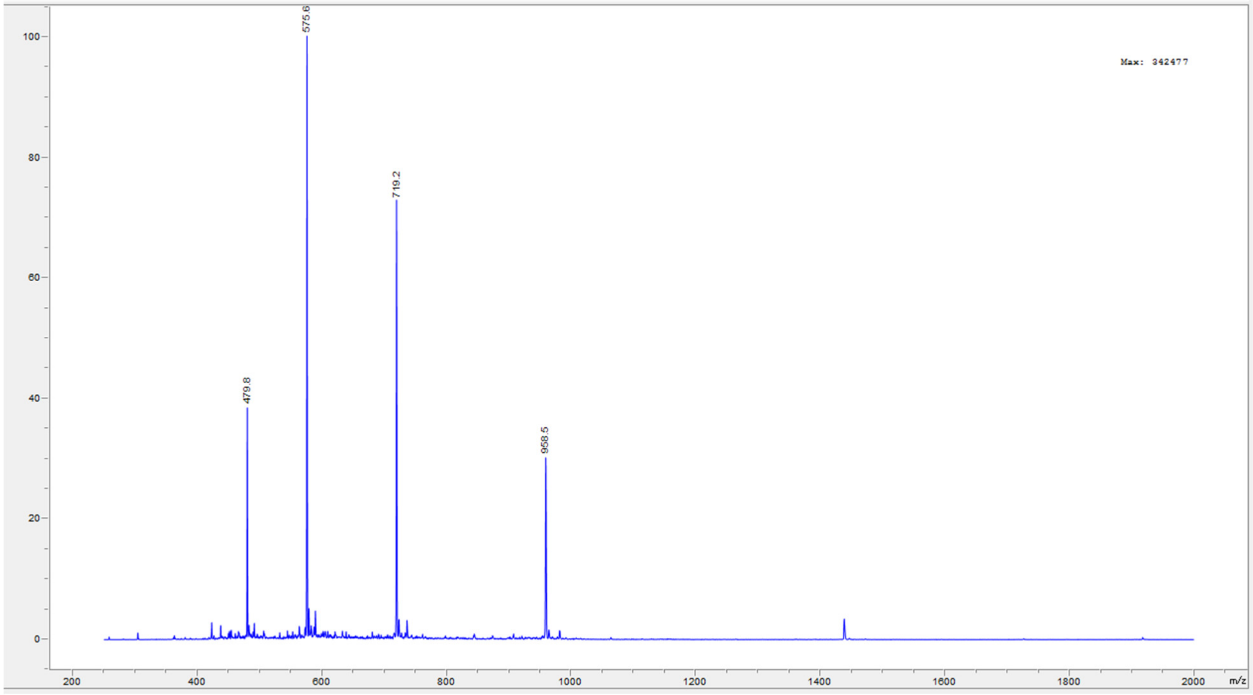

100
80
60
40
20
0
575.6
719.2
479.8
958.5
Max: 342477
200
400
600
800
1000
1200
1400
1600
1800
2000
m/z

Peptide **50**.

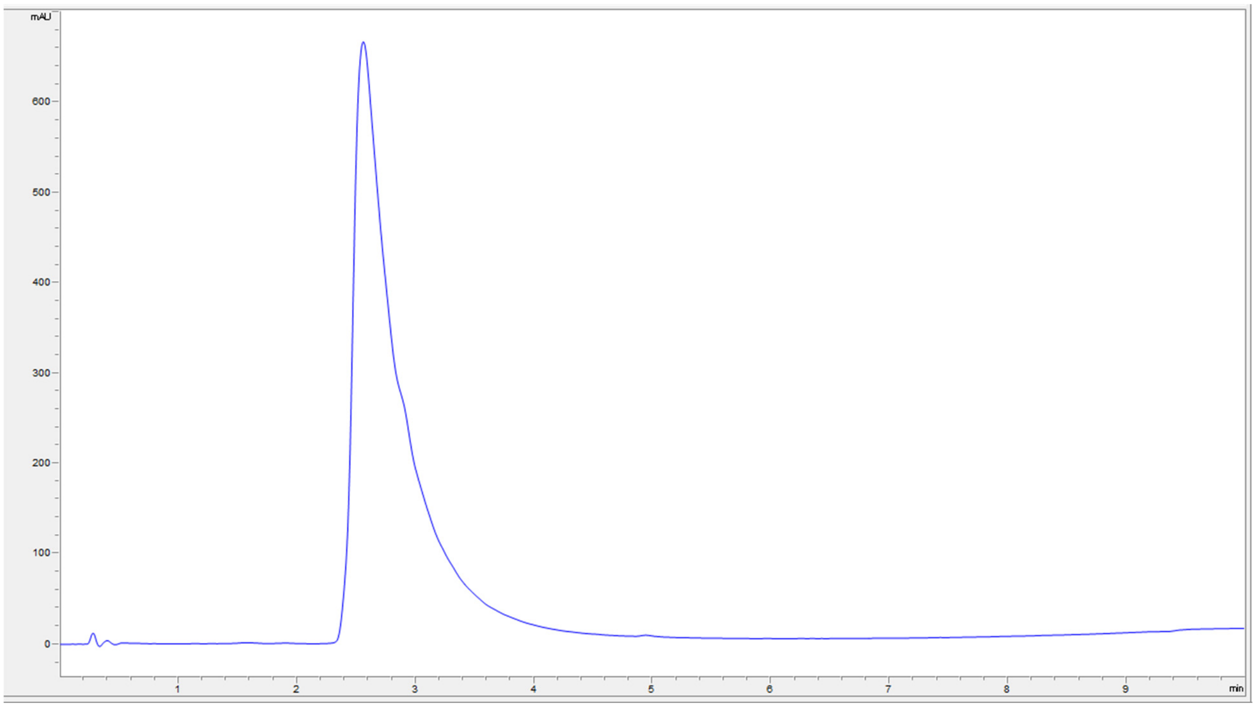
mAU
600
500
400
300
200
100
0
1
2
3
4
5
6
7
8
9
min

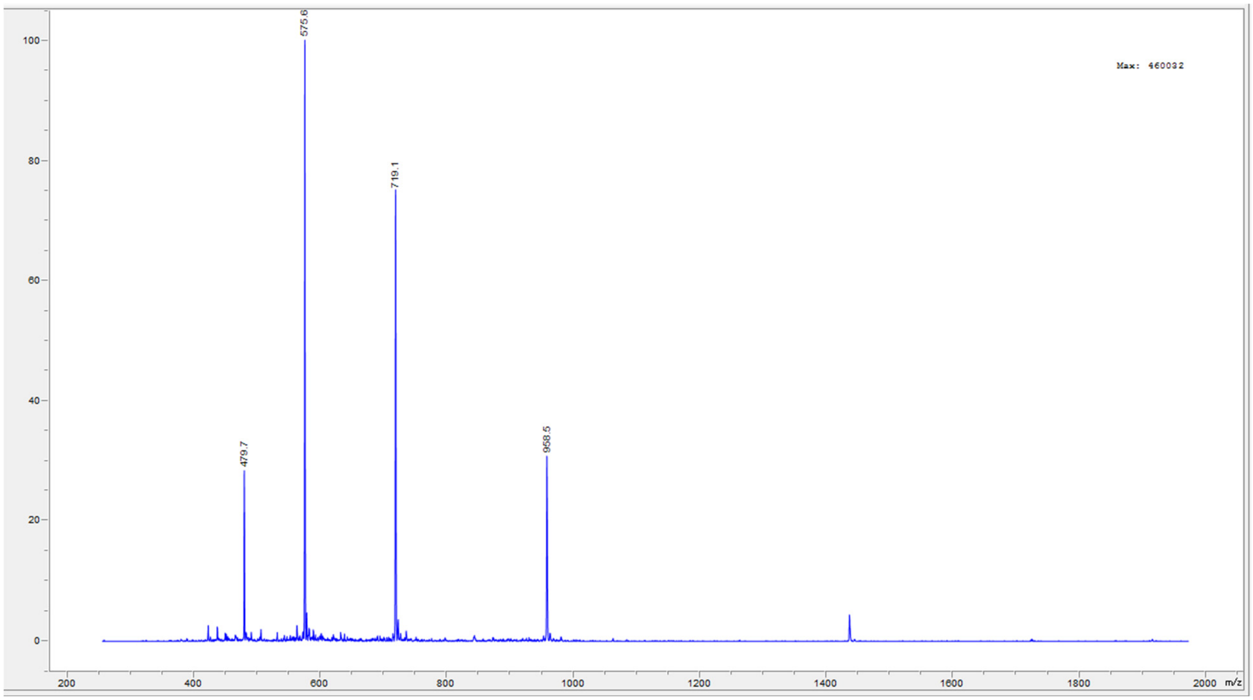
Max: 460032
575.6
719.1
479.7
958.5
100
80
60
40
20
0
200
400
600
800
1000
1200
1400
1600
1800
2000
m/z

Peptide **51**.

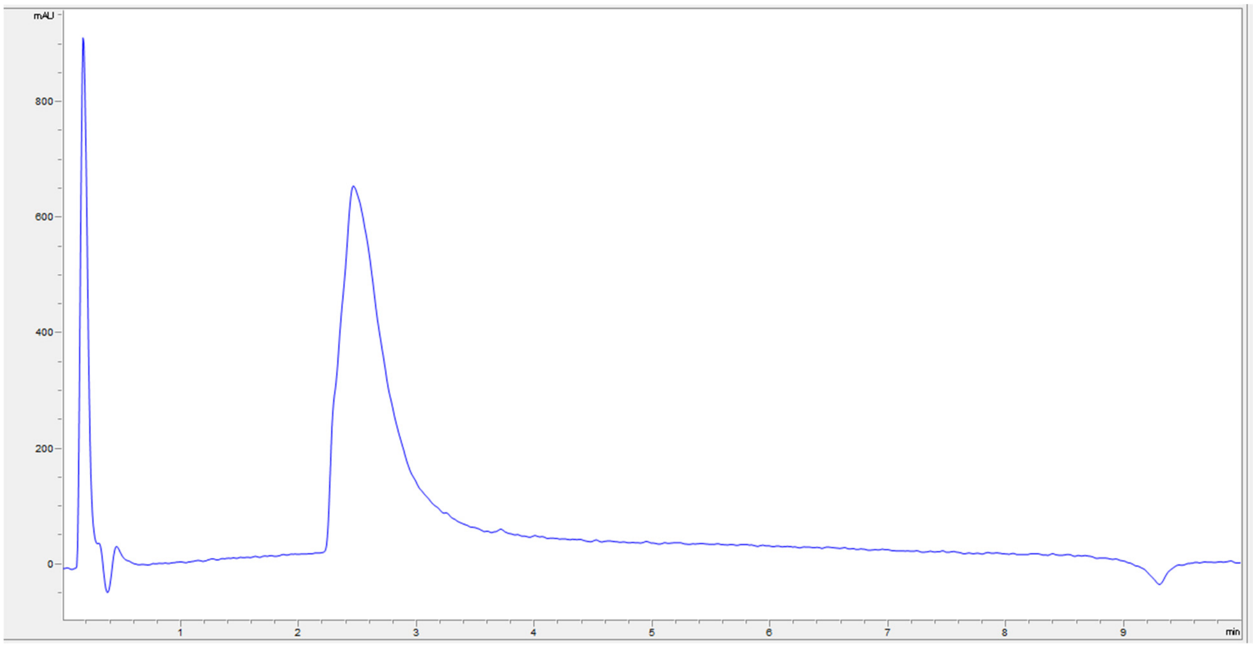
mAU
800
600
400
200
0
1
2
3
4
5
6
7
8
9
min

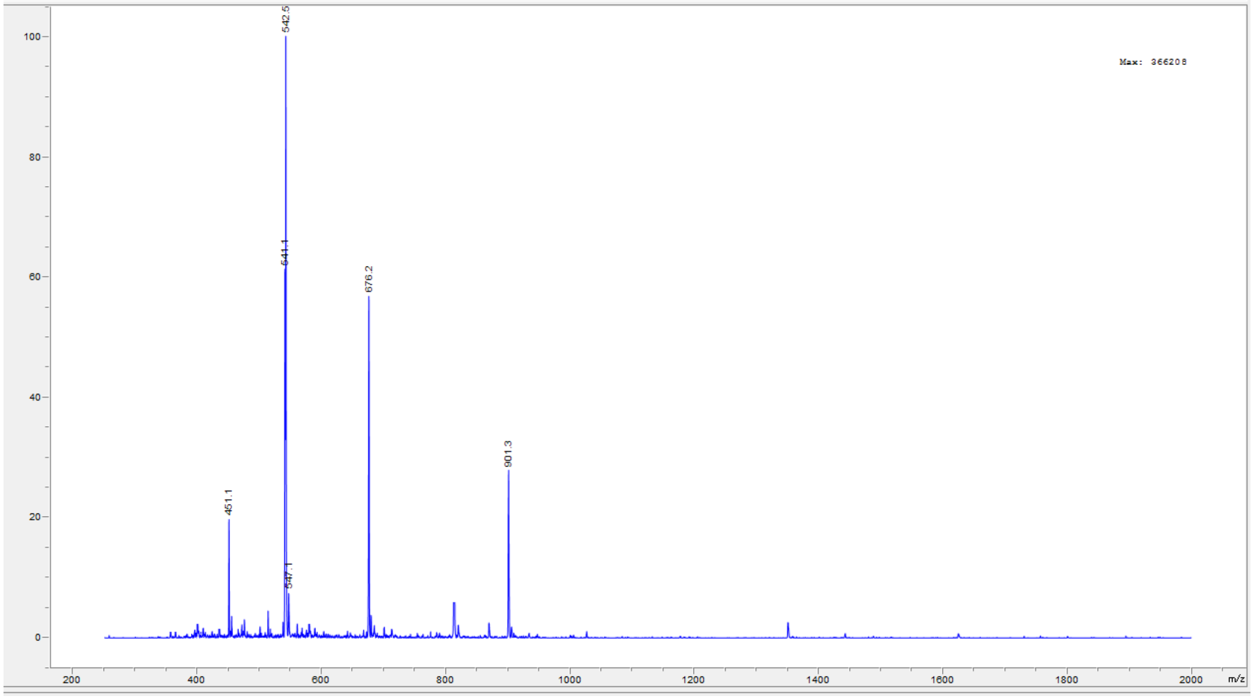
Max: 366208
542.5
541.1
676.2
901.3
451.1
547.1
100
80
60
40
20
0
200
400
600
800
1000
1200
1400
1600
1800
2000
m/z
